# A post-transcriptional regulatory checkpoint controls the response of tumor-infiltrating cytotoxic CD4^+^ T cells to immunotherapy

**DOI:** 10.1101/2025.09.09.675154

**Authors:** María Vila de Mucha, Julia Hłond, Cristobal Costoya, Ching-In Lau, Anna Śledzińska, Imran Uddin, Mariela Navarrete, Mansi Shah, Gerasimos Mastrokalos, Despoina Karagianni, Sarah E. Bell, Callum Nattress, Gordon Beattie, Christopher J. Tape, Martin Turner, Sergio A. Quezada, Richard G. Jenner

## Abstract

Acquisition of cytotoxic activity in CD4^+^ T cells (T_CTX_) can promote potent anti-tumor activity thus holding promise as a therapeutic approach. However, how this activity is regulated remains poorly understood. Here, we demonstrate that tumor-infiltrating CD4^+^ T_CTX_ activity is restrained by a post-transcriptional regulatory checkpoint. In untreated tumors, CD4^+^ T_CTX_ exist in a poised state, characterized by abundant *Gzmb* mRNA but limited Granzyme B (GzmB) protein. Differentiation into poised T_CTX_ is regulated by the Blimp-1-Bcl6 axis and requires type-I interferon signaling. Treatment with anti-CTLA-4 or anti-LAG-3 plus anti-PD-1 removed the block to GzmB protein production by repressing expression of the post-transcriptional regulator *Zfp36l1*. Constitutive *Zfp36l1* expression abrogated the effects of anti-CTLA-4 while deletion of *Zfp36l1* and its paralog *Zfp36* triggered GzmB protein production and promoted tumor control. These data identify ZFP36/ZFP36L1 as a key post-transcriptional regulatory checkpoint of CD4^+^ T_CTX_ activity and a potential immunotherapy target in cancer.

## Main

Immunotherapy has revolutionized cancer treatment, but responses remain limited to a subset of patients. Studies of the contribution of CD4^+^ T cells to tumor immunity primarily focus on their helper activity. However, tumor-infiltrating CD4^+^ T cells can also acquire direct cytotoxic activity that controls tumor growth^1,2^. These cells hold promise as a therapeutic modality but how their activity is regulated remains poorly understood.

Cytotoxic CD4^+^ T cells (T_CTX_) are marked by production of cytolytic molecules such as Granzyme B (GzmB) and Perforin^3,4^. During viral infection in mouse and humans, CD4^+^ T_CTX_ are induced in response to inflammatory cytokines such as IL-2 and IFNγ and can directly kill infected cells^5^. CD4^+^ T_CTX_ have also been described in murine tumor models, where both adoptively transferred and endogenous CD4^+^ T cells can acquire direct, MHC class II-restricted anti-tumor cytotoxic activity^3,4,6,7^. Similarly, CD4^+^ T_CTX_ that directly lyse tumor cells *ex vivo* have been identified in patients with both solid and hematological malignancies^8,9,10,11,12^. Single-cell RNA sequencing (scRNA-seq) studies have also shown that CD4^+^ T cells with high levels of *GZMB* mRNA are commonly found in colorectal cancer (CRC), liver cancer, bladder cancer, non-small cell lung cancer (NSCLC), and basal cell carcinoma (BCC) tumor microenvironments^13,14,15,16,17^.

Despite being identified in many cancer types, we have limited knowledge of the factors that regulate tumor-infiltrating CD4^+^ T_CTX_ differentiation and function. Tumor-reactive CD4^+^ T_CTX_ are enriched in patients with advanced melanoma after treatment with the anti-CTLA-4 targeting antibody, ipilimumab^8^, suggesting that acquisition of a cytotoxic phenotype in CD4^+^ T cells can be induced by immunomodulatory agents. Similarly in murine tumor models, treatment with anti-CTLA-4 or other T_REG_ depleting therapies drives the appearance of CD4^+^ T_CTX_ and tumor regression in an IL-2 dependent manner^7,18^.

We have previously shown that CD4^+^ T_CTX_ differentiation in tumors is dependent on the transcriptional repressor Blimp-1^7^, consistent with observations from viral infection models^19,20,21^. Differentiation of CD4^+^ T_CTX_ in viral models occurs in direct opposition to the T_FH_ cell fate, which is promoted by the transcriptional repressor Bcl6^19^. Blimp-1 and Bcl6 also act in a mutually antagonistic manner to regulate cell fate decisions in other contexts^22,23^. IL-2 promotes Blimp-1 expression and inhibits Bcl6 expression^24,25,26^, suggesting that it may drive CD4^+^ T cell cytotoxicity in tumors through modulation of the Blimp-1-Bcl6 axis.

A sole focus on identifying transcriptional regulatory mechanisms that underlie tumor infiltrating lymphocyte (TIL) function risks overlooking potential roles for post-transcriptional processes. Indeed, it is becoming increasingly apparent that T cell function and homeostasis are also regulated at the level of mRNA stability and translation^27,28,29^. Production of effector molecules, including IFNγ and TNFα, are regulated at a post-transcriptional level in T cells in vitro and in the context of viral infections^30,31,32,33,34^, with similar regulation recently having been observed in CD8^+^ TILs^35,36,37,38^.

Here, we investigated the transcriptional and post-transcriptional factors that govern acquisition of a cytotoxic phenotype by tumor-infiltrating CD4^+^ T cells. We propose a two-step model in which acquisition of cytotoxic activity in CD4^+^ TILs requires activation of both transcriptional and post-transcriptional switches, which may have important implications for immunotherapy approaches aiming to boost the anti-tumor activity of CD4^+^ T_CTX_.

## Results

### Untreated tumors contain inactive but poised CD4^+^ T_CTX_

We sought to investigate the transcriptional profile of tumor-infiltrating CD4^+^ T_CTX_. To do this, we used the MCA205 fibrosarcoma model, in which we have previously shown that treatment with anti-CTLA-4 mouse IgG2a antibody (henceforth anti-CTLA-4)^39,40^ drives differentiation of CD4^+^ T cells into GzmB-producing T_CTX_ and complete tumor rejection in a manner dependent on depletion of intra-tumoral T_REG_^7^ (Extended Data Fig. 1a-c).

We performed scRNA-seq of CD4^+^ TILs from MCA205-bearing mice treated with anti-CTLA-4 on days 6 and 9 following tumor inoculation or left untreated. Mice were sacrificed on day 10 to ensure capture of responses during the initiation of tumor control (Fig. 1a). High dimensionality reduction and unsupervised clustering of CD4^+^ T cells from untreated and anti-CTLA-4 treated tumors identified 13 populations, which included three populations of *Foxp3*-expressing T_REG_ (clusters 0, 6 and 10), one proliferative population (cluster 5; high expression of *Mki67*) and nine T_EFF_ populations (Fig. 1a,b, Extended Data Fig. 1d and Supplementary Table 1). The T_EFF_ populations could be broadly grouped into cells with high expression of the activation markers *Tnfrsf9* (4-1BB) and *Pdcd1* (PD-1) and low expression of the early differentiation markers *Tcf7* and *Sell*, and cells with the opposite expression pattern (Fig. 1b). We also identified a population of activated T_EFF_ (cluster 7) that expressed high levels of the T_H_1-associated genes *Ifng*, *Csf2* and *Tnf*, as well as a distinct population of *Gzmb* expressing activated T_EFF_ (T_CTX_, cluster 1) (Fig. 1a,b and Extended Data Fig. 1d).

**Fig. 1:**
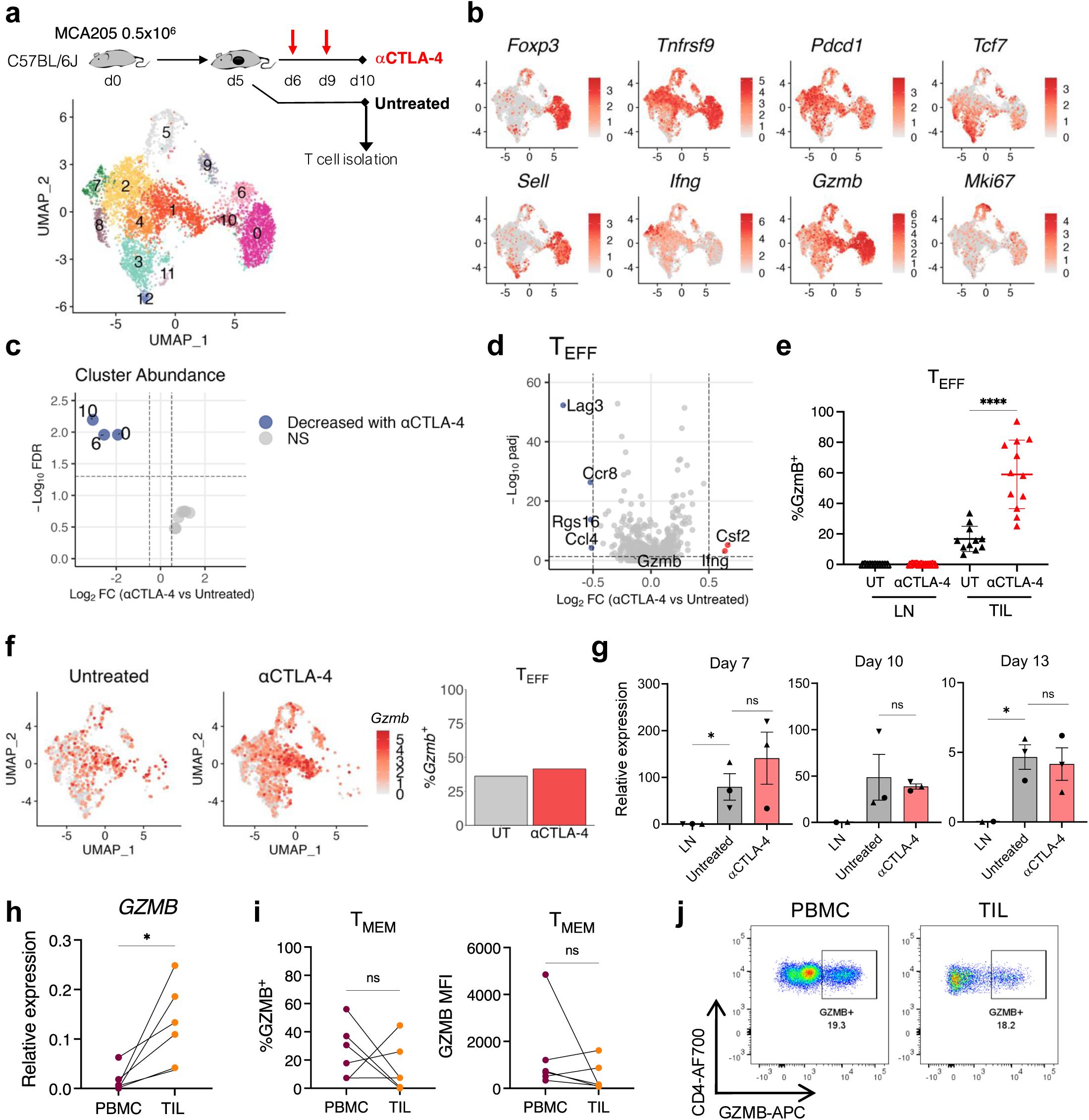
Poised CD4^+^ T_CTX_ are found in untreated tumors. **a,** Above: Schematic of the treatment strategy for scRNA-seq experiments. C57BL/6J mice challenged with MCA205 sarcoma were treated with anti-CTLA-4 on days 6 and 9 post-tumor inoculation or left untreated. T cells were isolated on day 10. Below: UMAP of *Cd4*-expressing TILs from untreated and anti-CTLA-4 treated mice (n=2592 untreated and n=3261 anti-CTLA-4, two independent experiments). **b,** Normalized expression of *Foxp3*, *Tnfrsf9*, *Pdcd1*, *Tcf7*, *Sell*, *Ifng*, *Gzmb* and *Mki67* per cell. **c,** Differential cluster abundance between untreated and anti-CTLA-4 treated tumors (FDR <0.05, |Log2FC|>0.5, quasi-likelihood F-test, edgeR). **d,** Volcano plot showing genes differentially expressed between MCA205-infiltrating T_EFF_ from untreated and anti-CTLA-4 treated mice (q <0.05, |Log_2_FC| > 0.5, MAST analysis). **e,** Proportion of GzmB^+^ CD4^+^ T_EFF_ in LN and MCA205 tumors treated with anti-CTLA-4 on days 6, 9 and 11 post-tumor inoculation or left untreated. Mice were sacrificed on day 12 (n=11/group, 2 independent experiments; one-way ANOVA). **f,** Left: UMAP of tumor-infiltrating *Cd4^+^* T_EFF_ from untreated and anti-CTLA-4 treated mice showing normalized *Gzmb* expression per cell (n=1246 untreated and n=2873 anti-CTLA-4, two independent experiments). Right: Proportions of *Cd4^+^* T_EFF_ positive for *Gzmb*^+^ mRNA in each condition. **g,** RT-qPCR of *Gzmb* mRNA in T_EFF_ (CD4^+^CD25^-^) from tumor-draining LN and MCA205 tumors from untreated and anti-CTLA-4 treated mice sacrificed at days 7, 10 and 13 post-tumor inoculation. Expression is relative to *Hprt1* (mean±SD of 3 independent experiments (each point is the average of 3-5 mice); two-tailed Student’s t-test). **h,** *GZMB* mRNA levels (relative to *ACTB)* from memory (T_MEM_) CD4^+^ T cells in PBMC and TILs of CRC patients (n=6, two-tailed paired t-test). **i,** Proportion of GzmB^+^ T_MEM_ and GZMB MFI in PBMC and TIL from all patients (n=6, two-tailed paired t-test). **j,** Concatenated dot plots showing GzmB protein levels in TMEM from PBMC and TILs of CRC patients.

We next assessed the impact of anti-CTLA-4 treatment on the balance between the different CD4^+^ TIL populations. Anti-CTLA-4 significantly depleted all T_REG_ populations, particularly those corresponding to activated T_REG_ (Fig. 1c; Extended Data Fig. 1e-g). Consistent with this, differential gene expression analysis revealed significant downregulation of genes such as *Lag3*, *Il10*, *Il2ra, Gzmb,* and *Entpd1* in T_REG_ from anti-CTLA-4 treated tumors (Extended Data Fig. 1h, Supplementary Table 2).

In contrast to the effect on T_REG_, we found that anti-CTLA-4 treatment had no effect on the relative abundance of any CD4^+^ T_EFF_ populations. Anti-CTLA-4 treatment also had limited impact on tumor-infiltrating T_EFF_ gene expression, only increasing expression of *Ifng* and *Csf2*, and downregulating *Lag3*, *Ccr8, Rsg16* and *Ccl4* (Fig. 1d, Supplementary Table 3). Furthermore, in direct contrast to the marked increase in GzmB protein upon anti-CTLA-4 treatment (Fig. 1e), we observed no difference in *Gzmb* mRNA between untreated and treated conditions (Fig. 1f and Extended Data Fig. 2a). We also found that expression levels of the cytotoxic genes *Nkg7, Gzma*, *Prf1* and *Gzmk* did not differ greatly between CD4^+^ T_EFF_ from untreated and anti-CTLA-4 tumors (Extended Data Fig. 2b).

To exclude the possibility that *Gzmb* expression in T_EFF_ increased at another timepoint during the course of anti-CTLA-4 treatment, we treated MCA205-bearing mice with anti-CTLA-4 on days 6, 9 and 12 post tumor inoculation and used RT-qPCR to measure *Gzmb* mRNA in sorted CD4^+^CD25^-^ T_EFF_ from tumors or draining LN on days 7, 10 or 13 (Extended Data Fig. 2c). *Gzmb* mRNA levels were increased in tumor-infiltrating T_EFF_ versus equivalent cells isolated from tumor-draining lymph nodes (LN) at all timepoints but there were no significant differences in *Gzmb* mRNA in tumor-infiltrating T_EFF_ between untreated and anti-CTLA-4 conditions (Fig. 1g). Similarly, CD4^+^ T_EFF_ from MC38 tumors, a colorectal carcinoma model which responds to anti-CTLA-4 T_REG_ depleting treatment^18,40^, contained comparable levels of *Gzmb* mRNA in untreated and anti-CTLA-4-treated conditions, whereas the proportion of GzmB^+^ T_EFF_ and levels (MFI) of GzmB protein in CD4^+^ T_EFF_ were higher in tumors from anti-CTLA-4 treated animals (Extended Data Fig. 2d-f). These data suggest that in untreated tumors a population of poised CD4^+^ T_CTX_ exists in which *Gzmb* is transcribed but production of GzmB protein is inhibited.

We next sought to determine if a similar population of poised, CD4^+^ T_CTX_-like cells were present in human tumors. We sorted naïve (CCR7^+^ CD45RA^+^; T_N_) and memory (CD45RA^-^; T_MEM_) populations from CD4^+^CD25^-^ PBMCs and TILs from patients with colorectal carcinoma (Extended Data Fig. 2g) and measured *GZMB* mRNA by RT-qPCR and GzmB protein by flow cytometry. T_N_ did not have detectable levels of *GZMB* mRNA and contained little GZMB protein in most patients (Extended Data Fig. 2h). Although *GZMB* mRNA levels were significantly higher in tumor-infiltrating CD4^+^ T_MEM_ than in peripheral CD4^+^ T_MEM_ from the same patients (Fig. 1h), there was no difference in the proportion of GzmB^+^ cells or GZMB MFI between the two CD4^+^ T_MEM_ populations (Fig. 1i,j). Thus, consistent with our findings in mouse tumor models, untreated human tumors contain a population of CD4^+^ T_EFF_ that contain higher levels of *GZMB* mRNA compared to T_EFF_ in peripheral blood but not higher levels of GzmB protein. Taken together, these data show that poised CD4^+^ T_CTX_ are present within untreated murine tumors and indicate that equivalent cells can be found in human tumors.

### Differentiation of poised CD4^+^ T_CTX_ is regulated by the Blimp-1-Bcl6 axis

The transcription factor Blimp-1 (encoded by *Prdm1*) is necessary for GzmB protein production by CD4^+^ TILs^7^. Blimp-1 also promotes CD4^+^ T cell cytotoxicity during viral infections, where its activity is antagonized by Bcl6^19,20^. This could indicate a role for the Blimp-1-Bcl6 axis in regulating differentiation into the poised CD4^+^ T_CTX_ state in tumors and/or the subsequent induction of GzmB protein production. Toexplore this, we challenged *Cd4*^Cre^, *Cd4*^Cre^ *Prdm1*^fl/fl^ (*Prdm1*^-/-^) and *Cd4*^Cre^ *Bcl6*^fl/fl^ (*Bcl6*^-/-^) mice with MCA205 and performed scRNA-seq on CD4^+^ TILs from untreated and anti-CTLA-4 treated animals. This revealed 13 populations of CD4^+^ TILs, similar to the populations that we had observed in WT mice (Extended Data Fig. 3a,b; Supplementary Table 4).

Tumor-infiltrating CD4^+^ T_EFF_ from *Prdm1*^-/-^ mice expressed lower levels of *Gzmb* mRNA than cells from *Cd4*^Cre^ control mice (Fig. 2a,b). Reciprocally, both the proportion of *Gzmb*-expressing T_EFF_ and the level of *Gzmb* mRNA were increased in Bcl6-deficient CD4^+^ T_EFF_. As observed in WT mice, *Gzmb* mRNA levels in T_EFF_ did not vary greatly between untreated or anti-CTLA-4 treated conditions in either *Cd4*^Cre^, *Bcl6*^-/-^ or *Prdm1*^-/-^ animals (Fig. 2a,b), suggesting that Blimp-1 and Bcl6 regulate *Gzmb* transcription in poised CD4^+^ T_CTX_ independently of treatment. Furthermore, although Bcl6 deficiency induced *Gzm*b expression at the RNA level, it had no significant effect on GzmB protein levels in either untreated or anti-CTLA-4 treated tumors, despite efficient T_REG_ depletion in this model (Fig. 2c,d). These observations demonstrate that the Blimp-1-Bcl6 axis regulates differentiation of CD4^+^ cells into poised T_CTX_ in untreated tumors and shows that establishment of this state at a transcriptional level is insufficient for GzmB protein production and acquisition of cytotoxic activity.

**Fig. 2:**
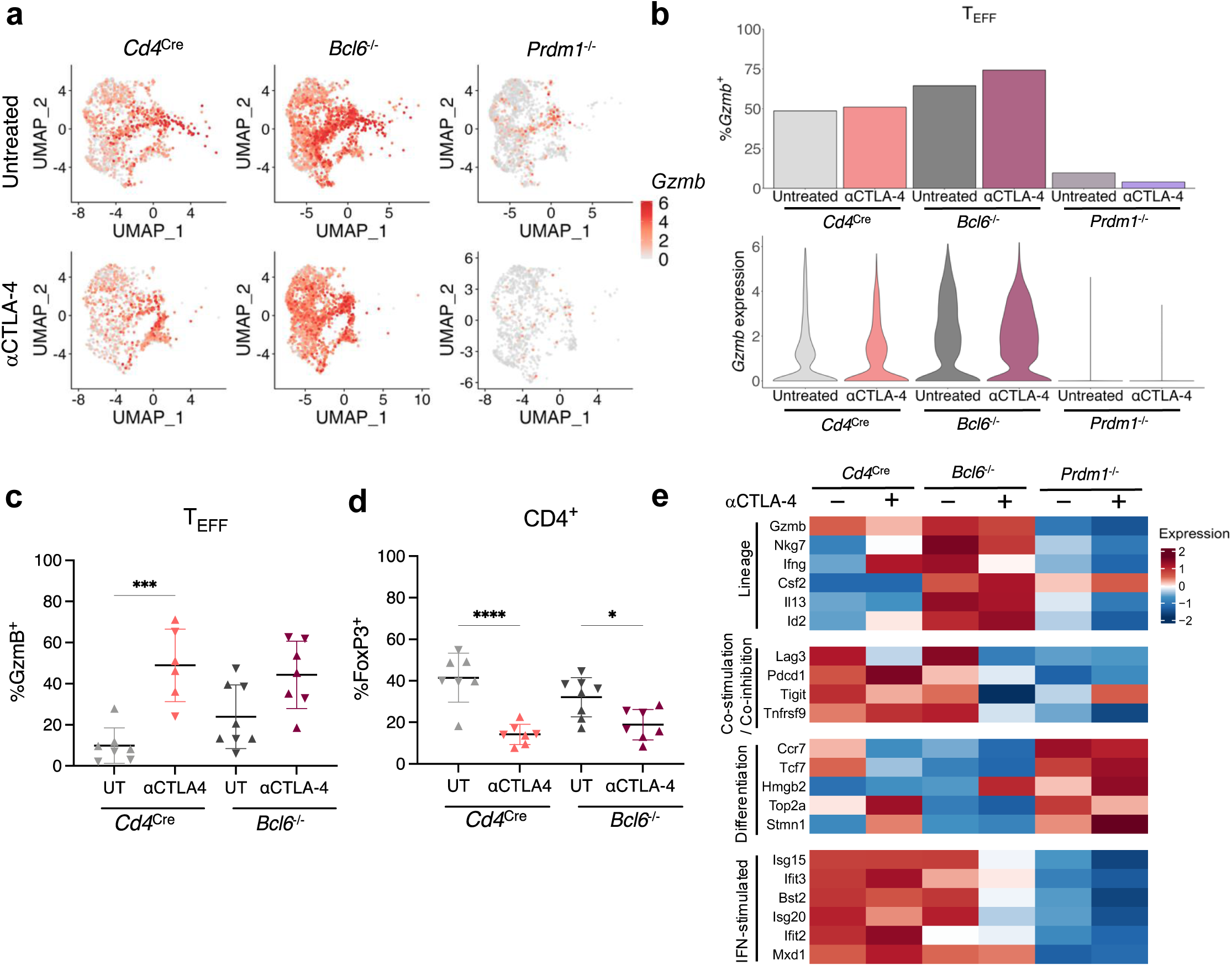
Acquisition of a cytotoxic transcriptional program in poised CD4^+^ T_CTX_ is regulated by the Blimp-1-Bcl6 axis. **a,** UMAP of MCA205-infiltrating T_EFF_ from untreated (n=1830 *Cd4*^Cre^, 2898 *Bcl6*^-/-^ and 1910 *Prdm1*^-/-^; two independent experiments) and anti-CTLA-4 treated (n=1427 *Cd4*^Cre^, 2597 *Bcl6*^-/-^ and 1093 *Prdm1*^-/-^) *Cd4*^Cre^, *Bcl6*^-/-^ or *Prdm1*^-/-^ mice, showing normalized *Gzmb* expression per cell. Mice were treated on days 6 and 9 post-tumor inoculation and sacrificed on day 10. **b,** Proportion of *Gzmb*^+^ *Cd4*^+^ T_EFF_ (above) and average *Gzmb* expression (below) in each treatment condition from a. **c-d,** Proportion of (**c**) GzmB^+^ CD4^+^ T_EFF_ and (**d**) FoxP3^+^ CD4^+^ T cells in MCA205 tumors from *Cd4*^Cre^ and *Bcl6*^-/-^ mice treated with anti-CTLA-4 on days 6, 9 and 11 or left untreated and sacrificed at day 12 post-tumor inoculation (n=7-8/group from two independent experiments, one-way ANOVA). **e,** Heatmap showing average expression of selected genes in each treatment condition from a. Genes shown are differentially expressed in *Cd4*^+^ T_EFF_ between *Cd4*^Cre^ and *Prdm1*^-/-^ or *Cd4*^Cre^ and *Bcl6*^-/-^ animals in either treatment condition (q <0.05, |Log_2_FC| > 0.5, MAST analysis).

To investigate further the roles of Bcl6 and Blimp-1 in regulating the gene expression programs of CD4^+^ TILs, we performed differential gene expression analysis between *Cd4*^Cre^ and *Bcl6*^-/-^ T_EFF_ and between *Cd4*^Cre^ and *Prdm1*^-/-^ T_EFF_ in both untreated and anti-CTLA-4 treated conditions. *Bcl6*^-/-^ T_EFF_ upregulated genes associated with multiple T_H_ fates, including T_H_1 (*Ifng*, *Csf2*) and T_H_2 (*Il13*), suggesting that Bcl6 broadly antagonized CD4^+^ T cell differentiation (Fig. 2e, Extended Data Fig. 3c and Supplementary Table 5). Blimp-1-deficient T_EFF_ exhibited increased expression of genes associated with cell cycle progression (*Hmgb2*, *Top2a*, *Stmn1*) and decreased expression of genes encoding the checkpoint molecules LAG-3, PD-1, TIGIT and 4-1BB, consistent with a broad role for Blimp-1 in promoting terminal differentiation. This analysis also showed that the high level of *Gzmb* expression in *Cd4*^Cre^ and *Bcl6*^-/-^ T_EFF_ was accompanied by high expression of interferon stimulated genes (ISGs), which were also significantly reduced in the absence of Blimp-1 (Fig. 2e). We conclude that differentiation of poised CD4^+^ T_CTX_ is promoted by Blimp-1 and antagonized by Bcl6, which further regulate the balance between terminal differentiation and proliferation of tumor-infiltrating CD4^+^ T cells.

### Differentiation into poised CD4^+^ T_CTX_ requires type I IFN signaling

We considered that co-regulation of *Gzmb* and ISG expression may indicate a role for type I IFN signaling in the differentiation of CD4^+^ TILs into poised T_CTX_. Consistent with this, differential gene expression analysis of *Gzmb*-expressing vs non-expressing CD4^+^ T_EFF_ in MCA205 tumors from WT animals revealed upregulation of ISGs in *Gzmb*-expressing cells (Fig. 3a). To test for a role for type I IFN signaling, we treated MCA205-bearing mice with a blocking anti-IFNAR antibody (Fig. 3b). As we have previously shown that IL-2 is necessary for differentiation into CD4^+^ T_CTX_ in tumors^7^, we treated additional mice with a neutralizing anti-IL-2 antibody to determine whether IL-2 signaling regulates CD4^+^ T_CTX_ differentiation at the transcriptional level.

**Fig. 3:**
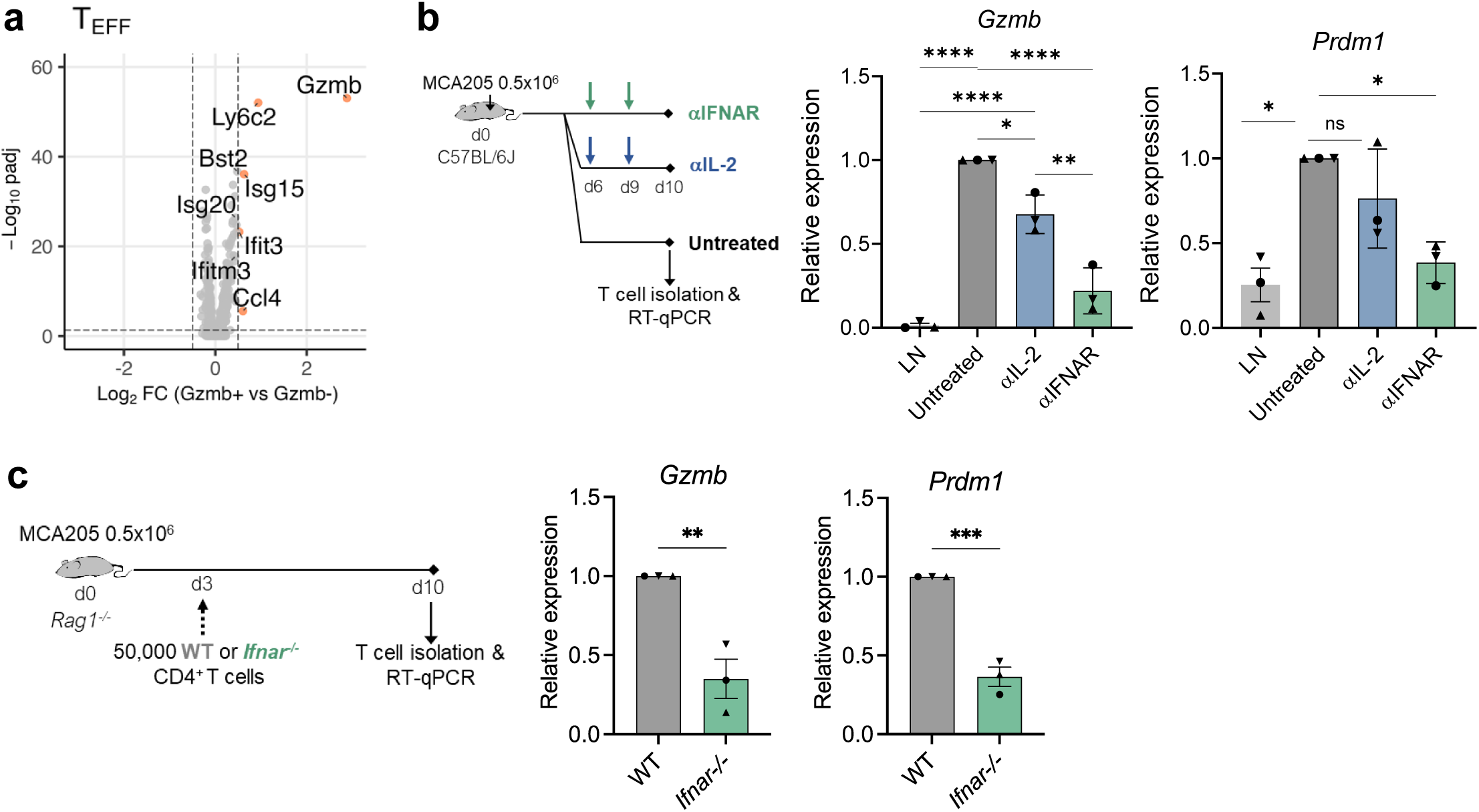
Differentiation into poised T_CTX_ is induced by type I IFN signaling in tumors. **a,** Volcano plot showing genes differentially expressed between *Gzmb*^+^ and *Gzmb*^-^ *Cd4^+^* T_EFF_ from MCA205 tumors (q <0.05, |Log_2_FC| > 0.5, MAST analysis). **b,** Left: schematic of treatment strategy for type I IFN and IL-2 blocking experiments. C57BL/6J mice challenged with MCA205 sarcoma were treated with anti-IFNAR or anti-IL-2 on days 6 and 9 post-tumor inoculation or left untreated. Mice were sacrificed and T cells isolated on day 10. Right: RT-qPCR analysis of *Gzmb* and *Prdm1* expression in CD4^+^ T_EFF_ from tumor-draining LN of untreated mice and MCA205 tumors from mice treated with anti-IL-2 or anti-IFNAR. Expression is relative to *Hprt1* and normalized to untreated TILs (mean±SD of 3 independent experiments, each point is the average of 3-5 mice; Welch’s t-test). **c,** Left: schematic of treatment strategy for WT or *Ifnar^-/-^* T cell transfer experiments. WT or *Ifnar^-/-^* CD4^+^ T_EFF_ were adoptively transferred into MCA205 tumor-bearing *Rag1^-/-^* mice on day 3 post-tumor inoculation. Mice were sacrificed and T cells isolated on day 10. Right: RT-qPCR analysis of *GzmB* and *Prdm1* expression in WT or *Ifnar*^-/-^ CD4^+^ T_EFF_ isolated from MCA205 tumors. Expression is relative to *Hprt1* and normalized to WT (mean±SD of 3 independent experiments, each point is the average of 3-5 mice; Welch’s t-test).

We found that treatment of MCA205 tumors with anti-IL-2 significantly reduced *Gzmb* expression in T_EFF_ when compared to untreated animals but did not have a significant effect on *Prdm1* expression (Fig. 3b). Strikingly, anti-IFNAR treatment reduced *Gzmb* expression even further and significantly reduced expression of *Prdm1* in CD4^+^ T_EFF_. This suggests that type I IFN signaling, and to a lesser extent IL-2 signaling, are required for establishment of the poised CD4^+^ T_CTX_ state in tumors.

To determine whether type I IFN signaling is required for CD4^+^ T_CTX_ differentiation in a cell-intrinsic manner, we transferred CD4^+^ T cells from WT or *Ifnar^-/-^* mice into MCA205 tumor-bearing *Rag1*^-/-^ mice (Fig. 3c). We found that tumor-infiltrating *Ifnar*^-/-^ CD4^+^ T cells expressed significantly lower levels of *Gzmb* and *Prdm1* than WT cells (Fig. 3c). Thus, type I IFN signaling is required in CD4^+^ T cells for differentiation into poised T_CTX_ in tumors.

### Blockade of LAG-3 and PD-1 promotes GzmB protein production in poised CD4^+^ T_CTX_

Having identified that GzmB protein production is repressed by a post-transcriptional checkpoint in tumor-infiltrating CD4^+^ T_EFF_, we next wished to investigate how this may be regulated. As depletion of activated T_REG_ from tumors by anti-CTLA-4 was sufficient to induce GzmB protein production, we searched for markers on CD4^+^ T_EFF_ that were altered by anti-CTLA-4 treatment. We noted that anti-CTLA-4 treatment caused a significant downregulation of *Lag3* expression (Fig. 1d), as well as a decrease in the proportion of LAG-3^+^ GzmB^+^ CD4^+^ T_EFF_ at the protein level (Fig. 4a, Extended Data Fig. 4a). Previous studies have shown a role for LAG-3 and PD-1 in inhibiting differentiation into CD4^+^ T_CTX_ following viral infections^19,41^, suggesting that LAG-3 and PD-1 may block GzmB protein production in untreated tumors.

**Fig. 4:**
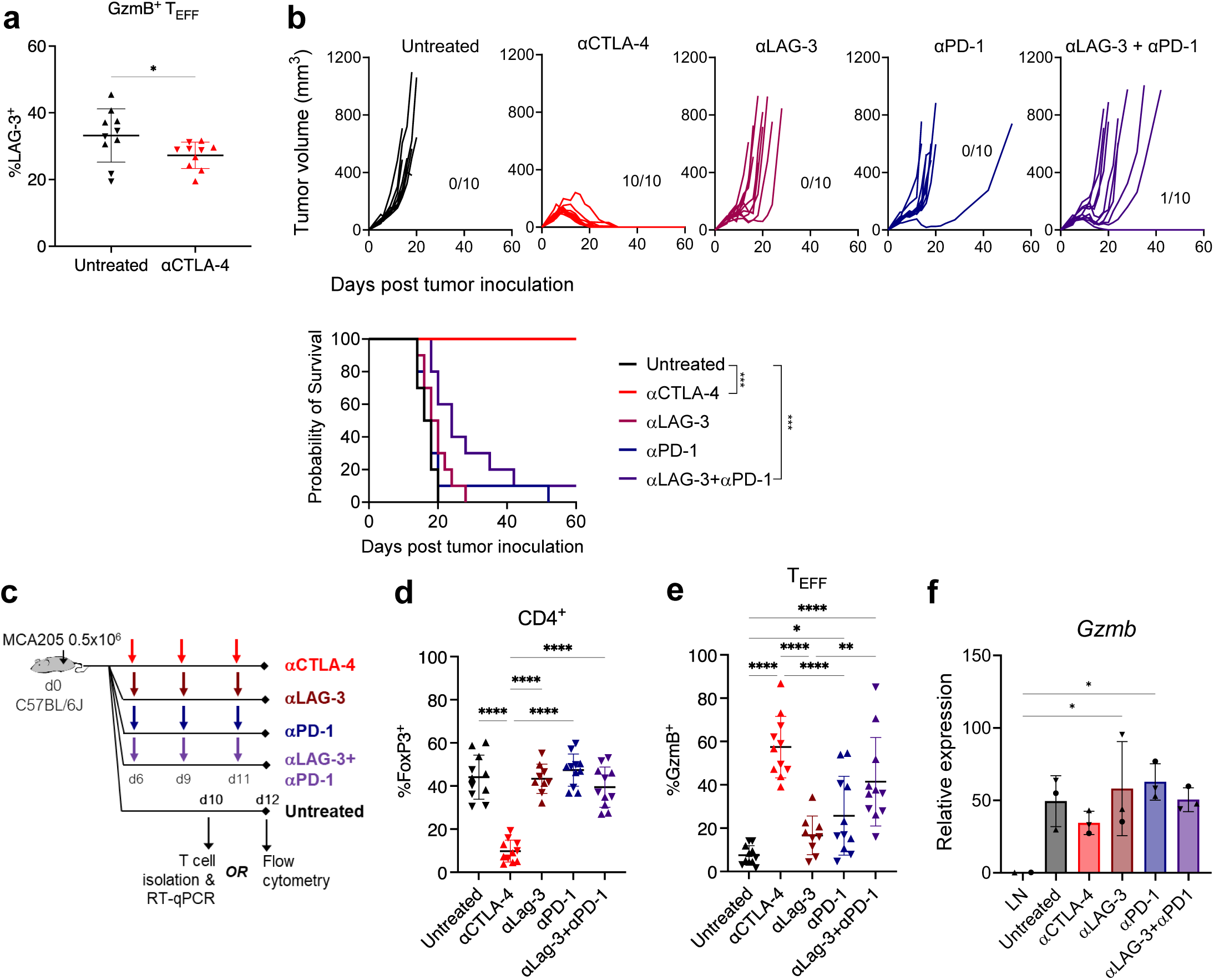
Blockade of LAG-3 and PD-1 induces GzmB protein production in poised CD4^+^ T_CTX_. **a,** Proportions of GzmB^+^ MCA205-infiltrating CD4^+^ T_EFF_ positive for LAG-3 in mice treated with anti-CTLA-4 on days 6, 9 and 11 post-tumor challenge or left untreated. Mice were sacrificed on day 12 (n=10/group from 2 independent experiments, Student’s t-test). **b,** Above: MCA205 tumor volume in untreated, anti-CTLA-4, anti-LAG-3, anti-PD-1 or anti-LAG3+anti-PD-1 treated mice, with the number of mice surviving to day 60 post-tumor inoculation indicated (n=10/group). Below: Probability of survival of untreated, anti-CTLA-4, anti-LAG-3, anti-PD-1 or anti-LAG3+anti-PD-1 treated mice; ***p<0.001 vs untreated, Mantel-Cox log-rank test). **c,** Schematic of treatment strategy for LAG-3/PD-1 blocking experiments. For mRNA analysis, C57BL/6J mice challenged with MCA205 sarcoma were treated with indicated antibodies on days 6 and 9 post-tumor inoculation or left untreated and mice were sacrificed and T cells isolated on day 10. Alternatively, for flow cytometry analysis, MCA205 tumor-bearing mice were treated with indicated antibodies on days 6, 9 and 11 post-tumor inoculation or left untreated and mice were sacrificed and tumors and lymph nodes analyzed on d12. **d-e,** Proportion of (**d**) FoxP3^+^ CD4^+^ T cells (T_REG_) and (**e**) CD25^+^ T_REG_ in MCA205 tumors from mice treated with anti-CTLA-4, anti-LAG-3, anti-PD-1 or anti-LAG3+anti-PD-1 on days 6, 9 and 11 post-tumor inoculation or left untreated (n=9-11/group from 2 independent experiments, one-way ANOVA). **f,** RT-qPCR analysis of *Gzmb* expression in CD4^+^ T_EFF_ from tumor-draining LN and MCA205 tumors treated with anti-CTLA-4, anti-LAG-3, anti-PD-1 or anti-LAG-3+anti-PD-1. Expression is relative to *Hprt1* (mean±SD of 3 independent experiments (each point is the average of 3-5 mice); one-way ANOVA).

We first asked whether LAG-3 and PD-1 impact tumor growth and survival in the MCA205 model by treating mice with anti-LAG-3, anti-PD-1 or anti-LAG3+anti-PD1 in combination, or anti-CTLA-4, on days 6, 9 and 12 post-tumor challenge. We found that treatment with anti-LAG-3 or anti-PD-1 alone had no effect on survival but that the combination treatment significantly extended survival when compared to untreated animals, although tumors eventually escaped immune control and grew in 9 out of 10 animals (Fig. 4b).

We next investigated whether LAG-3 and PD-1 regulated GzmB protein production in CD4^+^ TILs (Fig. 4c). In contrast to the effect of anti-CTLA-4, treatment with anti-LAG-3, anti-PD-1, or anti-LAG-3+anti-PD-1 did not deplete tumor-infiltrating T_REG_ (Fig. 4d). Anti-LAG3+anti-PD-1 treatment decreased the proportion of activated CD25^+^ and KLRG1^+^ T_REG_ in MCA205 tumors, albeit only by 6% and 7%, respectively (Extended Data Fig. 4b,c). Treatment with anti-PD1 and anti-LAG-3+anti-PD-1 significantly increased the proportion of GzmB-protein producing CD4^+^ T_EFF_ when compared to untreated animals or LAG-3 treatment alone (Fig. 4e), albeit to a lesser extent than observed following anti-CTLA-4 treatment. In contrast to the effect on GzmB protein, anti-LAG-3+anti-PD-1 blockade had no effect on *Gzmb* mRNA levels (Fig. 4c,f), consistent with post-transcriptional regulation. Unlike anti-CTLA-4 treatment, blockade of LAG-3+PD-1 had no effect on phospho-STAT5 or phospho-STAT3 levels within tumor-infiltrating CD4^+^ T_EFF_, suggesting the treatment does not upregulate GzmB protein by increasing IL-2R signaling (Extended Data Fig. 4d). We conclude that although *Gzmb* mRNA is transcribed by CD4^+^ T_EFF_ in untreated tumors, GzmB protein production is repressed through the activity of the co-inhibitory checkpoint molecules LAG-3 and PD-1.

### ZFP36L1 represses GzmB protein production in CD4^+^ TILs

We next explored how GzmB protein production was blocked in poised CD4^+^ T_CTX_ in untreated tumors and how checkpoint inhibitor treatment overcomes this suppression. In other contexts, the RNA binding proteins (RBPs) ZFP36 and ZFP36L1 act redundantly at a post-transcriptional level to inhibit T cell effector function^31,33,34,36^. We therefore considered that these RBPs may also regulate GzmB protein production in tumor-infiltrating CD4^+^ T_EFF_. Consistent with this hypothesis, we found that anti-CTLA-4 and anti-LAG-3+anti-PD-1 treatment of MCA205 tumors significantly decreased expression of *Zfp36l1* in CD4^+^ T_EFF_ (Fig. 5a).

**Fig. 5:**
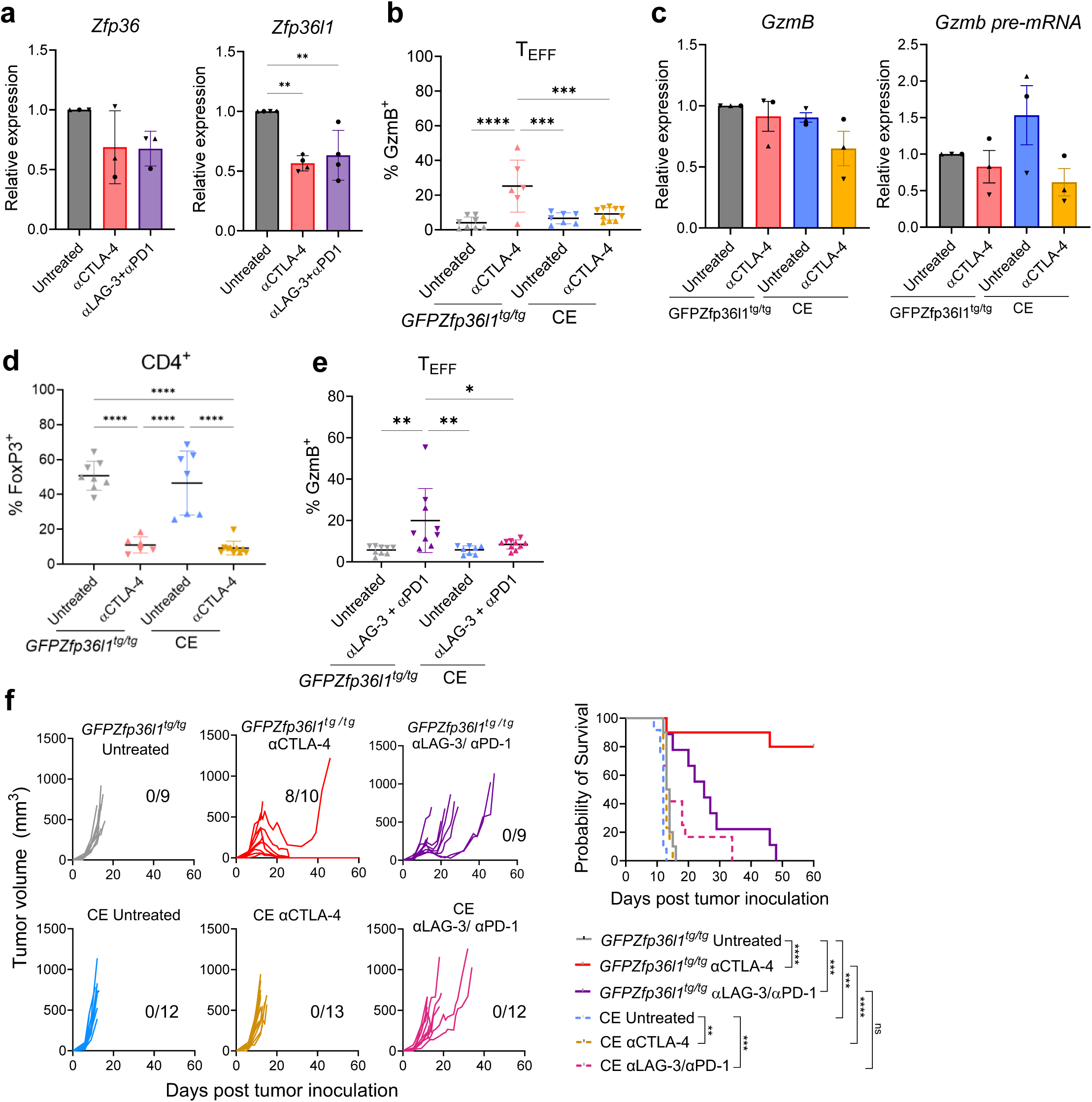
ZFP36L1 blocks GzmB protein production in tumor-infiltrating CD4^+^ T_EFF_. **a,** RT-qPCR analysis of *Zfp36 and Zfp36l1* expression in CD4^+^ T_EFF_ from MCA205 tumors from mice treated with anti-CTLA-4 or anti-LAG-3+anti-PD-1 on days 6 and 9 post-tumor inoculation, or left untreated, and sacrificed on day 10. Expression is relative to *Hprt1* (mean±SD of 3 independent experiments (each point is the average of 3-5 mice); one-way ANOVA). **b,** Proportion of GzmB^+^ CD4^+^ T_EFF_ in MCA205 tumors of *Cd4^Cre^ GFPZfp36l1^tg/tg^* (CE) or *GFPZfp36l1^tg/tg^* littermates treated with anti-CTLA-4 on days 6, 9 and 11 post-tumor inoculation or left untreated. Mice were sacrificed on day 12 (n=6-10/group from 2 independent experiments, one-way ANOVA). **c,** RT-qPCR analysis of *Gzmb* mRNA (left) and pre-mRNA (right) in CD4^+^ T_EFF_ from tumors of untreated or anti-CTLA-4-treated CE or littermate animals. Mice were sacrificed at day 10 post-tumor inoculation. Expression is relative to *Hprt1* (mean±SD of 3 independent experiments (each point is the average of 3-5 mice); Student’s t-test). **d,** Proportion of FoxP3^+^ CD4^+^ T cells (T_REG_) in MCA205 tumors. Details as for b. **e,** Proportion of GzmB^+^ CD4^+^ T_EFF_ in MCA205 tumors from CE or littermate animals treated with anti-LAG-3 + anti-PD-1 on days 6, 9 and 11 post-tumor inoculation or left untreated (n=8-10/group from 2 independent experiments, one-way ANOVA). **f,** Left: MCA205 tumor volume in untreated CE or littermate mice treated with anti-CTLA-4 or anti-LAG-3 + anti-PD-1 on days 6, 9 and 12 post-tumor inoculation or left untreated, with the number of mice surviving to day 60 post-tumor inoculation indicated (n=9-12/group from two independent experiments). Right: Probability of survival between treatments or between genotypes (**p<0.01; ***p<0.001; ****p<0.0001, Mantel-Cox log-rank test).

We reasoned that this downregulation of *Zfp36l1* expression in CD4^+^ T_EFF_ could be necessary for the induction of GzmB protein production by anti-CTLA-4. To test this, we crossed *Cd4^Cre^ with GFPZfp36l1^tg/tg^* mice^42^ to generate a line in which *Zfp36l1* is constitutively expressed in T cells (henceforth CE mice; Extended Data Fig. 5a,b). We then challenged CE and *GFPZfp36l1^tg/tg^* littermate controls with MCA205 and treated the animals with anti-CTLA-4 as before. Strikingly, anti-CTLA-4 treatment failed to increase GzmB protein in tumor-infiltrating CD4^+^ T_EFF_ in CE mice (Fig. 5b and Extended Data Fig. 5c). The failure to increase GzmB protein was not due to changes at the mRNA or pre-mRNA levels (Fig. 5c) or defects in T_REG_ depletion in anti-CTLA-4-treated tumors (Fig. 5d and Extended Data Fig. 5d). These data thus support our hypothesis that *Zfp36l1* downregulation is necessary for GzmB protein production triggered by anti-CTLA-4. Consistent with this, CD4^+^ T cells purified from spleen and LN exhibited reduced GzmB protein production after activation in vitro showing that the suppressive effect of *Zfp36l1* constitutive expression was intrinsic to CD4^+^ T cells (Extended Data Fig. 5e).

In addition to GzmB, *Zfp36l1* constitutive expression also blocked upregulation of the known ZFP36 family target IFNγ upon *ex vivo* re-stimulation of tumor-infiltrating CD4^+^ T_EFF_ (Extended Data Fig. 5f). Further examining the effects of *Zfp361* constitutive expression, we found that tumors established in CE mice contained similar numbers of CD4^+^ T cells as WT mice but did not show an increase in T cell numbers following anti-CTLA-4 treatment (Extended Data Fig. 5g). Tumor-infiltrating CD4^+^ T_EFF_ from CE mice showed reduced levels of the activation markers ICOS and 4-1BB in response to anti-CTLA-4, as well as elevated levels of the inhibitory receptors PD-1 and TIM-3 (Extended Data Fig. 5h), suggesting decreased functionality of CE cells. The repressive effect of *Zfp36l1* constitutive expression on GzmB protein was also apparent in the CD8^+^ T cell compartment (Extended Data Fig. 5i).

We next asked whether *Zfp36l1* downregulation was also necessary for upregulation of GzmB protein production in response to anti-LAG-3 and anti-PD-1. We found that, as for anti-CTLA-4 treatment, constitutive expression of *Zfp36l1* prevented GzmB induction by anti-LAG-3+anti-PD-1 in both CD4^+^ T_EFF_ and CD8^+^ T cells (Fig. 5e and Extended Data Fig. 5j). Thus, the ability of these checkpoint inhibitors to increase GzmB protein production also required downregulation of *Zfp36l1*. Constitutive *Zfp36l1* expression also prevented the induction of proliferation and activation markers, and increased inhibitory receptor expression, in CD4^+^ T_EFF_ in response to anti-LAG-3+anti-PD-1 treatment (Extended Data Fig. 5k).

Given the impact of constitutive *Zfp36l1* expression on GzmB protein, as well as on cytokines, activation markers and inhibitory receptors, we considered that ZFP36L1 might also abrogate the ability of checkpoint inhibitors to control tumor growth. Indeed, we found that *Zfp36l1* constitutive expression completely abrogated the ability of anti-CTLA-4 treatment to induce MCA205 tumor regression and increase survival (Fig. 5f). *Zfp36l1* constitutive expression also blunted the effect of anti-LAG-3+anti-PD-1: tumors grew more rapidly in CE mice than in littermate controls and survival was shortened, although this did not reach significance. Thus, *Zfp36l1* downregulation is necessary to overcome the block to cytotoxic activity in tumor-infiltrating CD4^+^ T_EFF_ and for the anti-tumor response elicited by anti-CTLA-4.

### Targeting ZFP36 and ZFP36L1 increases GzmB protein production in CD4^+^ TILs and reduces tumor growth

The prevention of GzmB protein production caused by constitutive *Zfp36l1* expression suggests that targeting *Zfp36l1* could enhance CD4^+^ T_CTX_ activity in tumors. To test this, and to account for reported redundancies between ZFP36L1 and its paralog ZFP36^31^, we challenged *Cd4*^Cre^*Zfp36*^fl/fl^*Zfp36l1*^fl/fl^ (henceforth DKO) mice and *Zfp36*^fl/fl^*Zfp36l1*^fl/fl^ littermates with MCA205 and treated them with anti-CTLA-4 or left them untreated as before. Strikingly, double knockout of *Zfp36* and *Zfp36l1* released suppression of GzmB protein production by poised CD4^+^ T_CTX_ (Fig. 6a and Extended Data Fig. 6a). Tumors from untreated DKO mice contained a significantly higher proportion of GzmB^+^ cells and increased GzmB MFI in the CD4^+^ T_EFF_ population when compared with littermate controls. In contrast, we observed no significant difference in GzmB levels between DKO and littermates following anti-CTLA-4 treatment (Fig. 6a), suggesting that ZFP36 and ZFP36L1 limit CD4^+^ T_CTX_ function primarily in the untreated state where their expression is elevated (Fig. 5a). Similar effects were seen in CD8^+^ T cells (Extended Data Fig. 6b) and for IFNγ in ex vivo re-stimulated CD4^+^ T_EFF_ (Extended Data Fig. 6c).

**Fig. 6:**
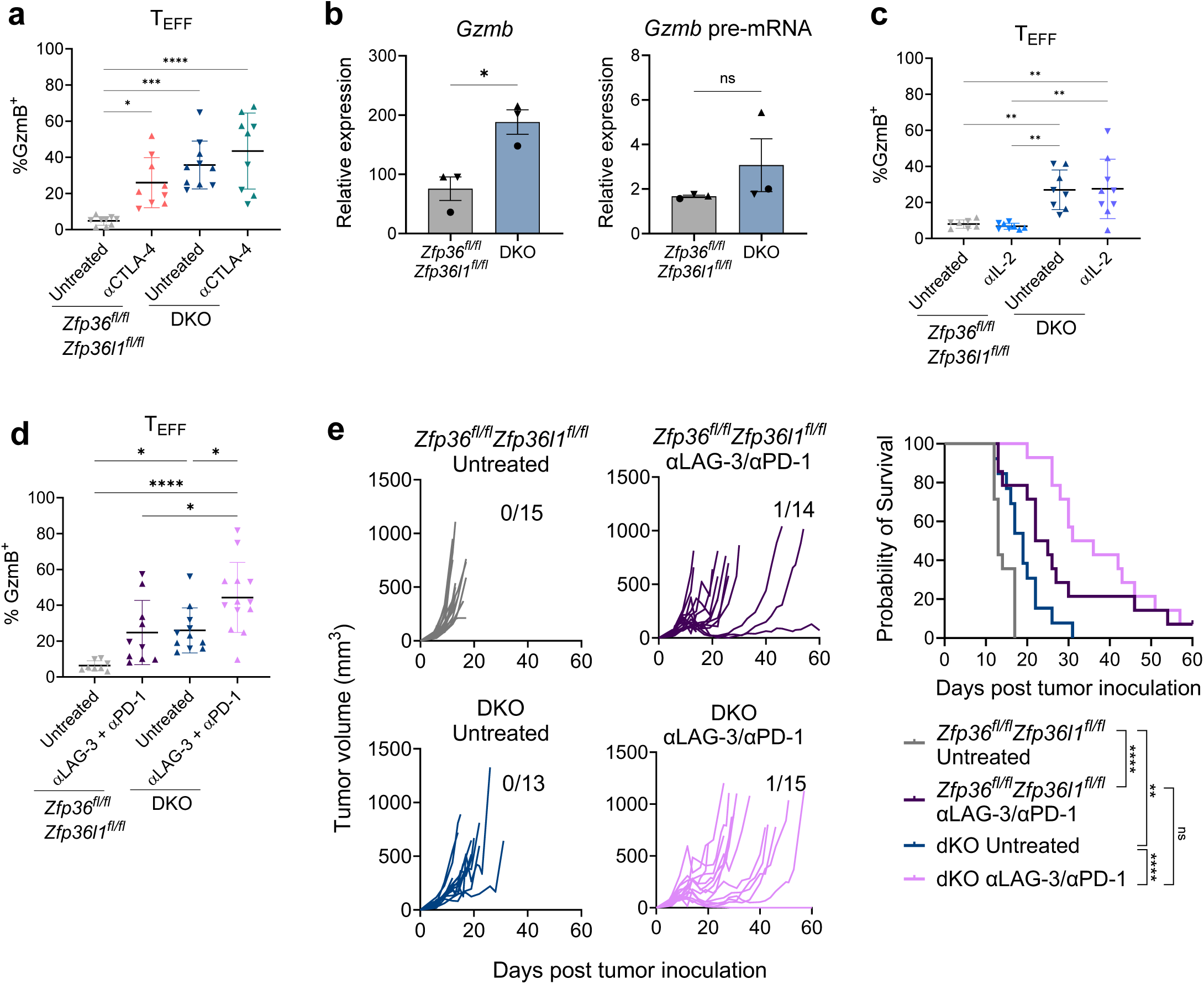
Targeting ZFP36 and ZFP36L1 increases GzmB protein production in tumor-infiltrating CD4^+^ T_EFF_ and reduces tumor growth. **a,** Proportion of GzmB^+^ CD4^+^ T_EFF_ in MCA205 tumors of *Cd4*^Cre^ *Zfp36*^fl/fl^ *Zfp36l1*^fl/fl^ (DKO) or *Zfp36*^fl/fl^ *Zfp36l1*^fl/fl^ littermate controls treated with anti-CTLA-4 on days 6, 9 and 11 post-tumor inoculation or left untreated, and sacrificed on day 12 (n=9-10/group from two independent experiments; one-way ANOVA). **b,** RT-qPCR analysis of *Gzmb* mRNA (left) and *Gzmb* pre-mRNA (right) in CD4^+^ T_EFF_ from tumors of DKO or littermate animals. Mice were sacrificed at day 10 post-tumor inoculation. Expression is relative to *Hprt1* (mean±SD of 3 independent experiments (each point is the average of 3-5 mice); Student’s t-test). **c,** Proportion of GzmB^+^ CD4^+^ T_EFF_ in MCA205 tumors of DKO or littermate controls treated with anti-IL2 on days 6, 9 and 11 post-tumor inoculation or left untreated, and sacrificed on day 12 (n=9-10/group from two independent experiments; one-way ANOVA). **d,** Proportion of GzmB^+^ CD4^+^ T_EFF_ in MCA205 tumors from mice treated with anti-LAG3+anti-PD-1 on days 6, 9 and 11 post-tumor inoculation or left untreated, and sacrificed on day 12 (n=8-13/group, one-way ANOVA). **e,** Left: MCA205 tumor volume in untreated DKO or littermate mice treated with anti-LAG3+anti-PD-1 on days 6, 9 and 12 post-tumor inoculation or left untreated, with the number of mice surviving to day 60 post-tumor inoculation indicated (n=13-15/group from four independent experiments). Right: Probability of survival between treatments or between genotypes (**p<0.01, ****p<0.0001, Mantel-Cox log-rank test).

We observed no differences in the number of tumor-infiltrating CD4^+^ or CD8^+^ T cells between DKO and littermate animals, suggesting no effect on T cell infiltration (Extended Data Fig. 6d). No changes in the expression of the activation markers ICOS and 4-1BB or the inhibitory receptors LAG-3, PD-1 or TIM-3 were apparent in CD4^+^ T_EFF_ from untreated tumors from DKO and littermate mice (Extended Data Fig. 6e). The proportion of total T_REG_, as well as activated CD25^+^ T_REG_, were also comparable between untreated DKO animals and littermates (Extended Data Fig. 6f), suggesting that the increase in GzmB protein production in DKO animals was not caused by defects in T_REG_ infiltration or activation.

CD4^+^ T_EFF_ purified from spleen and LN of DKO mice exhibited increased GzmB protein production after *in vitro* activation compared to cells purified from littermate controls, demonstrating this to be a cell autonomous effect (Extended Data Fig. 6g). RT-qPCR for *Gzmb* mRNA in CD4^+^ T_EFF_ from untreated tumors showed elevated levels of mature transcripts in DKO mice compared to littermates but comparable levels of pre-mRNA (Fig. 6b), consistent with the role of these RBPs in regulating mRNA stability without affecting transcription^27,28,29^.

The elevated production of GzmB by T_EFF_ in untreated tumors in DKO animals suggested that loss of ZFP36/ZFP36L1 increased GzmB protein production by bypassing the dependence on IL-2, which is limiting in untreated tumors^7^. Indeed, we found that IL-2 neutralization had no effect on GzmB production by tumor-infiltrating CD4^+^ T_EFF_ in DKO animals (Fig. 6c), demonstrating that GzmB induction caused by loss of ZFP36/ZFP36L1 is independent of IL-2.

We next tested the effect of anti-LAG-3+anti-PD-1 treatment on GzmB production in DKO and control mice harboring MCA205 tumors. We found that the increased proportion of GzmB^+^ CD4^+^ T_EFF_ observed in untreated DKO mice was further elevated upon LAG-3+PD-1 blockade (Fig. 6d). This was also the case for CD8^+^ T cells and occurred without an effect on tumor-infiltrating T_REG_ numbers (Extended Data Fig. 6h,i).

Our results suggested that targeting ZFP36L1/ZFP36 could promote tumor control, even within an immunosuppressive microenvironment. Indeed, we found that tumors grew slower in DKO mice compared to littermate controls and that survival was significantly extended (Fig. 6e). Furthermore, survival of DKO mice was further extended by treatment with anti-LAG-3+anti-PD-1 (Fig. 6e). Taken together with our other results, these data support a key role for ZFP36 and ZFP36L1 in mediating a post-transcriptional checkpoint that limits the cytotoxic activity of poised CD4^+^ T cells and indicate that targeting this pathway could provide a means to limit tumor growth.

## Discussion

Our data support a model in which acquisition of cytotoxic activity by tumor-infiltrating CD4^+^ T_EFF_ occurs in a two-step process (Fig. 7). In untreated tumors, CD4^+^ T_EFF_ acquire a cytotoxic gene expression program that is regulated by the Blimp-1-Bcl6 axis in response to type I IFN. However, although these cells express *Gzmb* mRNA, GzmB protein production is restrained by a post-transcriptional checkpoint dependent on ZFP36 and ZFP36L1. This post-transcriptional checkpoint can be overcome therapeutically. Treatment with anti-CTLA-4 or with anti-LAG-3+anti-PD-1 downregulates *Zfp36l1*, which allows GzmB protein production. Preventing *Zfp36l1* downregulation prevents anti-CTLA-4 and anti-LAG-3+anti-PD-1 from inducing GzmB and nullifies the ability of anti-CTLA-4 to induce tumor regression. Reciprocally, knockout of *Zfp36l1* and *Zfp36* in T cells induces GzmB protein production in CD4^+^ T_EFF_ in untreated tumors and promoted tumor control. Thus, our results suggest that this post-transcriptional checkpoint is a promising target for future therapeutic strategies.

**Fig. 7:**
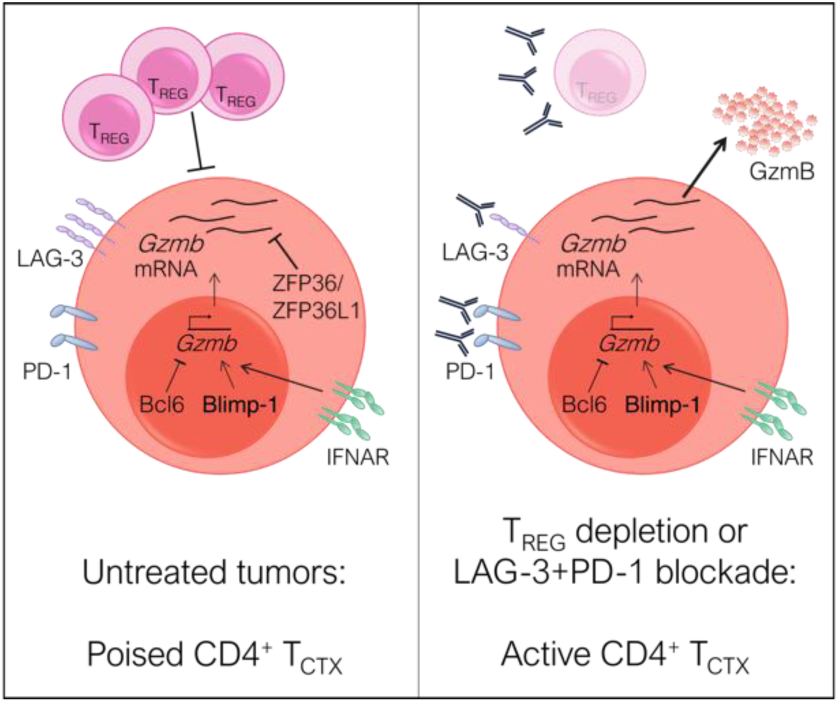
Two-step model for acquisition of cytotoxic activity by tumor-infiltrating CD4^+^ T cells. In untreated tumors CD4^+^ T_EFF_ acquire a cytotoxic gene expression program, which is regulated by the Blimp-1-Bcl6 axis in response to type-I IFN signals. However, GzmB protein production is blocked by ZFP36 and ZFP36L1. Anti-CTLA-4-mediated T_REG_ depletion or LAG-3/PD-1 blockade downregulate *Zfp36l1*, releasing the block to GzmB protein production, and reduces tumor growth.

Our work reveals the existence of a poised population of CD4^+^ T_CTX_ in untreated tumors that contain high levels of *Gzmb* mRNA but fail to produce GzmB protein. We did not detect these poised CD4^+^ T_CTX_ in tumor-draining lymph nodes, suggesting that the cells differentiate inside the tumor microenvironment in response to tumor-specific signals. We also identified a lack of correlation between the levels of GzmB mRNA and protein in tumor-infiltrating CD4^+^ T cells from patients, suggesting GzmB protein levels are also under post-transcriptional regulation in humans. A similar disconnect between total mRNA and protein levels has been observed for other genes in T cells following *in vitro* stimulation^30,43,44,45,46,47^, suggesting additional T cell effector functions may be similarly regulated in tumors.

We found that poised CD4^+^ T_CTX_ differentiate in tumors in response to type-I IFNs. Type-I IFNs play a complex role in tumors; they contribute to pro-tumor chronic inflammation, while also initiating anti-tumor immune responses^48^. Blockade of IFNα/β signaling in untreated tumors reduced expression of *Gzmb* in CD4^+^ TILs while *Ifnar^-/-^* CD4^+^ T cells also exhibited reduced *Gzmb* expression, demonstrating that detection of type-I IFN by CD4^+^ T cells is necessary for their differentiation into poised T_CTX_. Our data suggest that type-I IFN mediates this effect by inducing *Prdm1* expression, consistent with in vitro observations^49^.

We have previously shown that Blimp-1 is necessary for acquisition of cytotoxic activity in CD4^+^ T cells in response to anti-CTLA-4 treatment^7^. We provide new insight here by demonstrating that this reflects a role for Blimp-1 in establishing the poised state. Furthermore, we show that differentiation into poised CD4^+^ T_CTX_ is antagonized by Bcl6, consistent with previous findings from viral infection models^19^. Our data suggest that Bcl6 does not specifically antagonize T_CTX_ differentiation but rather inhibits multiple CD4^+^ T cell differentiation pathways in tumors. This is reminiscent of the role of the Blimp-1-Bcl6 axis in regulating proliferation versus terminal differentiation of CD8^+^ T cells and B cells^23^.

We demonstrate that the block to GzmB protein production is mediated by the RBPs ZFP36 and ZFP36L1. The ZFP36 family mediates post-transcriptional regulation of T cell effector function in a number of contexts by modulating mRNA stability and translation^27,28,29,31,36^. ZFP36 family members directly regulate a set of effector proteins, including IFNγ, TNFα and IL-2, by binding to AU-rich elements in the 3’UTR of their mRNAs^30,31,32,33,36^. Our observation that GzmB protein production by tumor-infiltrating CD4^+^ T_CTX_ in DKO mice is independent of IL-2 shows that de-repression of IL-2 production is not responsible for the phenotype. Similarly, although ZFP36 family members also function in T_REG_^50^, the IL-2-independent nature of GzmB upregulation in DKO CD4^+^ T_CTX_ rules out defects n T_REG_-mediated IL-2 depletion as the cause.

In addition to effector cytokines, ZFP36 family members bind a wide range of additional transcripts in CD8^+^ T cells to regulate multiple aspects of T cell biology including metabolism, proliferation and dependence on co-stimulation^31,32,33,34^. It is likely that this broad functionality of ZFP36 and ZFP36L1 in CD4^+^ and CD8^+^ T cells, rather than effects on any single protein, underlies the phenotypes we observe in CE and DKO mice but it will be important to determine which proteins are regulated by ZFP36/ZFP36L1 in tumor-infiltrating T cells to further understand the nature of this post-transcriptional regulatory checkpoint.

We show that anti-CTLA-4 and anti-LAG-3 plus anti-PD-1 treatments downregulate *Zfp36l1* and that constitutive expression of *Zfp36l1* blocked the ability of both treatments to upregulate GzmB protein production.. We also show that *Zfp36l1* downregulation is necessary for control of tumor growth. In comparison, the effect of *Zfp36l1* constitutive expression on animal survival in response to anti-LAG-3+anti-PD-1 was more modest. This suggests that the therapeutic effect of these antibodies is also mediated through additional pathways.

*Zfp36* and *Zfp36l1* DKO increased production of the effector molecules GzmB and IFNγ without increasing expression of the exhaustion markers LAG-3, PD-1 or TIM-3. *Zfp36* and *Zfp36l1* DKO also prolonged survival and this was further extended when combined with anti-LAG-3+anti-PD-1 treatment. These results suggest that treatments that inhibit the function of ZFP36/ZFP36L1 or downregulate *Zfp36l1* may hold promise as therapies, particularly in combination with anti-LAG-3+anti-PD-1 or other checkpoint inhibitors.

In summary, our findings demonstrate that CD4^+^ T_CTX_ are present in tumors in a poised state that is subject to a post-transcriptional regulatory checkpoint mediated by ZFP36 and ZFP36L1 that restrains GzmB protein production and controls the response to immunotherapy. This work reveals mechanisms that regulate CD4^+^ T_CTX_ function in tumors and highlights the potential therapeutic utility of targeting post-transcriptional regulatory pathways in T cells for the treatment of cancer.

## Acknowledgements

We thank S. Martin, C. Qing, J. Clancy, K. Foster, Y. Morris, T. Clark, P. Levy and the staff of the UCL Biological Services Units for technical expertise. Thanks to the UCL Cancer Institute Flow Cytometry Translational Technology Platform (TTP) for cell sorting and to the UCL Cancer Institute Genomics TTP and UCL Genomics for sequencing. We thank Manuel Rodriguez-Justo and the UCL/UCLH Biobank for Studying Health and Disease for the provision of human tissue samples. M.V.d.M. was supported by a Cancer Research UK (CRUK) UCL Centre (C416/A18088) PhD studentship. M.V.d.M. and C.L. were funded by a Medical Research Council grant (MR/W002337/1) awarded to R.G.J and S.A.Q.; J.H. was funded by a CRUK City of London Centre [CANCTA-2022/100001] PhD studentship. S.A.Q. was funded by a Cancer Research UK (CRUK) Senior Cancer Research Fellowship (C36463/A22246), and S.A.Q., C.C. and D.K. were funded by the CRUK Biotherapeutic Program grant (C36463/A20764) and Biotherapeutic Program grant (DRCRPGTD-May21\100001). M.N.S was funded by Becas Chile. M.T. was funded by a BBSRC Institute Strategic Programme Grant (BBS/E/B/000C0407). C.N and C.J.T. were funded by the UKRI Medical Research Council (MR/T028270/1), Cancer Research UK (C60693/A23783), the Cancer Research UK City of London Centre (CTRQQR-2021/100004), and the UCLH Biomedical Research Centre (BRC422). Work at the CRUK City of London Centre Single Cell Genomics Facility and UCL Cancer Institute Bioinformatics Hub was supported by the CRUK City of London Centre Award [CTRQQR-2021/100004]. This work was supported by the CRUK City of London Centre Award [CANCTA-2022/100001] and the Cancer Immuno-therapy Accelerator Award (CITA-CRUK) (C33499/A20265).

## Contributions

Conceptualization: M.V.d.M, S.A.Q, R.G.J. Data curation: M.V.d.M Formal analysis: M.V.d.M, J.H, C.L, C.N. Funding acquisition: M.T, S.A.Q, R.G.J. Investigation: M.V.d.M, C.C., J.H., C.L, A.S, I.U., M.N.S, M.S, G.M, D.K., C.N. Project administration: M.V.d.M, S.A.Q, R.G.J. Resources: S.E.B., M.T. Supervision: C.J.T., M.T., S.A.Q, R.G.J. Validation: M.V.d.M, J.H, C.L, A.S, C.C. Writing – original draft: M.V.d.M, S.A.Q, R.G.J. Writing – review & editing: M.V.d.M, J.H, C.C., C.L, M.T, S.A.Q, R.G.J. Ethics Declaration M.T. has a funded collaboration with AZ on a topic unrelated to this study. The remaining authors declare no competing interests.

## Methods

### Mice

C57BL/6J and *Bcl6*^fl/fl^ mice^51^ were purchased from Charles River Laboratories. *Ifnar^-/-^* mice were purchased from The Jackson Laboratory. *Cd4*^Cre^ mice were a kind gift from B. Seddon (UCL), *Prdm1*^fl/fl^ mice^52^ from T. Korn (TUM, Munich, Germany) and *Rag1^-/-^* mice from G. Kassiotis (Francis Crick Institute, London, UK). *Zfp36*^fl/fl^ *Zfp36l1*^fl/fl^ mice were described in Petkau et al.^31^. *GFPZfp36l1^tg/tg^* mice were described in Vogel et al.^42^. All transgenic mice were of C57BL/6 background and bred in Charles River Laboratories or University College London (UCL) BSUs. Mice were 5-10 weeks old and age and sex-matched within experiments unless otherwise stated. All animal studies were performed under UCL and UK Home Office ethical approval and regulations.

### Cell lines

MCA205 sarcoma tumor cells were cultured in complete DMEM supplemented with 2 mM GlutaMAXTM, 10% FBS, 100 U/mL penicillin and 100 μg/mL streptomycin (all from ThermoFisher). MC38 cells were cultured in complete RPMI-1640 (cRPMI, ThermoFisher) media supplemented as above.

### Patient samples

Fresh tumor samples and peripheral blood from newly diagnosed patients were collected from University College London Hospital (UCLH) via the UCL/UCLH Biobank, reference 20/YH/0088. Adult patients with cancer of the gastrointestinal tract, including patients with cancer of unknown primary, that attend cancer clinicals as well as screening clinics at UCL Partners were eligible to participate in the study. Written and informed consent, as regulated by the UCL/UCLH Biobank Research Ethics Committee, was obtained from all patients included in this study.

### Mouse tumor models

C57BL/6J WT or transgenic mice were injected subcutaneously with 0.5×10^6^ MCA205 or 0.5×10^6^ MC38 cells in PBS. Mice received 100 μg anti-CTLA-4 clone 4F10 mouse IgG2a or 4F10 mouse IgG1 (Evitria); 100 μg anti-PD-1 rat IgG2a (RMP1-14; BioXcell); 200 μg anti-LAG-3 rat IgG1 (C9B7W; BioXcell); 200 μg anti-IL-2 rat IgG2a (JES6-1A12; BioXcell); 200 μg anti-IFNAR mouse IgG1 (MAR15A3; BioXcell) by intraperitoneal injection. When measuring protein levels by flow cytometry, mice received 3 doses of the corresponding antibodies on days 6, 9 and 11 post-tumor inoculation and were sacrificed on day 12. When measuring RNA levels by scRNA-seq or RT-qPCR, mice received 2 doses of treatment on days 6 and 9, and were sacrificed on day 10. For tumor growth experiments, mice received 3 doses of the corresponding antibodies on days 6, 9 and 12 post-tumor inoculation and mice were sacrificed when any orthogonal tumor diameter reached 15mm, or if tumors ulcerated. Tumors volumes were calculated as 4/3π*abc*, where *a*, *b*, and *c* are radii.

### Adoptive CD4^+^ T cell transfer

*Rag1^-/-^* mice were inoculated subcutaneously with 0.5×10^6^ MAC205 cells. At day 3, CD4^+^ T cells were isolated from LNs of WT or *Ifnar^-/-^* mice by magnetic separation using a mouse CD4+ T Cell Isolation Kit (Miltenyi Biotec) according to the manufacturer’s instructions. 0.5×10^5^ CD4^+^ T cells from WT or *Ifnar^-/-^* mice were injected intravenously into MCA205 tumor-bearing *Rag1^-/-^* mice. Mice were sacrificed on day 10.

### Mouse tissue processing

Tumor draining lymph nodes and tumors were dissected into cRPMI medium (ThermoFisher). Lymph nodes were strained through a 70 μm mesh using RPMI, pelleted and re-suspended in cRPMI for downstream analysis. Tumors were mechanically disrupted using scissors, followed by digestion with 0.33 mg/mL Liberase TL (Roche) and 0.2 mg/mL DNase (Roche) in serum-free RPMI at 37°C for 30 min. Tumors were then dispersed through a 70 μm filter in PBS + 2 mM EDTA and cells resuspended in cRPMI for downstream processing and analysis. Lymphocytes were enriched from tumor samples by centrifugation using a Histopaque 1119 density gradient (Sigma-Aldrich), followed by flow cytometry staining or *ex vivo* re-stimulation with phorbol 12-myristate 13-acetate (PMA, 20 ng/mL) and ionomycin (500 ng/mL) for 4 hours at 37°C in the presence of GolgiPlug^TM^ (containing brefeldin A, BD Biosciences). For samples requiring fluorescence activated cell sorting (FACS), T cells were enriched using Dynabead FlowComp Pan-T CD90.2 beads (Invitrogen) according to the manufacturer’s instructions prior to staining.

### *In vitro* murine T cell activation assays

Naïve CD4^+^ T cells were isolated from the lymph nodes and spleen of either CE, DKO, or littermate control animals, as indicated, by magnetic separation using the naïve CD4^+^ T cell isolation kit (Miltenyi Biotec), as per the manufacturer’s instructions. Purified T cells were stimulated and cultured with anti-CD3 (145-2C11, BioXCell; 0.2 or 1 ug/mL) and anti-CD28 (37.51, BioXCell; 0.5 ug/mL). Cells were harvested after 4 days and stained with directly conjugated antibodies.

### Human tissue processing

Human TILs were isolated from tumor tissue samples no later than 24 hours after collection. Tissue was cut into 1-2 mm^3^ fragments and processed using gentleMACS^TM^ Octo Dissociator at 37°C for 60 minutes in serum-free RPMI-1640 containing 30 μg/mL Collagenase Type I (ThermoFisher) and 75 μg/mL DNase I (Roche/Sigma). The homogenate was passed through a 70 μm cell strainer and lymphocytes were isolated with Ficoll-Paque as above. Single-cell suspensions were cryopreserved and thawed as above.

### Human blood processing

Human PBMCs were isolated no later than 24 hours after collection. Patient blood was mixed with serum-free RPMI-1640 at a 1:1 ratio and lymphocytes enriched by gradient centrifugation using Ficoll-Paque Plus (VWR). PBMCs were cryopreserved in FBS + 10% DMSO prior to further use. Samples were thawed in RPMI-1640 media supplemented with 20% FBS, 2 mM GlutaMAX^TM^ 100 U/mL penicillin, 100 μg/mL streptomycin (all from ThermoFisher) and 35 ng/mL DNase I (Roche) prior to staining for FACS or flow cytometry.

### Fluorescently activated cell sorting (FACS)

Cells were stained for cell surface markers as above and resuspended in PBS with 1% BSA and 2 mM EDTA, using 1 μg/mL DAPI (BD Biosciences) as a viability dye. Cells were filtered and sorted using a FACSAria^TM^ Fusion or FACS Aria III^TM^ (BD Biosciences). For mouse tumor experiments, cells from 3-6 mice were pooled per sample prior to staining and sorting, apart from experiments in *Rag1^-/-^* mice, in which each tumor sample was sorted individually. Antibodies used are listed in Supplementary Table 6.

### Flow cytometry

Directly conjugated antibodies employed for flow cytometry are listed in Supplementary Table 6. Cells were stained using directly conjugated antibodies against extracellular antigens diluted in PBS with 1% BSA and 2 mM EDTA containing Mouse BD Fc Block™ (BD Biosciences) or Human BD Fc Block™ (BD Biosciences), and Fixable viability dye eFluor780 (ThermoFisher) for 30 min at 4°C. Intracellular antigens were stained using the FoxP3 staining kit (eBioscience) according to the manufacturer’s instructions. For quantification of absolute cell numbers, a defined number of CountBright^TM^ Absolute counting Beads (Invitrogen) was added to the sample. Data were acquired using a BD LSR-^FortessaTM^ X-20 or ^FACSymphonyTM^ X30 (both BD Biosciences) and analyzed using FlowJo v.10.8.1/v.10.10.0 software (TreeStar). Common gating strategies can be found in Supplementary Figures 1 and 2.

### CyTOF

#### Sample preparation

Inguinal tumor draining lymph nodes and tumors were dissected for each animal individually to allow rapid fixation. Tumors were cut into 2-3 mm fragments using a scalpel on a glass petri dish on ice before being placed in ice-cold fixation buffer (4% PFA/PBS; Cambridge Bioscience). Lymph nodes were placed directly into fixation buffer. Tissues were fixed at 4°C for 4 hrs, followed by digestion with 0.4 mg/mL Liberase TL (Roche) and 0.2 mg/mL DNase (Roche) in serum-free RPMI at 37°C for 30 min. Both lymph nodes and tumors were dispersed through a 70 μm filter in PBS + 2 mM EDTA.

#### Sample staining

5 million cells were pelleted into each well of a round-bottom 96-well plate (Thermo 163320) before labelling with thiol-reactive organoid barcoding in situ (TOBis) barcodes, as described^53^. Pellets were resuspended in 200 μL of 126-plex (9-choose-4) TOBis barcode overnight at 4°C. Unbound barcodes were quenched in 200 μL of 2mM L-Glutathione (GSH) three times. Cells were then washed twice in PBS before pooling barcoded samples into a single staining tube. Twelve million pooled cells were washed in cell staining buffer (CSB) (Standard BioTools 201068) and stained with rare-earth metal-conjugated antibodies (conjugated in-house as described^53,54^; Supplementary Table 7) against extracellular antigens for 30 min at room temperature. Cells were then permeabilised in 0.1% (v/v) Triton X-100/PBS (Sigma T8787) for 1 hr, washed with CSB, and stained with anti-FoxP3 antibody for 30 min at room temperature. Cells were then permeabilised with 50% methanol/PBS (Fisher 10675112) and stained with rare-earth metal conjugated antibodies (Key Resources Table) against intracellular antigens for 30 min at room temperature. Cells were washed, resuspended in cell acquisition solution plus (CAS+) (Standard BioTools 201244) with 2 mM EDTA (Sigma 03690), and analysed using a CyTOF XT (Standard BioTools, SCR\_02634) at 200–400 events s-1.

#### Data processing

Multiplexed FCS files were debarcoded into individual experimental conditions using the Zunder Lab Single Cell Debarcoder tool (https://github.com/zunderlab/single-cell-debarcoder)^55^. Debarcoded FCS files were uploaded to Cytobank and gated for Gaussian parameters and DNA content (Ir191/193). CD4^+^ T_EFF_ were gated as CD45^+^ CD3^+^ CD4^+^ CD8α^-^FoxP3^-^, and Earth Mover’s Distance (EMD) scores for each marker calculated using the Python package scprep. Standard EMD scores were calculated between probability distributions (markers) across experimental conditions using defined controls as references. Signs were applied to EMD scores to denote an increase or decrease in expression.

### Reverse transcription quantitative PCR (RT-qPCR)

RNA from FACS-purified cells was extracted using RNeasy micro kit (Qiagen) according to the manufacturer’s instructions, including the in-column DNase I digestion step. RNA was quantified using the Qubit High sensitivity RNA assay (ThermoFisher). cDNA was synthesized using Superscript III Reverse Transcriptase (ThermoFisher) and random primers according to the manufacturer’s instruction. qPCR analysis was carried out using QuantiTect SYBR Green PCR Kit (Qiagen) with gene specific primers (Supplementary Table 8). Gene expression levels were calculated relative to *Hprt1* in mouse samples and *ACTB* in human samples.

### Single-cell RNA sequencing (scRNA-seq)

scRNA-seq was carried out using the using the Chromium Single Cell 5’ V(D)J Reagent kit, v1 and v2 (10X Genomics). Live CD3^+^ T cells were sorted and counted manually prior to capture on the 10X Chromium Controller. Purified single cell suspensions were loaded onto the 10X Chromium Controller and libraries prepared as per the manufacturer’s instructions, using AMPure beads for size selection and purification and the Bioanalyzer High sensitivity DNA assay (Agilent) to assess cDNA concentration and size of the final libraries. Library concentrations were quantified by qPCR using the Illumina Library Quantification kit (KAPA). Libraries were pooled at a final concentration of 10 nM for sequencing. Pooled libraries were sequenced using a High Output run of the NextSeq 550 or NovaSeq platforms using 26 x 91 paired-end reads. Samples were sequenced at a minimum depth of 20,000 reads/cell, based on the estimated number of cells loaded onto the Chromium controller.

### scRNA-seq data analysis

Quality control and alignment of sequencing reads was performed using the Cell Ranger analysis pipelines from 10X Genomics (version 3.0.2 and version 7.0.0, mm10 genome assembly). Analysis of aligned mouse scRNA-seq data was performed using Seurat (version 3.1.5)^56^. High quality cells in which 500-3000 genes were detected and contained < 5% of mitochondrial reads were selected for downstream analysis. Counts were normalized and data from each sample were log normalized using default parameters of the NormalizeData function. Highly variable transcripts were identified using the FindVariableFeatures function, employing a variance stabilizing transformation. Normalized data from each sample were integrated using default parameters of the FindIntegrationAnchors and IntegrateData functions. Integrated data were scaled using ScaleData function and principal component analysis performed using RunPCA. The top 30 principal components were used as the input for a uniform manifold approximation and projection (UMAP) and Shared Nearest Neighbor clustering. The most appropriate clustering resolution for each integrated data set was determined using a hierarchical clustering tree, generated using the Clustree package.

CD4^+^ T cells were selected by filtering for normalized expression values of *Cd3e* > 0 and *Cd4* > 0 and any cells expressing *Klrb1c* or *Cd8a* removed. T_REG_ were selected from bulk CD4^+^ T cells by filtering for *Foxp3 > 0.* T_EFF_ were selected from CD4^+^ T cells by filtering for *Foxp3* ≤ 0 and removing cells belonging to a cluster in which the majority of cells expressed *Foxp3* (>0) in order to exclude potential contamination from T_REG_ in which *Foxp3* transcript was not detected due to shallow sequencing depth.

Marker genes for each cluster were identified using the FindAllMarkers function, with only genes detected in a minimum of 25% of cells from each group included in the analysis. Differentially expressed genes were identified using the FindMarkers feature, using the statistical test MAST^57^. Genes with an adjusted p value <0.05 and an average log_2_ fold change (Log_2_FC) > 0.5 were deemed significantly different between samples tested. Volcano plots were generated using the EnhancedVolcano package^58^ and heatmaps generated using log-transformed, average expression values in the ComplexHeatmap package. Differential abundance analysis was carried out using a genewise negative binomial generalized linear models with quasi-likelihood tests in the package edgeR. Average expression of gene modules in each cell were calculated using Seurat’s AddModuleScore function which calculates average expression of genes of interest on a single cell and gives an enrichment score for these genes relative to the expression levels of an equivalent number of randomly selected genes.

## Statistical analysis

Statistical analyses were performed with Prism v9 or v10 (GraphPad Software). P values were calculated using one-way ANOVA with Tukey post-hoc tests, two-way ANOVA with Šídák’s multiple comparisons test, two-tailed paired-t-test or two-tailed unpaired t-test with Welch’s correction when appropriate. Kaplan-Meier curves were analyzed using the log-rank test. Unless otherwise indicated, ns = p > 0.05, * p < 0.05, **p < 0.01, ***p < 0.001, ****p < 0.0001 for all tests. N in animal experiments refers to number of animals per experimental group.

## Data availability

Single-cell RNA-seq data have been deposited at GEO with accession number GSE230230. Any additional information required to reanalyze the data reported in this paper is available from the corresponding authors upon request.

## Reporting Summary

Further information on research design is available in the Nature Research Reporting Summary.

## Supplementary Information

**Supplementary Figure 1.** Gating strategies for mouse samples.

**Supplementary Figure 2.** Gating strategy for human samples.

**Supplementary Table 1.** Marker genes for CD4^+^ T cell clusters (C57BL/6J WT mice).

**Supplementary Table 2.** Genes differentially expressed between tumor-infiltrating T_REG_ from untreated and anti-CTLA-4 treated mice.

**Supplementary Table 3.** Genes differentially expressed between tumor-infiltrating CD4^+^ T_EFF_ from untreated and anti-CTLA-4 treated mice.

**Supplementary Table 4.** Marker genes for CD4^+^ T cell clusters (*Cd4*^Cre^, *Bcl6*^-/-^ and *Prdm1*^-/-^mice).

**Supplementary Table 5.** Genes differentially expressed between tumor-infiltrating CD4^+^ T_EFF_ from *Cd4*^Cre^ and *Bcl6*^-/-^ or *Cd4*^Cre^ and *Prdm1*^-/-^ mice.

**Supplementary Table 6.** Antibodies used for FACS and flow cytometry

**Supplementary Table 7.** Antibodies used for CyTOF

**Supplementary Table 8.** Primers used for RT-qPCR

---

**Extended Data Fig. 1.**
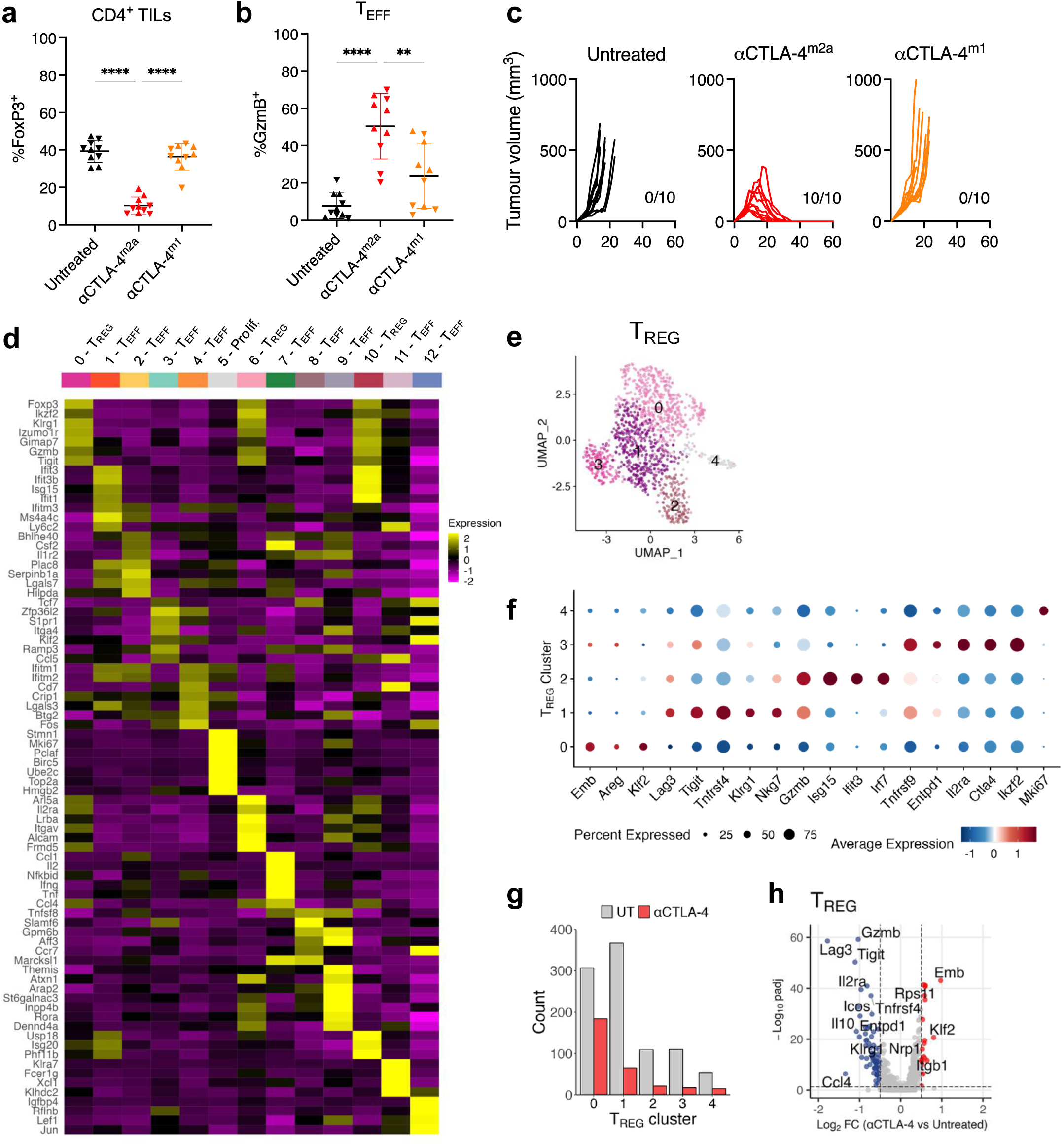
Gene expression in CD4^+^ TILs in untreated and anti-CTLA-4 treated tumors. **a-b,** Proportions of (**a**) FoxP3^+^ CD4^+^ T cells and (**b**) GzmB^+^ CD4^+^ T_EFF_ (pre-gated as FoxP3^-^CD25^-^) in MCA205 tumors treated with anti-CTLA-4^m2a^ (mouse IgG2a) or anti-CTLA-4^m1^ (mouse IgG1) on days 6, 9 and 11 post-tumor inoculation or left untreated. Mice were sacrificed on day 12 post-tumor inoculation (n=10/group from two independent experiments, one-way ANOVA). **c,** MCA205 tumor growth in mice treated as indicated. Number of mice surviving to day 60 post-tumor inoculation is indicated (n=10/group). **d,** Heatmap showing average expression of marker genes for each cluster of CD4^+^ T cells identified in MCA205 tumors from untreated and anti-CTLA-4 treated mice. Relative expression is shown by color according to the scale on the right. **e,** UMAP of MCA205-infiltrating T_REG_ (n=1249; two independent experiments). **f,** Heat map of marker genes for each cluster. Size of the dot indicates percentage of expressing cells and color indicates normalized expression in each cluster. **g,** Number of cells in each T_REG_ cluster in tumors from either treatment condition. **h,** Volcano plot showing genes differentially expressed between tumor-infiltrating T_REG_ from untreated and anti-CTLA-4^m2a^ treated mice (q<0.05, |Log_2_FC|>0.5, MAST).

**Extended Data Fig. 2.**
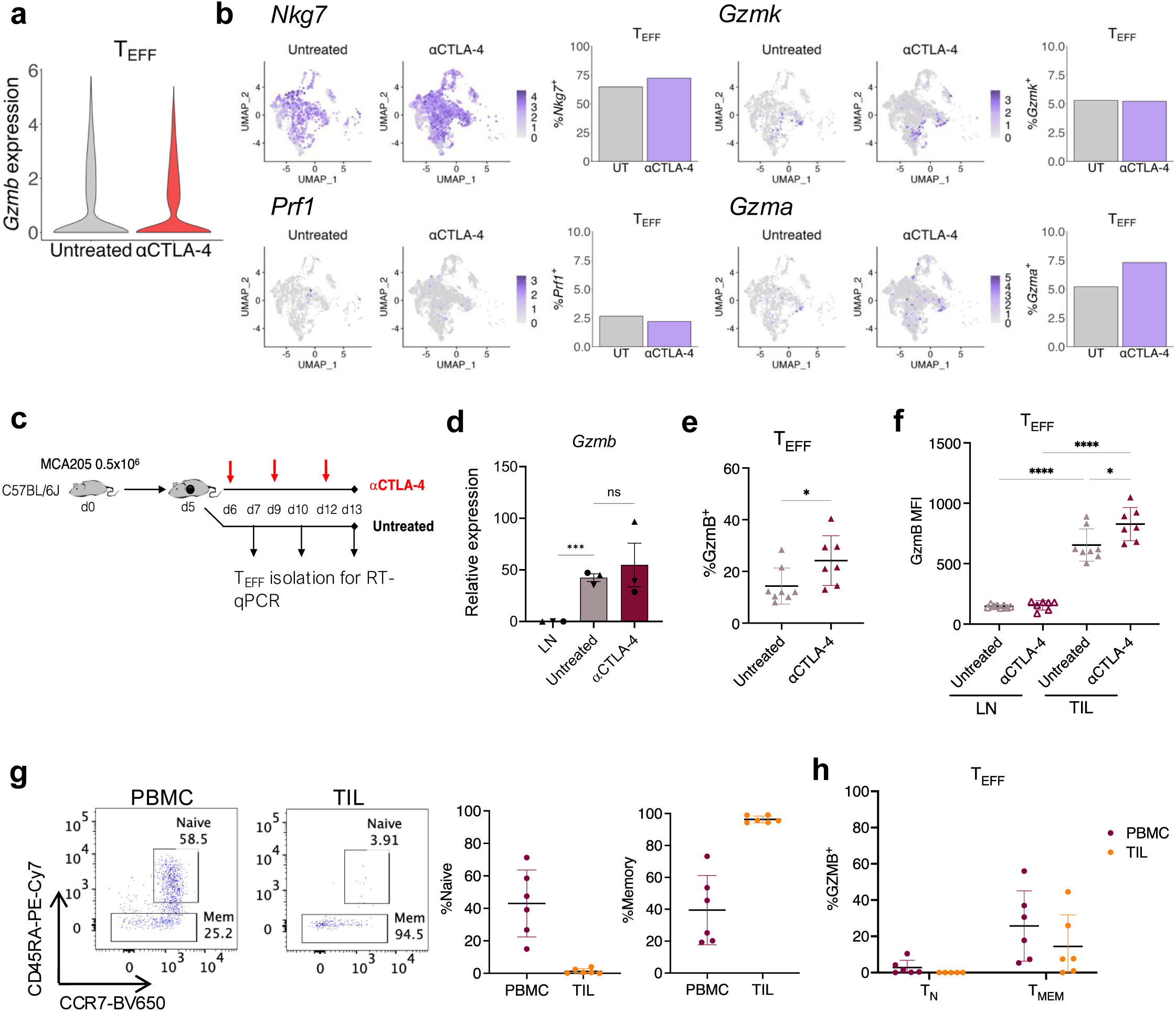
Poised CD4^+^ T_CTX_ are found in untreated tumors. **a,** Violin plot showing *Gzmb* mRNA levels in *Cd4^+^* T_EFF_ from MCA205 tumors from untreated and anti-CTLA-4 treated mice. **b,** Left: UMAP of *Cd4^+^* T_EFF_ from MCA205 tumors from untreated and anti-CTLA-4 treated mice, showing normalized expression of *Nkg7, Prf1, Gzmk* and *Gzma* per cell. Right: Proportions of *Cd4^+^* T_EFF_ expressing each gene in either treatment condition **c,** Schematic of treatment strategy for RT-qPCR time course (Fig. 1g). C57BL/6J mice challenged with MCA205 tumors were treated with anti-CTLA-4 on days 6, 9 and 12 post-tumor inoculation, or left untreated. Mice were sacrificed at days 7, 10 or 13 and CD4^+^ T_EFF_ (CD4^+^CD25^-^) isolated for analysis. **d,** RT-qPCR of *Gzmb* mRNA in T_EFF_ (CD4^+^CD25^-^) from tumor-draining LN or MC38 tumors analyzed at day 10 post-tumor inoculation. Expression is relative to *Hprt1* (mean±SD of 3 independent experiments (each point is the average of 3-5 mice); one-way ANOVA). **e,** Proportions of GzmB^+^ CD4^+^ T_EFF_ in MC38 tumors treated with anti-CTLA-4 on days 6, 9 and 11 post-tumor inoculation or left untreated. Mice were sacrificed on day 12 (n=11/group, 2 independent experiments; one-way ANOVA). **f,** GzmB MFI in CD4^+^ T_EFF_ from LN and MC38 tumors from untreated and anti-CTLA-4 mice (n=7-8/group, one-way ANOVA). **g,** Left: Representative dot plot showing T_N_ and T_MEM_ populations in PBMC and TILs from CRC patients. Right: proportion of T_N_ and T_MEM_ in PBMCs and TILs from each patient (n=6). **h,** Proportion of T_N_ and T_MEM_ from PBMCs and TILs positive for GZMB.

**Extended Data Fig. 3.**
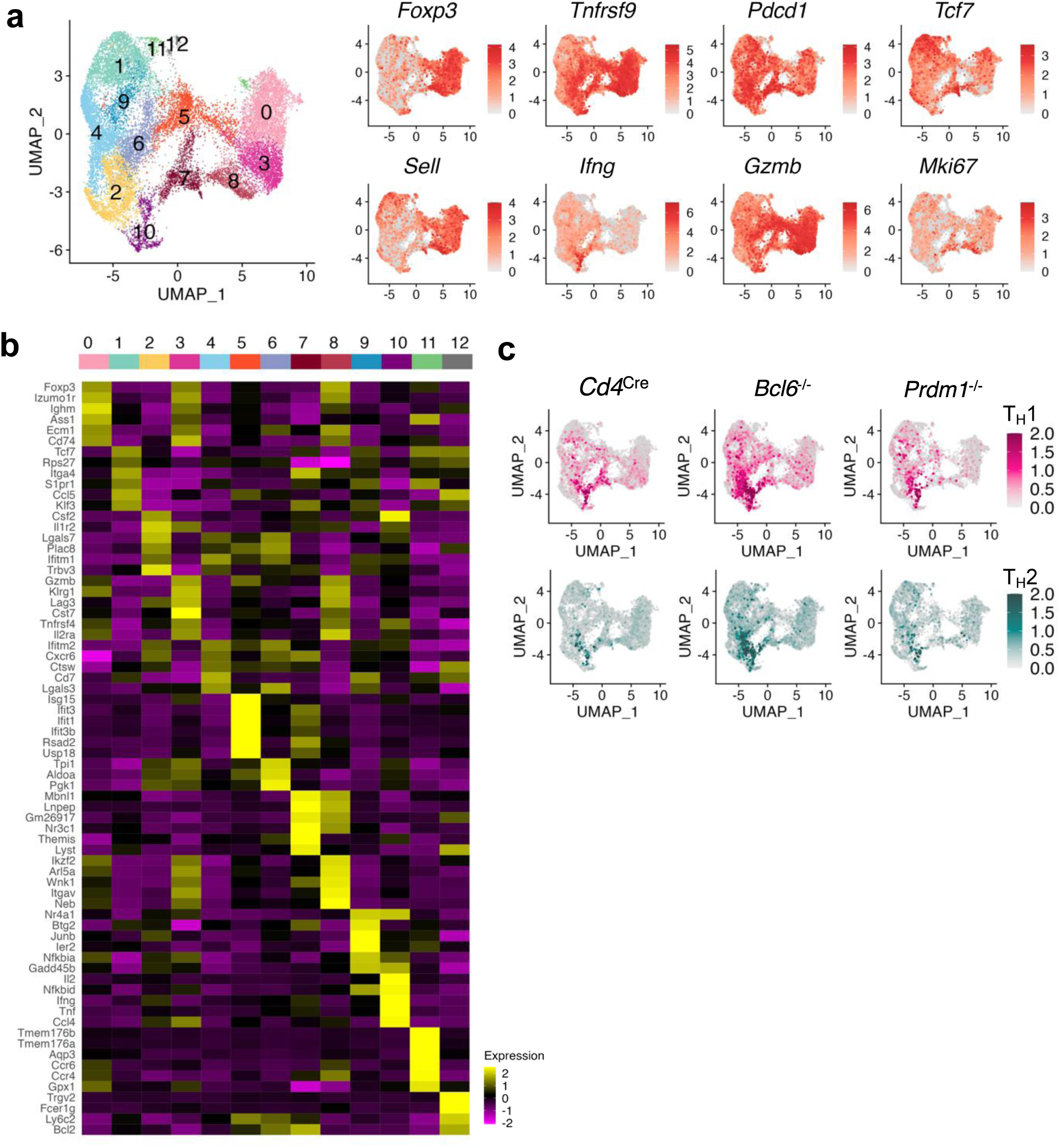
Acquisition of a cytotoxic transcriptional program in poised CD4^+^ T_CTX_ is regulated by the Blimp-1-Bcl6 axis. **a,** UMAP of *Cd4*-expressing T cells from MCA205 tumors from *Cd4*^Cre^, *Bcl6*^-/-^ and *Prdm1*^-/-^ mice showing normalized expression of *Foxp3*, *Tnfrsf9*, *Pdcd1*, *Tcf7*, *Sell*, *Ifng*, *Gzmb* and *Mki67* per cell. **b,** Heatmap showing average expression of marker genes for each cluster of *Cd4^+^* T cells identified in MCA205 tumors from *Cd4*^Cre^, *Bcl6*^-/-^ and *Prdm1*^-/-^ mice. **c,** Average expression of T_H_1 (*Tbx21, Ifng, Hopx, Csf2*) and T_H_2 (*Gata3, Il13, Il5, Il4*) gene signatures in CD4^+^ T cells from MCA205 tumors from *Cd4*^Cre^, *Bcl6*^-/-^ and *Prdm1*^-/-^ mice.

**Extended Data Fig. 4.**
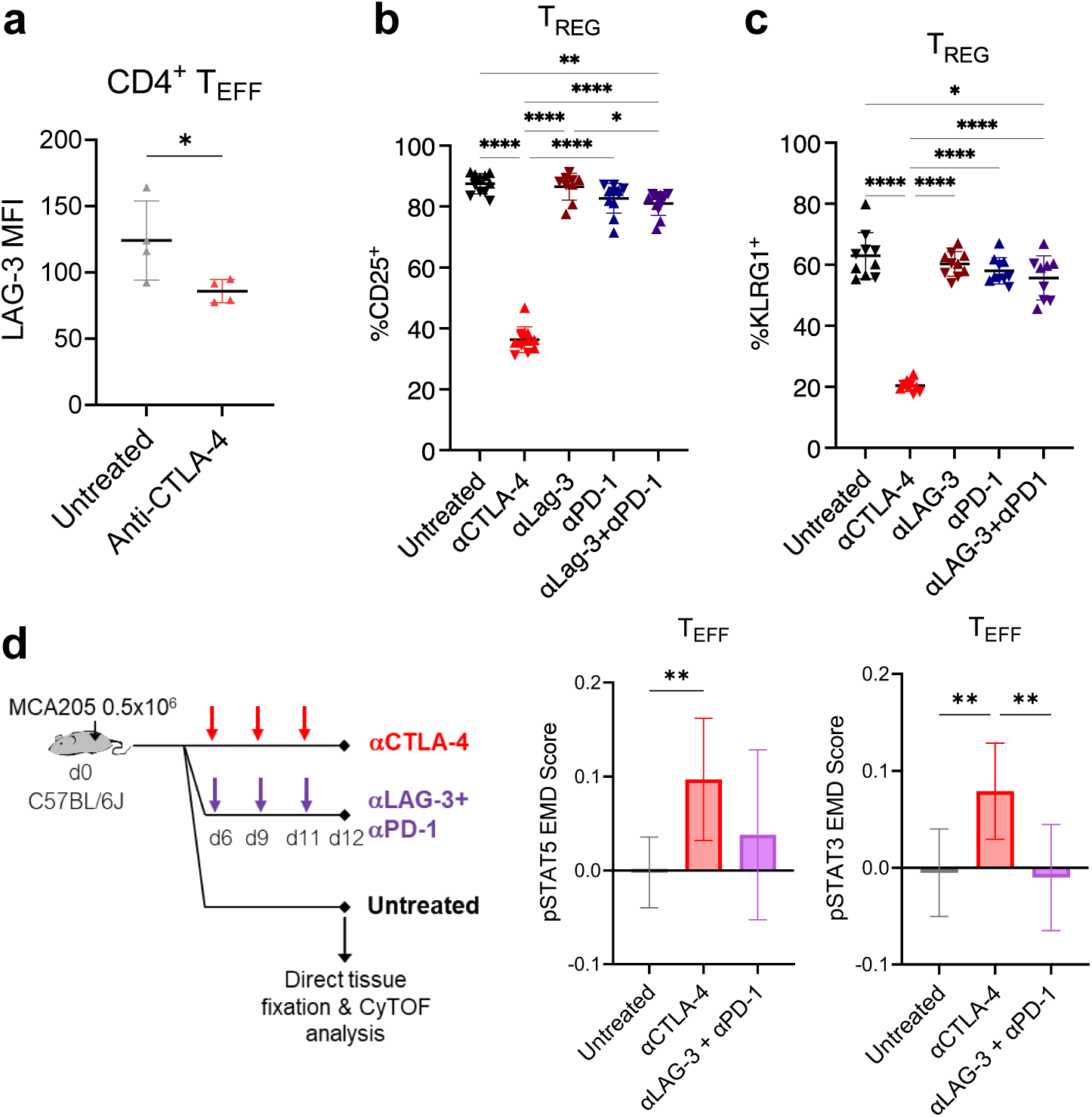
Blockade of LAG-3 and PD-1 induces GzmB protein production in poised CD4^+^ T_CTX_. **a,** LAG-3 MFI in MCA205-infiltrating CD4^+^ T_EFF_ mice treated with anti-CTLA-4 or left untreated (n=4/group from 1 independent experiment, Student’s t-test). **b-c,** Proportion of (**b**) CD25^+^ and (**c**) KLRG1^+^ T_REG_ (pre-gated as CD4^+^FoxP3^+^) from MCA205 tumors from mice treated with anti-CTLA-4, anti-LAG-3, anti-PD-1 or anti-LAG3+anti-PD-1 on days 6, 9 and 11 post-tumor inoculation or left untreated (n=9-11/group from 2 independent experiments, one-way ANOVA). **d,** Left: Schematic of mouse treatments for CyTOF experiment. C57BL/6J MCA205 tumor-bearing mice were treated with indicated antibodies on days 6, 9 and 11 post-tumor inoculation or left untreated and mice were sacrificed and tumors and lymph nodes fixed and analyzed on d12. Right: Earth mover’s distance (EMD) score showing the change in pSTAT5 (left) and pSTAT3 (right) in T_EFF_ from MCA205 tumors from mice treated with anti-CTLA-4 or anti-LAG-3 + anti-PD-1 relative to mice left untreated (n=9-10/group from 2 independent experiments; one-way ANOVA).

**Extended Data Fig. 5.**
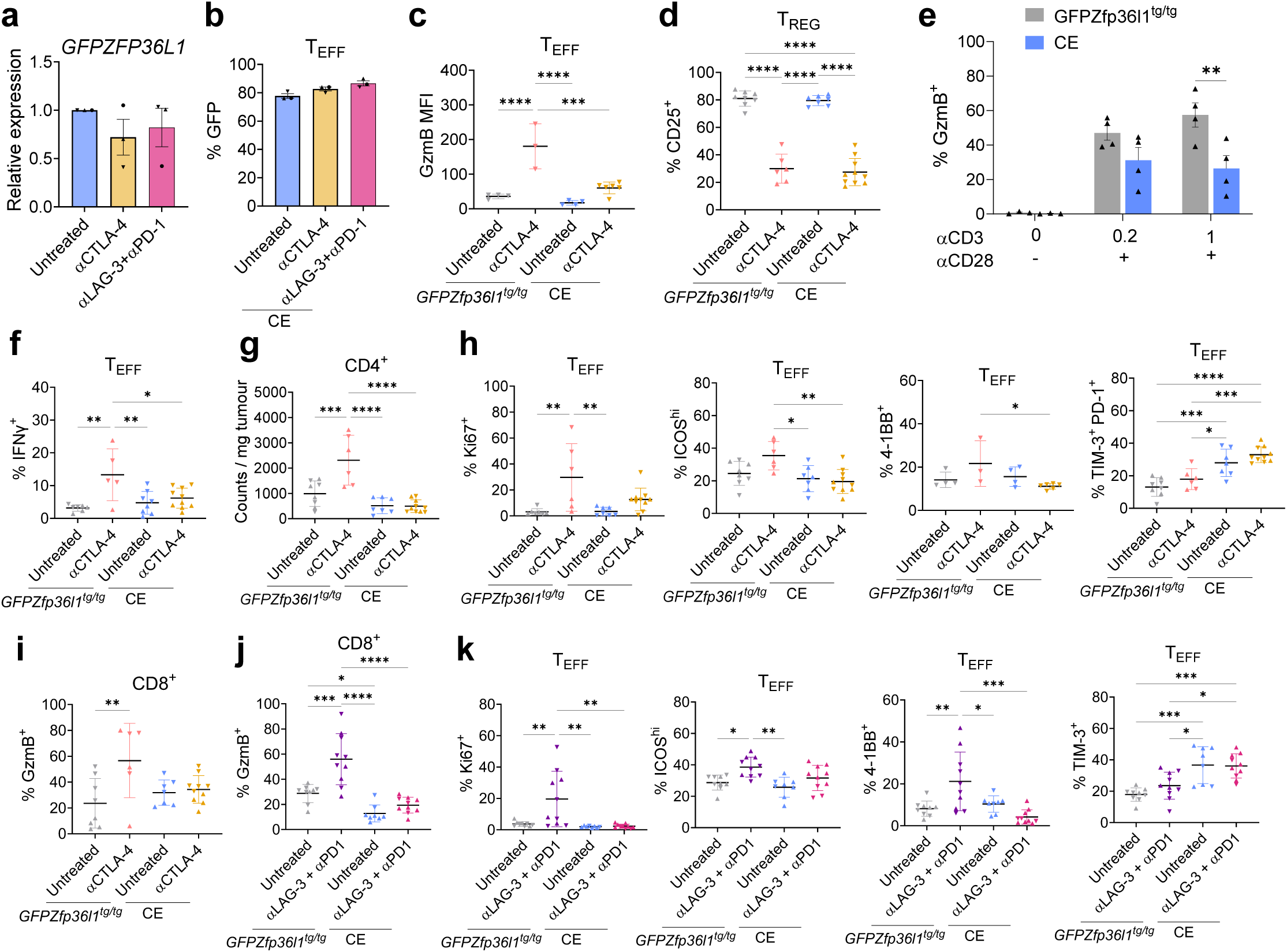
ZFP36L1 blocks GzmB protein production in tumor-infiltrating CD4^+^ T_EFF_. **a,** RT-qPCR analysis of *GFPZFP36L1* mRNA in CD4^+^ T_EFF_ from MCA205 tumors from *Cd4^Cre^ GFPZfp36l1*^tg/tg^ (CE) mice treated with anti-CTLA-4 or anti-LAG-3 + anti-PD-1 on days 6 and 9 post-tumor inoculation, or left untreated, and sacrificed on day 10. Expression is relative to *Hprt1* (mean±SD of 3 independent experiments (each point is the average of 4-6 mice); one-way ANOVA). **b,** Proportion of GFP^+^ CD4^+^ T_EFF_ from MCA205 tumors from CE mice treated as in a (mean±SD of 3 independent experiments (each point is the average of 4-6 mice); one-way ANOVA) **c,** Mean fluorescence intensity of GzmB in T_EFF_ from MCA205 tumors from CE mice or littermate controls treated with anti-CTLA-4 on days 6, 9 and 11 post-tumor inoculation, or left untreated, and sacrificed on day 12 (n=3-6 mice/group; one-way ANOVA). **d,** Proportion of CD25^+^ T_REG_ from MCA205 tumors from untreated or anti-CTLA-4 treated CE animals or littermate controls sacrificed on day 12 post-tumor inoculation (n=6-10 mice/group from two independent experiments; one-way ANOVA). **e,** Proportion of GzmB^+^ CE and littermate control CD4^+^ T cells isolated from spleen and LN and left unstimulated or stimulated with anti-CD3 and anti-CD28 (n=3-4/group, two-way ANOVA). **f,** Proportion of IFNγ^+^ T_EFF_ from MCA205 tumors from untreated or anti-CTLA-4 treated littermate or CE animals following *ex vivo* restimulation with PMA/ionomycin (n=5-10 mice/group from two independent experiments; one-way ANOVA). **g,** Number of CD4^+^ T cells per mg of MCA205 tumor from untreated and anti-CTLA-4 treated CE animals or littermate controls sacrificed on day 12 post-tumor inoculation (n=6-10 mice/group from two independent experiments; one-way ANOVA). **h,** Proportion of Ki67^+^, ICOS^hi^, 4-1BB^+^ and TIM-3^+^PD-1^+^ CD4^+^ T_EFF_ from MCA205 tumors from untreated or anti-CTLA-4 treated littermate or CE animals (n=6-10 mice/group from two independent experiments (except 3-6 mice/group from one experiment for 4-1BB); one-way ANOVA). **i,** Proportion of GzmB^+^ CD8^+^ T cells from untreated and anti-CTLA-4 treated CE animals or littermate controls sacrificed on day 12 post tumor inoculation (n=6-10 mice/group from two independent experiments; one-way ANOVA). **j,** Proportion of GzmB^+^ CD8^+^ T cells from MCA205 tumors from untreated and anti-LAG-3+anti-PD-1-treated CE animals or littermate controls (n=8-10 mice/group from two independent experiments; one-way ANOVA). **k,** Proportion of Ki67^+^, ICOS^hi^, 4-1BB^+^ or TIM-3^+^ CD4^+^ T_EFF_ from MCA205 tumors from untreated and anti-LAG-3+anti-PD-1-treated CE animals or littermate controls (n=8-10 mice/group from two independent experiments; one-way ANOVA).

**Extended Data Fig. 6.**
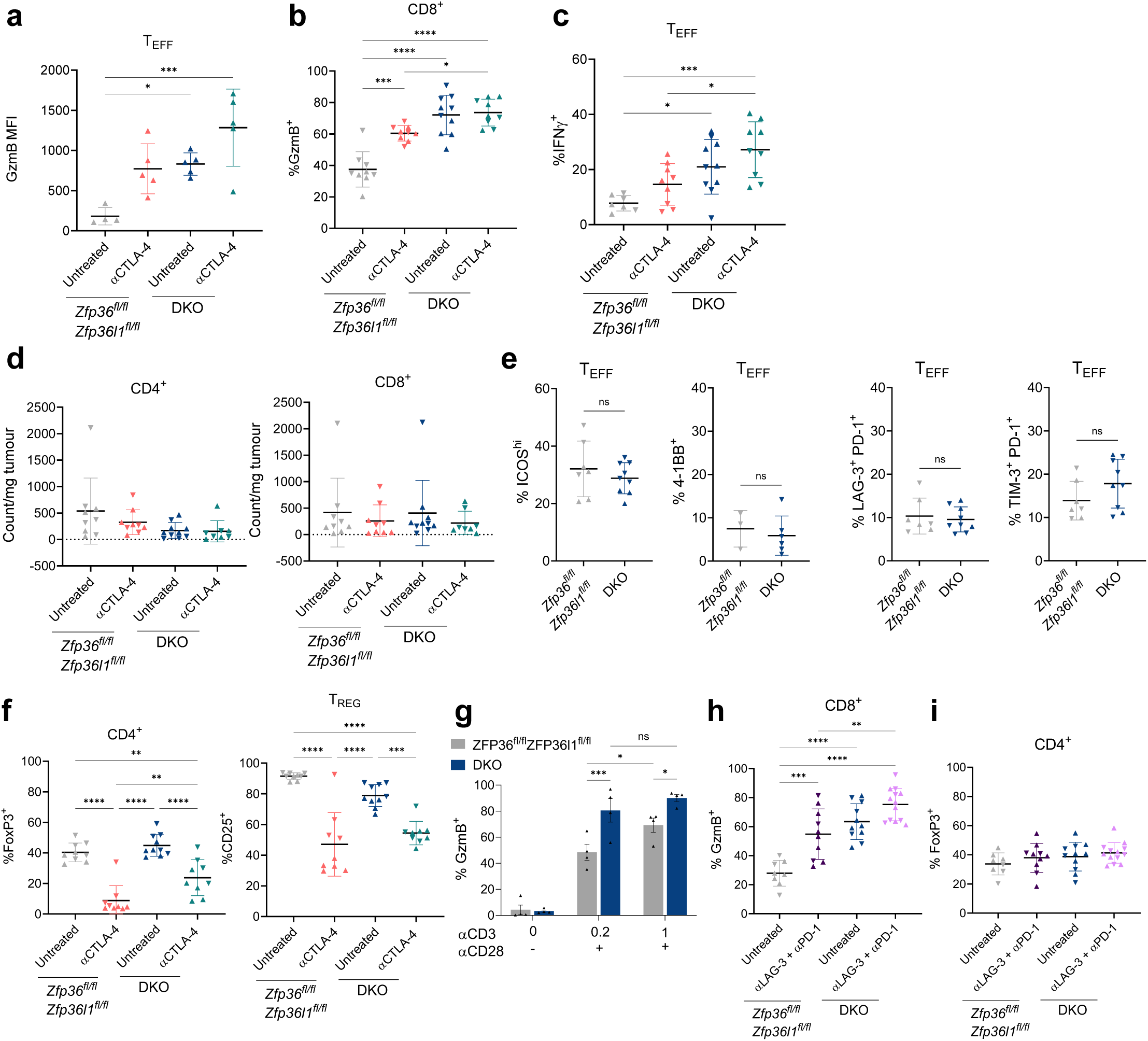
Targeting ZFP36 and ZFP36L1 increases GzmB protein production in tumor-infiltrating CD4^+^ T_EFF_. **a)** Mean fluorescence intensity of GzmB in T_EFF_ from MCA205 tumors from untreated or anti-CTLA-4 treated littermate or DKO animals (n=4-5 mice/group; one-way ANOVA). **b)** Proportion of CD8^+^ T cells positive for GzmB in MCA205 tumors from untreated or anti-CTLA-4 treated littermate or DKO animals (n=9-10 mice/group from 2 independent experiments; one-way ANOVA). **c)** Proportion of IFNγ^+^ T_EFF_ from MCA205 tumors from untreated or anti-CTLA-4 treated littermate or DKO animals following *ex vivo* restimulation with PMA/ionomycin (n=9-10 mice/group from two independent experiments; one-way ANOVA). **d)** Number of CD4^+^ (left) and CD8^+^ (right) cells per mg of MCA205 tumor from untreated and anti-CTLA-4 treated DKO animals or littermate controls. **e)** Proportion of ICOS^hi^, 4-1BB^+^, LAG-3^+^PD-1^+^ and TIM-3^+^PD-1^+^ CD4^+^ T_EFF_ in MCA205 tumors from littermate or DKO animals (n=6-9 mice/group from two independent experiments (except 3-6 mice from one experiment for 4-1BB); Student’s t-test). **f)** Proportion of FoxP3^+^ CD4^+^ T cells (left) and proportion of CD25^+^ T_REG_ (right) from MCA205 tumors from untreated or anti-CTLA-4 treated littermate or DKO animals (n=9-10 mice/group, two independent experiments; one-way ANOVA). **g)** Proportion of GzmB^+^ DKO and littermate control CD4^+^ T cells isolated from spleen and LN and left unstimulated or stimulated with anti-CD3 (0.2 µg/mL or 1 µg/mL) and anti-CD28 (n=3-4/group, two-way ANOVA). **h)** Proportion of CD8^+^ T cells positive for GzmB in MCA205 tumors from untreated or anti-LAG-3+anti-PD-1 treated littermate or DKO animals (n=8-13 mice/group from 2 independent experiments; one-way ANOVA). **i)** Proportion of FoxP3^+^ CD4^+^ T cells in MCA205 tumors from untreated or anti-LAG-3+anti-PD-1 treated littermate or DKO animals (n=8-13 mice/group from 2 independent experiments; one-way ANOVA).

