## Supplementary Tables 1-5 for "A post-transcriptional regulatory checkpoint controls the response of tumor-infiltrating cytotoxic CD4^+^ T cells to immunotherapy"

| | |
|---|---|
| **Supplementary Table 1** | Marker genes for CD4+ T cell clusters (C57BL/6J WT mice). |
| **Supplementary Table 2** | Genes differentially expressed between tumor-infiltrating TREG from untreated and anti-CTLA-4 treated mice. |
| **Supplementary Table 3** | Genes differentially expressed between tumor-infiltrating TEFF from untreated and anti-CTLA-4 treated mice. |
| **Supplementary Table 4** | Marker genes for CD4+ T cell clusters (Cd4Cre, Bcl6-/- and Prdm1-/- mice). |
| **Supplementary Table 5** | Genes differentially expressed between tumor-infiltrating TEFF from Cd4Cre and Bcl6-/- or Cd4Cre and Prdm1-/- mice. |

**Supplementary Table 1.** Marker genes for CD4+ T cell clusters (C57BL/6J WT mice)

| p_val | avg_logFC | pct.1 | pct.2 | p_val_adj | cluster | gene |
|---|---|---|---|---|---|---|
| 0 | 1.52915828 | 0.741 | 0.089 | 0 | 0 | Foxp3 |
| 0 | 1.0907645 | 0.922 | 0.269 | 0 | 0 | Ikzf2 |
| 5.04E-268 | 1.2785519 | 0.5 | 0.081 | 9.28E-264 | 0 | Klrg1 |
| 2.09E-249 | 1.1363222 | 0.617 | 0.154 | 3.85E-245 | 0 | Izumo1r |
| 1.25E-248 | 1.16623089 | 0.787 | 0.34 | 2.30E-244 | 0 | Gimap7 |
| 1.23E-246 | 1.07157908 | 0.792 | 0.323 | 2.27E-242 | 0 | Il2ra |
| 9.09E-221 | 1.3428899 | 0.843 | 0.461 | 1.67E-216 | 0 | Gzmb |
| 5.38E-220 | 1.0151958 | 0.959 | 0.731 | 9.90E-216 | 0 | Tnfrsf4 |
| 1.35E-199 | 0.93894142 | 0.641 | 0.22 | 2.48E-195 | 0 | Cd27 |
| 4.53E-189 | 1.1119303 | 0.77 | 0.411 | 8.33E-185 | 0 | Tigit |
| 2.15E-164 | 1.03192703 | 0.771 | 0.446 | 3.96E-160 | 0 | Pglyrp1 |
| 2.64E-144 | 0.97142448 | 0.778 | 0.499 | 4.85E-140 | 0 | Glrx |
| 3.74E-72 | 0.93981693 | 0.449 | 0.22 | 6.88E-68 | 0 | Lag3 |
| 4.45E-66 | 1.08309774 | 0.269 | 0.087 | 8.18E-62 | 0 | Areg |
| 9.36E-33 | 1.01471763 | 0.301 | 0.161 | 1.72E-28 | 0 | Ccl4 |
| 5.16E-239 | 1.46559517 | 0.611 | 0.155 | 9.50E-235 | 1 | Ifit3 |
| 1.17E-191 | 0.95528555 | 0.399 | 0.066 | 2.15E-187 | 1 | Ifit3b |
| 2.43E-190 | 1.29790051 | 0.842 | 0.493 | 4.48E-186 | 1 | Isg15 |
| 3.88E-185 | 1.13765719 | 0.609 | 0.188 | 7.15E-181 | 1 | Ifit1 |
| 4.68E-171 | 0.9249257 | 0.829 | 0.449 | 8.61E-167 | 1 | Slfn1 |
| 1.54E-154 | 0.87502757 | 0.89 | 0.609 | 2.83E-150 | 1 | Bst2 |
| 3.31E-151 | 1.21710227 | 0.807 | 0.524 | 6.09E-147 | 1 | Ifitm3 |
| 1.38E-144 | 0.84475785 | 0.807 | 0.443 | 2.53E-140 | 1 | Isg20 |
| 2.06E-135 | 0.96361108 | 0.545 | 0.201 | 3.79E-131 | 1 | Ms4a4c |
| 3.88E-131 | 0.89823001 | 0.722 | 0.371 | 7.13E-127 | 1 | Irf7 |
| 2.30E-123 | 0.83024322 | 0.303 | 0.059 | 4.23E-119 | 1 | Mx1 |
| 1.73E-93 | 0.85859391 | 0.724 | 0.432 | 3.19E-89 | 1 | Ifitm2 |
| 5.50E-86 | 0.86668291 | 0.799 | 0.545 | 1.01E-81 | 1 | Rgs1 |
| 1.40E-85 | 1.1839001 | 0.506 | 0.236 | 2.58E-81 | 1 | Ly6c2 |
| 9.38E-63 | 0.85446285 | 0.568 | 0.311 | 1.73E-58 | 1 | Ifitm1 |
| 6.01E-196 | 0.83145094 | 0.949 | 0.517 | 1.11E-191 | 2 | Bhlhe40 |
| 3.74E-140 | 0.67493176 | 0.945 | 0.565 | 6.88E-136 | 2 | Cxcr6 |
| 7.97E-124 | 1.05507926 | 0.338 | 0.072 | 1.47E-119 | 2 | Csf2 |
| 1.50E-105 | 0.85335671 | 0.411 | 0.125 | 2.76E-101 | 2 | Il1r2 |
| 1.47E-100 | 0.76885209 | 0.538 | 0.214 | 2.71E-96 | 2 | Klrc1 |
| 1.74E-96 | 0.8889766 | 0.667 | 0.356 | 3.20E-92 | 2 | Plac8 |
| 3.73E-95 | 0.83473854 | 0.364 | 0.106 | 6.87E-91 | 2 | Serpinb1a |
| 1.19E-92 | 0.66979905 | 0.457 | 0.169 | 2.19E-88 | 2 | AA467197 |
| 2.28E-90 | 0.8269832 | 0.407 | 0.139 | 4.19E-86 | 2 | Lgals7 |
| 1.88E-88 | 0.85444655 | 0.631 | 0.327 | 3.46E-84 | 2 | Hilpda |
| 3.62E-86 | 0.690844 | 0.608 | 0.298 | 6.66E-82 | 2 | Serpinb6b |
| 1.40E-81 | 0.70368501 | 0.611 | 0.312 | 2.58E-77 | 2 | Pdcd1 |
| 4.36E-71 | 0.81292806 | 0.592 | 0.306 | 8.01E-67 | 2 | Ifitm1 |
| 3.72E-67 | 0.68883022 | 0.665 | 0.388 | 6.85E-63 | 2 | Dusp2 |

| | | | | | | |
|---|---|---|---|---|---|---|
| 1.12E-62 | 0.67735989 | 0.323 | 0.116 | 2.06E-58 | 2 | Rgs16 |
| 5.44E-103 | 0.52782047 | 0.983 | 0.946 | 1.00E-98 | 3 | Rps27 |
| 5.33E-102 | 0.87153835 | 0.471 | 0.158 | 9.81E-98 | 3 | Tcf7 |
| 1.10E-82 | 0.87273325 | 0.757 | 0.532 | 2.03E-78 | 3 | Zfp36l2 |
| 9.65E-80 | 0.82944722 | 0.45 | 0.173 | 1.77E-75 | 3 | S1pr1 |
| 7.38E-61 | 0.80972585 | 0.326 | 0.113 | 1.36E-56 | 3 | Itga4 |
| 9.20E-60 | 0.78305902 | 0.582 | 0.313 | 1.69E-55 | 3 | Klf2 |
| 4.57E-59 | 0.66551196 | 0.294 | 0.096 | 8.41E-55 | 3 | Klf3 |
| 1.03E-30 | 0.62683901 | 0.383 | 0.213 | 1.90E-26 | 3 | Gm2682 |
| 6.38E-25 | 0.4953907 | 0.333 | 0.187 | 1.17E-20 | 3 | Slamf6 |
| 7.85E-23 | 0.51809033 | 0.653 | 0.573 | 1.44E-18 | 3 | Itgb1 |
| 1.82E-20 | 0.49016402 | 0.487 | 0.357 | 3.34E-16 | 3 | Gramd3 |
| 9.28E-18 | 0.57539867 | 0.306 | 0.191 | 1.71E-13 | 3 | Socs3 |
| 2.57E-15 | 0.69064361 | 0.271 | 0.17 | 4.73E-11 | 3 | Ramp3 |
| 8.00E-15 | 0.52208767 | 0.54 | 0.453 | 1.47E-10 | 3 | Zfp36 |
| 9.05E-09 | 0.95717442 | 0.373 | 0.299 | 0.00016643 | 3 | Ccl5 |
| 1.12E-74 | 0.93381503 | 0.717 | 0.323 | 2.07E-70 | 4 | Ifitm1 |
| 1.86E-68 | 0.76793303 | 0.828 | 0.452 | 3.42E-64 | 4 | Ifitm2 |
| 1.51E-64 | 0.97632924 | 0.451 | 0.159 | 2.78E-60 | 4 | Cd7 |
| 2.43E-59 | 0.7319595 | 0.976 | 0.872 | 4.47E-55 | 4 | Crip1 |
| 1.85E-57 | 0.53111512 | 0.987 | 0.895 | 3.41E-53 | 4 | S100a10 |
| 5.53E-52 | 0.52931111 | 0.944 | 0.774 | 1.02E-47 | 4 | Hcst |
| 1.10E-50 | 0.55735335 | 0.964 | 0.825 | 2.03E-46 | 4 | S100a6 |
| 4.39E-50 | 0.57485971 | 0.914 | 0.604 | 8.08E-46 | 4 | Cxcr6 |
| 1.64E-47 | 0.63651636 | 0.79 | 0.565 | 3.02E-43 | 4 | Itgb1 |
| 3.92E-45 | 0.78120399 | 0.68 | 0.423 | 7.20E-41 | 4 | Lgals3 |
| 6.53E-39 | 0.6506108 | 0.794 | 0.604 | 1.20E-34 | 4 | Btg2 |
| 9.87E-39 | 0.54160822 | 0.738 | 0.526 | 1.82E-34 | 4 | Zyx |
| 6.44E-26 | 0.53277663 | 0.365 | 0.177 | 1.18E-21 | 4 | Klrk1 |
| 4.27E-21 | 0.58620487 | 0.657 | 0.518 | 7.86E-17 | 4 | Ier2 |
| 1.49E-09 | 0.74384661 | 0.427 | 0.328 | 2.74E-05 | 4 | Fos |
| 0 | 1.8753127 | 0.85 | 0.076 | 0 | 5 | Stmn1 |
| 0 | 1.69803314 | 0.695 | 0.073 | 0 | 5 | Mki67 |
| 0 | 1.6156637 | 0.71 | 0.04 | 0 | 5 | Pclaf |
| 0 | 1.42764559 | 0.621 | 0.02 | 0 | 5 | Birc5 |
| 0 | 1.14503683 | 0.478 | 0.027 | 0 | 5 | Rrm2 |
| 0 | 1.07096042 | 0.489 | 0.013 | 0 | 5 | Ccnb2 |
| 1.06E-262 | 1.33823914 | 0.545 | 0.05 | 1.96E-258 | 5 | Ube2c |
| 5.20E-202 | 1.4032612 | 0.623 | 0.1 | 9.56E-198 | 5 | Top2a |
| 1.44E-169 | 1.60853664 | 0.982 | 0.657 | 2.65E-165 | 5 | Hmgb2 |
| 2.62E-165 | 1.192881 | 0.88 | 0.3 | 4.82E-161 | 5 | Hmgn2 |
| 2.93E-145 | 1.25165275 | 0.774 | 0.24 | 5.38E-141 | 5 | Tuba1b |
| 1.80E-138 | 1.15140838 | 0.977 | 0.702 | 3.32E-134 | 5 | H2afz |
| 5.52E-129 | 1.09328005 | 0.667 | 0.175 | 1.02E-124 | 5 | Cenpa |
| 1.79E-95 | 1.12691931 | 0.412 | 0.085 | 3.29E-91 | 5 | Hist1h2ae |
| 1.15E-93 | 1.28350118 | 0.893 | 0.544 | 2.12E-89 | 5 | Tubb5 |

| | | | | | | |
|---|---|---|---|---|---|---|
| 4.57E-148 | 2.0160125 | 0.92 | 0.367 | 8.42E-144 | 6 | Ikzf2 |
| 4.43E-141 | 1.78967043 | 0.811 | 0.245 | 8.14E-137 | 6 | Arl5a |
| 1.66E-109 | 1.71782553 | 0.873 | 0.39 | 3.05E-105 | 6 | Il2ra |
| 6.23E-109 | 1.53371572 | 0.938 | 0.674 | 1.15E-104 | 6 | Dock2 |
| 1.90E-103 | 1.6824923 | 0.716 | 0.254 | 3.50E-99 | 6 | Lrba |
| 3.24E-102 | 1.51315618 | 0.396 | 0.061 | 5.95E-98 | 6 | Itgb8 |
| 8.32E-100 | 1.52844725 | 0.876 | 0.561 | 1.53E-95 | 6 | Wnk1 |
| 1.79E-90 | 1.50631759 | 0.498 | 0.116 | 3.30E-86 | 6 | Neb |
| 4.52E-85 | 1.60805781 | 0.698 | 0.281 | 8.32E-81 | 6 | Itgav |
| 2.19E-80 | 1.49807374 | 0.807 | 0.467 | 4.03E-76 | 6 | Rabgap1l |
| 2.13E-70 | 1.40516686 | 0.789 | 0.51 | 3.92E-66 | 6 | Prkch |
| 6.88E-68 | 1.56506407 | 0.509 | 0.157 | 1.27E-63 | 6 | Alcam |
| 4.73E-64 | 1.44932062 | 0.665 | 0.332 | 8.70E-60 | 6 | Picalm |
| 2.29E-57 | 1.88671189 | 0.335 | 0.073 | 4.22E-53 | 6 | Frmd5 |
| 1.44E-45 | 1.43525883 | 0.753 | 0.573 | 2.65E-41 | 6 | Ctla4 |
| 6.40E-133 | 2.97789676 | 0.419 | 0.04 | 1.18E-128 | 7 | Ccl1 |
| 4.49E-107 | 3.20085859 | 0.3 | 0.024 | 8.27E-103 | 7 | Il2 |
| 2.45E-85 | 2.34541794 | 0.519 | 0.102 | 4.50E-81 | 7 | Csf2 |
| 1.59E-60 | 0.87078491 | 0.452 | 0.097 | 2.92E-56 | 7 | Mir155hg |
| 2.52E-60 | 1.02900908 | 0.557 | 0.146 | 4.64E-56 | 7 | Nr4a3 |
| 5.01E-59 | 0.93041253 | 0.39 | 0.076 | 9.22E-55 | 7 | Myc |
| 8.79E-59 | 1.26408884 | 0.671 | 0.231 | 1.62E-54 | 7 | Nfkbid |
| 6.64E-57 | 3.0568082 | 0.614 | 0.219 | 1.22E-52 | 7 | Ifng |
| 1.08E-52 | 0.95868742 | 0.867 | 0.411 | 1.98E-48 | 7 | Gadd45b |
| 2.43E-44 | 1.04825041 | 0.59 | 0.211 | 4.47E-40 | 7 | Tnfsf11 |
| 2.09E-38 | 1.15361179 | 0.719 | 0.352 | 3.85E-34 | 7 | Nr4a1 |
| 3.12E-32 | 1.40422485 | 0.471 | 0.174 | 5.73E-28 | 7 | Tnf |
| 3.76E-31 | 0.87205856 | 0.486 | 0.18 | 6.92E-27 | 7 | Irf8 |
| 6.49E-24 | 0.99436094 | 0.667 | 0.393 | 1.19E-19 | 7 | Cd40lg |
| 1.43E-05 | 1.79053916 | 0.286 | 0.184 | 0.26292439 | 7 | Ccl4 |
| 7.40E-143 | 0.69877251 | 0.374 | 0.027 | 1.36E-138 | 8 | Bace2 |
| 2.54E-83 | 1.09693654 | 0.7 | 0.173 | 4.67E-79 | 8 | Tnfsf8 |
| 2.43E-82 | 0.98221929 | 0.709 | 0.178 | 4.47E-78 | 8 | Tcf7 |
| 1.09E-77 | 0.99156289 | 0.7 | 0.187 | 2.00E-73 | 8 | Slamf6 |
| 3.69E-66 | 0.75780459 | 0.389 | 0.065 | 6.79E-62 | 8 | Gpm6b |
| 5.11E-61 | 0.7859786 | 0.404 | 0.074 | 9.40E-57 | 8 | Aff3 |
| 2.62E-59 | 0.66725718 | 0.448 | 0.093 | 4.82E-55 | 8 | Nsg2 |
| 4.82E-56 | 0.71179456 | 0.463 | 0.103 | 8.86E-52 | 8 | Tbc1d4 |
| 4.42E-47 | 0.65851449 | 0.512 | 0.142 | 8.13E-43 | 8 | Asap1 |
| 2.60E-46 | 0.88715838 | 0.355 | 0.073 | 4.79E-42 | 8 | Ccr7 |
| 5.44E-35 | 0.74218717 | 0.734 | 0.363 | 1.00E-30 | 8 | Sh2d1a |
| 2.00E-33 | 0.60508358 | 0.35 | 0.092 | 3.68E-29 | 8 | Tg |
| 1.96E-29 | 0.68131669 | 0.384 | 0.119 | 3.60E-25 | 8 | Nmrk1 |
| 5.70E-28 | 0.78191675 | 0.369 | 0.118 | 1.05E-23 | 8 | Marcksl1 |
| 8.72E-20 | 0.69572124 | 0.404 | 0.165 | 1.60E-15 | 8 | Il1r2 |
| 6.87E-81 | 1.59661423 | 0.849 | 0.313 | 1.26E-76 | 9 | Hivep2 |

| | | | | | | |
|---|---|---|---|---|---|---|
| 8.13E-79 | 1.83609347 | 0.861 | 0.378 | 1.50E-74 | 9 | Themis |
| 1.47E-74 | 1.61615348 | 0.886 | 0.379 | 2.71E-70 | 9 | Atxn1 |
| 1.61E-73 | 1.58553197 | 0.958 | 0.679 | 2.96E-69 | 9 | Dock2 |
| 3.21E-70 | 1.55119921 | 0.892 | 0.467 | 5.90E-66 | 9 | Elmo1 |
| 7.01E-66 | 1.68797629 | 0.771 | 0.314 | 1.29E-61 | 9 | Arap2 |
| 2.95E-64 | 1.5574518 | 0.699 | 0.223 | 5.43E-60 | 9 | Maml2 |
| 3.58E-64 | 1.60217235 | 0.904 | 0.517 | 6.59E-60 | 9 | Arhgap15 |
| 4.63E-63 | 1.93062353 | 0.681 | 0.216 | 8.52E-59 | 9 | St6galnac3 |
| 2.77E-62 | 1.70797002 | 0.892 | 0.543 | 5.10E-58 | 9 | Inpp4b |
| 4.04E-61 | 1.55249709 | 0.663 | 0.207 | 7.44E-57 | 9 | Arl15 |
| 8.58E-56 | 1.5662296 | 0.819 | 0.44 | 1.58E-51 | 9 | Stat4 |
| 8.13E-54 | 1.60495507 | 0.831 | 0.476 | 1.49E-49 | 9 | Rora |
| 5.87E-51 | 1.68749363 | 0.813 | 0.448 | 1.08E-46 | 9 | Dennd4a |
| 5.18E-49 | 1.56479992 | 0.675 | 0.283 | 9.53E-45 | 9 | Itpr1 |
| 1.69E-110 | 1.8232829 | 0.914 | 0.241 | 3.11E-106 | 10 | Ifit1 |
| 1.07E-84 | 1.44917424 | 0.753 | 0.195 | 1.97E-80 | 10 | Usp18 |
| 6.02E-83 | 1.27935562 | 0.611 | 0.115 | 1.11E-78 | 10 | Rsad2 |
| 7.49E-82 | 1.29240424 | 0.704 | 0.161 | 1.38E-77 | 10 | Iigp1 |
| 2.70E-80 | 1.74580922 | 0.988 | 0.539 | 4.97E-76 | 10 | Isg15 |
| 1.02E-75 | 1.42200414 | 0.957 | 0.491 | 1.88E-71 | 10 | Isg20 |
| 9.35E-70 | 1.4963974 | 0.759 | 0.217 | 1.72E-65 | 10 | Ifit3 |
| 2.99E-66 | 1.38725349 | 0.858 | 0.376 | 5.51E-62 | 10 | Phf11b |
| 8.15E-66 | 1.56617474 | 0.963 | 0.522 | 1.50E-61 | 10 | Gzmb |
| 1.57E-60 | 1.17936829 | 0.543 | 0.124 | 2.89E-56 | 10 | Ccrl2 |
| 5.25E-56 | 1.21113698 | 0.951 | 0.625 | 9.67E-52 | 10 | Samhd1 |
| 1.30E-53 | 1.07654721 | 0.963 | 0.648 | 2.39E-49 | 10 | Bst2 |
| 2.46E-51 | 1.38713975 | 0.438 | 0.09 | 4.53E-47 | 10 | Cxcl10 |
| 1.46E-48 | 1.1824392 | 0.84 | 0.418 | 2.69E-44 | 10 | Irf7 |
| 6.53E-43 | 1.10947208 | 0.407 | 0.092 | 1.20E-38 | 10 | Mx1 |
| 1.43E-301 | 1.40457584 | 0.543 | 0.011 | 2.62E-297 | 11 | Klra7 |
| 6.66E-188 | 0.66373137 | 0.309 | 0.005 | 1.23E-183 | 11 | Klra6 |
| 3.40E-134 | 1.49656462 | 0.66 | 0.052 | 6.26E-130 | 11 | Fcer1g |
| 1.84E-102 | 1.21710381 | 0.511 | 0.039 | 3.39E-98 | 11 | Xcl1 |
| 1.55E-93 | 0.73607019 | 0.436 | 0.031 | 2.85E-89 | 11 | Cd160 |
| 1.59E-56 | 1.60630374 | 0.947 | 0.298 | 2.93E-52 | 11 | Ccl5 |
| 9.05E-56 | 1.29510625 | 0.766 | 0.172 | 1.66E-51 | 11 | Cd7 |
| 1.84E-48 | 1.24550256 | 0.894 | 0.271 | 3.38E-44 | 11 | Ly6c2 |
| 2.18E-37 | 0.82637323 | 0.511 | 0.103 | 4.01E-33 | 11 | Klhdc2 |
| 5.41E-24 | 0.63509512 | 0.574 | 0.169 | 9.96E-20 | 11 | Sidt1 |
| 7.74E-24 | 0.7381619 | 0.745 | 0.293 | 1.42E-19 | 11 | Bcl2 |
| 2.80E-22 | 0.80579728 | 0.66 | 0.227 | 5.15E-18 | 11 | Gm2682 |
| 5.24E-16 | 0.74202554 | 0.404 | 0.125 | 9.65E-12 | 11 | Bach2 |
| 1.30E-14 | 0.62743383 | 0.415 | 0.141 | 2.39E-10 | 11 | Pim2 |
| 4.78E-13 | 0.74958779 | 0.5 | 0.211 | 8.79E-09 | 11 | Gas5 |
| 4.46E-241 | 2.05157356 | 0.588 | 0.017 | 8.21E-237 | 12 | Igfbp4 |
| 8.73E-111 | 1.68604622 | 0.75 | 0.074 | 1.61E-106 | 12 | Ccr7 |

| | | | | | | |
|---|---|---|---|---|---|---|
| 2.76E-106 | 1.08923182 | 0.325 | 0.012 | 5.09E-102 | 12 | Dapl1 |
| 1.71E-63 | 1.19475129 | 0.488 | 0.053 | 3.15E-59 | 12 | Rflnb |
| 2.52E-47 | 1.0541604 | 0.475 | 0.069 | 4.64E-43 | 12 | Actn1 |
| 6.70E-47 | 1.37667064 | 0.762 | 0.188 | 1.23E-42 | 12 | Tcf7 |
| 6.91E-46 | 1.67798887 | 0.925 | 0.337 | 1.27E-41 | 12 | Klf2 |
| 1.29E-44 | 0.95250074 | 1 | 0.978 | 2.37E-40 | 12 | Rplp1 |
| 6.60E-42 | 1.00839185 | 1 | 0.938 | 1.21E-37 | 12 | Rps29 |
| 1.84E-37 | 0.95551586 | 1 | 0.901 | 3.39E-33 | 12 | Rpl12 |
| 5.09E-37 | 1.02864961 | 0.988 | 0.851 | 9.36E-33 | 12 | Rpl36a |
| 2.05E-31 | 1.24400915 | 0.713 | 0.245 | 3.78E-27 | 12 | Lef1 |
| 3.07E-29 | 1.03315228 | 0.675 | 0.2 | 5.64E-25 | 12 | S1pr1 |
| 1.15E-26 | 1.3896146 | 0.7 | 0.262 | 2.12E-22 | 12 | Jun |
| 1.08E-15 | 0.95264996 | 0.5 | 0.193 | 1.99E-11 | 12 | Selenop |

**Supplementary Table 2.** Genes differentially expressed between tumor-infiltrating TREG from untreated and aCTLA-4 treated mice.

| Gene Symbol | p_val | avg_logFC | pct.1 | pct.2 | p_val_adj |
|---|---|---|---|---|---|
| Lag3 | 1.34E-63 | -1.781141 | 0.079 | 0.578 | 2.46E-59 |
| Ccl4 | 2.05E-11 | -1.343953 | 0.182 | 0.315 | 3.77E-07 |
| Tigit | 2.58E-55 | -1.110134 | 0.52 | 0.826 | 4.75E-51 |
| Il10 | 4.60E-28 | -1.077316 | 0.03 | 0.302 | 8.46E-24 |
| Cst7 | 1.10E-37 | -1.034112 | 0.222 | 0.57 | 2.03E-33 |
| Gzmb | 3.18E-64 | -1.03341 | 0.55 | 0.931 | 5.85E-60 |
| Ccr8 | 6.40E-26 | -1.00271 | 0.182 | 0.47 | 1.18E-21 |
| Sv2c | 4.29E-34 | -0.989161 | 0.076 | 0.418 | 7.89E-30 |
| Il2ra | 1.60E-44 | -0.965594 | 0.593 | 0.861 | 2.95E-40 |
| Frmd5 | 7.16E-18 | -0.94762 | 0.079 | 0.315 | 1.32E-13 |
| Arl5a | 1.03E-29 | -0.860966 | 0.368 | 0.674 | 1.89E-25 |
| Entpd1 | 3.75E-27 | -0.85871 | 0.219 | 0.532 | 6.90E-23 |
| Itgb8 | 2.01E-22 | -0.848595 | 0.046 | 0.294 | 3.70E-18 |
| Klrg1 | 2.06E-24 | -0.847012 | 0.252 | 0.567 | 3.78E-20 |
| Nkg7 | 2.78E-14 | -0.831606 | 0.411 | 0.606 | 5.12E-10 |
| Tmem163 | 1.83E-17 | -0.826842 | 0.036 | 0.218 | 3.37E-13 |
| Litaf | 1.12E-24 | -0.825115 | 0.202 | 0.48 | 2.05E-20 |
| Sdf4 | 5.97E-46 | -0.817448 | 0.838 | 0.928 | 1.10E-41 |
| Anxa2 | 2.36E-29 | -0.802417 | 0.454 | 0.746 | 4.35E-25 |
| Ctsz | 2.87E-23 | -0.77451 | 0.285 | 0.539 | 5.27E-19 |
| Maf | 5.07E-30 | -0.756756 | 0.563 | 0.769 | 9.32E-26 |
| Ap3b1 | 4.66E-15 | -0.746702 | 0.321 | 0.507 | 8.57E-11 |
| Isg20 | 1.30E-21 | -0.738644 | 0.454 | 0.706 | 2.39E-17 |
| Tnfrsf4 | 4.05E-42 | -0.722409 | 0.891 | 0.965 | 7.44E-38 |
| Zfp36l1 | 5.07E-26 | -0.709205 | 0.616 | 0.807 | 9.32E-22 |
| Coro2a | 1.76E-19 | -0.704718 | 0.182 | 0.439 | 3.24E-15 |
| Icos | 7.61E-35 | -0.699385 | 0.907 | 0.951 | 1.40E-30 |
| Itgav | 6.79E-18 | -0.692345 | 0.424 | 0.642 | 1.25E-13 |
| Asb2 | 6.87E-19 | -0.683548 | 0.076 | 0.302 | 1.26E-14 |
| Mdfic | 5.94E-18 | -0.679368 | 0.265 | 0.494 | 1.09E-13 |
| Ccr5 | 1.15E-16 | -0.674132 | 0.255 | 0.507 | 2.12E-12 |
| Pglyrp1 | 1.41E-22 | -0.662922 | 0.662 | 0.756 | 2.60E-18 |
| Gimap7 | 6.03E-28 | -0.661845 | 0.636 | 0.78 | 1.11E-23 |
| Pcyt1a | 2.75E-19 | -0.657915 | 0.053 | 0.28 | 5.06E-15 |
| Alcam | 1.17E-11 | -0.657402 | 0.159 | 0.346 | 2.14E-07 |
| Hip1 | 1.30E-13 | -0.642976 | 0.142 | 0.343 | 2.40E-09 |
| Glrx | 1.25E-16 | -0.641858 | 0.652 | 0.76 | 2.31E-12 |
| Nckap5 | 1.16E-13 | -0.636464 | 0.017 | 0.161 | 2.13E-09 |
| Ddit4 | 1.94E-16 | -0.633515 | 0.447 | 0.659 | 3.57E-12 |
| Max | 7.37E-14 | -0.622992 | 0.281 | 0.472 | 1.36E-09 |
| Havcr2 | 5.38E-16 | -0.615519 | 0.096 | 0.31 | 9.89E-12 |
| Snx9 | 9.85E-16 | -0.613419 | 0.209 | 0.451 | 1.81E-11 |

| | | | | | |
|---|---|---|---|---|---|
| Ctsb | 1.77E-21 | -0.603936 | 0.54 | 0.688 | 3.26E-17 |
| Ccl5 | 1.39E-08 | -0.603658 | 0.275 | 0.461 | 0.000256 |
| Ptms | 4.16E-16 | -0.597965 | 0.275 | 0.49 | 7.66E-12 |
| Mt1 | 6.64E-07 | -0.595633 | 0.103 | 0.23 | 0.012221 |
| Rbpj | 6.32E-11 | -0.585757 | 0.139 | 0.273 | 1.16E-06 |
| Neb | 5.25E-11 | -0.585694 | 0.205 | 0.394 | 9.66E-07 |
| Cep112 | 3.22E-15 | -0.581875 | 0.013 | 0.166 | 5.92E-11 |
| Ehd1 | 2.06E-16 | -0.580051 | 0.209 | 0.421 | 3.80E-12 |
| Socs2 | 5.85E-14 | -0.578365 | 0.228 | 0.438 | 1.08E-09 |
| Runx2 | 5.90E-11 | -0.574216 | 0.281 | 0.467 | 1.09E-06 |
| Sdcbp2 | 1.20E-14 | -0.573565 | 0.281 | 0.478 | 2.21E-10 |
| Nrip1 | 9.90E-11 | -0.571661 | 0.225 | 0.403 | 1.82E-06 |
| Itih5 | 9.68E-18 | -0.571067 | 0.02 | 0.189 | 1.78E-13 |
| Rabgap1l | 3.02E-12 | -0.570795 | 0.48 | 0.674 | 5.56E-08 |
| Casp3 | 2.08E-15 | -0.567163 | 0.225 | 0.428 | 3.83E-11 |
| Tnfrsf1b | 6.23E-19 | -0.557001 | 0.56 | 0.724 | 1.15E-14 |
| Lrba | 1.44E-07 | -0.55345 | 0.331 | 0.456 | 0.002647 |
| Sytl3 | 1.49E-14 | -0.548126 | 0.129 | 0.319 | 2.75E-10 |
| Osbpl3 | 1.11E-11 | -0.541971 | 0.189 | 0.361 | 2.05E-07 |
| Tnfrsf9 | 5.46E-29 | -0.539707 | 0.762 | 0.928 | 1.00E-24 |
| Nfil3 | 5.95E-13 | -0.53792 | 0.318 | 0.488 | 1.10E-08 |
| Odc1 | 2.68E-13 | -0.53416 | 0.672 | 0.777 | 4.93E-09 |
| Cmtm7 | 8.42E-21 | -0.525313 | 0.576 | 0.667 | 1.55E-16 |
| N4bp1 | 1.18E-09 | -0.520997 | 0.209 | 0.373 | 2.17E-05 |
| Cdk6 | 1.27E-11 | -0.518769 | 0.228 | 0.426 | 2.34E-07 |
| Irf7 | 5.52E-09 | -0.51207 | 0.454 | 0.517 | 0.000102 |
| Oxsr1 | 1.08E-09 | -0.510499 | 0.116 | 0.242 | 2.00E-05 |
| Wnk1 | 4.86E-16 | -0.507697 | 0.685 | 0.826 | 8.95E-12 |
| M6pr | 1.33E-14 | -0.505225 | 0.497 | 0.626 | 2.45E-10 |
| Bst2 | 5.88E-14 | -0.503026 | 0.675 | 0.739 | 1.08E-09 |
| Nedd9 | 1.11E-11 | -0.499437 | 0.318 | 0.49 | 2.03E-07 |
| H2-T22 | 3.38E-20 | -0.494535 | 0.788 | 0.74 | 6.21E-16 |
| Mmd | 6.97E-12 | -0.49353 | 0.096 | 0.276 | 1.28E-07 |
| Chchd10 | 2.09E-09 | -0.492305 | 0.149 | 0.294 | 3.84E-05 |
| Pik3ap1 | 1.76E-14 | -0.491983 | 0.036 | 0.195 | 3.23E-10 |
| Cmss1 | 1.09E-06 | -0.484141 | 0.377 | 0.536 | 0.019991 |
| Tax1bp1 | 6.68E-13 | -0.479342 | 0.586 | 0.71 | 1.23E-08 |
| Rilpl2 | 2.77E-09 | -0.478545 | 0.394 | 0.533 | 5.09E-05 |
| Fgl2 | 3.65E-11 | -0.477762 | 0.212 | 0.424 | 6.71E-07 |
| Gapdh | 1.45E-13 | -0.471599 | 0.725 | 0.787 | 2.67E-09 |
| Sat1 | 3.48E-12 | -0.470423 | 0.51 | 0.603 | 6.41E-08 |
| Eea1 | 7.65E-08 | -0.470298 | 0.275 | 0.422 | 0.001407 |
| Lamc1 | 3.70E-07 | -0.469507 | 0.272 | 0.421 | 0.006817 |
| Rab11a | 5.63E-15 | -0.468437 | 0.503 | 0.584 | 1.04E-10 |
| Aopep | 2.86E-07 | -0.466411 | 0.199 | 0.309 | 0.005257 |
| Smyd3 | 1.89E-06 | -0.466028 | 0.119 | 0.205 | 0.03473 |

| | | | | | |
|---|---|---|---|---|---|
| Grn | 7.23E-11 | -0.464328 | 0.136 | 0.304 | 1.33E-06 |
| Gm19585 | 5.27E-11 | -0.459503 | 0.212 | 0.36 | 9.71E-07 |
| Ormdl3 | 4.99E-09 | -0.455387 | 0.172 | 0.316 | 9.18E-05 |
| Vgll4 | 8.44E-10 | -0.453923 | 0.358 | 0.484 | 1.55E-05 |
| Ikzf3 | 2.86E-08 | -0.452697 | 0.421 | 0.564 | 0.000526 |
| Prkch | 1.42E-13 | -0.452385 | 0.616 | 0.754 | 2.62E-09 |
| Vmp1 | 6.94E-12 | -0.446953 | 0.424 | 0.57 | 1.28E-07 |
| 1700017B05Rik | 2.49E-09 | -0.443756 | 0.156 | 0.288 | 4.58E-05 |
| Irak2 | 1.31E-07 | -0.443533 | 0.219 | 0.329 | 0.002419 |
| Cd81 | 5.45E-08 | -0.440578 | 0.156 | 0.279 | 0.001004 |
| Phlda1 | 2.50E-07 | -0.439416 | 0.209 | 0.313 | 0.004597 |
| Ly6a | 6.37E-23 | -0.435531 | 0.94 | 0.984 | 1.17E-18 |
| Ciapin1 | 2.66E-10 | -0.434176 | 0.202 | 0.356 | 4.90E-06 |
| Mrps6 | 3.34E-10 | -0.433944 | 0.086 | 0.233 | 6.14E-06 |
| Ndfip1 | 2.06E-19 | -0.432646 | 0.848 | 0.901 | 3.80E-15 |
| Pkm | 6.46E-17 | -0.43226 | 0.851 | 0.868 | 1.19E-12 |
| Ube2l6 | 5.49E-07 | -0.431265 | 0.192 | 0.31 | 0.010106 |
| Supt3 | 1.09E-09 | -0.430032 | 0.06 | 0.203 | 2.01E-05 |
| Izumo1r | 3.68E-08 | -0.427838 | 0.513 | 0.582 | 0.000676 |
| Cysltr2 | 8.92E-07 | -0.426208 | 0.066 | 0.173 | 0.016409 |
| Lrrc32 | 1.01E-06 | -0.425565 | 0.228 | 0.375 | 0.018574 |
| Plp2 | 3.04E-11 | -0.422962 | 0.411 | 0.53 | 5.60E-07 |
| Rgs16 | 2.69E-09 | -0.422155 | 0.063 | 0.204 | 4.95E-05 |
| Tnfrsf13b | 1.19E-08 | -0.418689 | 0.172 | 0.286 | 0.00022 |
| Bnip3 | 1.63E-06 | -0.418173 | 0.119 | 0.249 | 0.029907 |
| Zap70 | 2.04E-11 | -0.414757 | 0.464 | 0.552 | 3.76E-07 |
| Tbx21 | 5.17E-10 | -0.414625 | 0.136 | 0.239 | 9.51E-06 |
| Bmyc | 1.06E-06 | -0.411308 | 0.113 | 0.215 | 0.01953 |
| Spp1 | 9.07E-08 | -0.410598 | 0.017 | 0.102 | 0.001669 |
| Cd28 | 1.55E-14 | -0.410219 | 0.649 | 0.671 | 2.85E-10 |
| Bcl2l1 | 9.70E-07 | -0.409082 | 0.272 | 0.372 | 0.01785 |
| Cd2 | 9.15E-23 | -0.40901 | 0.94 | 0.961 | 1.68E-18 |
| Dusp16 | 4.48E-08 | -0.40835 | 0.03 | 0.133 | 0.000825 |
| Il7r | 6.47E-07 | -0.406786 | 0.444 | 0.582 | 0.011913 |
| Skil | 1.17E-07 | -0.405514 | 0.169 | 0.305 | 0.00216 |
| Tnfrsf18 | 2.33E-20 | -0.403147 | 0.93 | 0.961 | 4.29E-16 |
| Twsg1 | 6.45E-08 | -0.402352 | 0.083 | 0.207 | 0.001187 |
| Dnajc1 | 2.87E-08 | -0.397775 | 0.417 | 0.542 | 0.000528 |
| Srgap3 | 9.07E-10 | -0.397597 | 0.01 | 0.103 | 1.67E-05 |
| Dctpp1 | 1.69E-09 | -0.39756 | 0.139 | 0.289 | 3.11E-05 |
| Plek | 1.81E-08 | -0.396825 | 0.036 | 0.148 | 0.000332 |
| Mier1 | 2.43E-07 | -0.396601 | 0.457 | 0.583 | 0.004477 |
| Leprotl1 | 9.59E-15 | -0.395164 | 0.755 | 0.804 | 1.77E-10 |
| Fam110a | 1.95E-07 | -0.394748 | 0.202 | 0.329 | 0.003592 |
| Rhof | 1.04E-07 | -0.393472 | 0.371 | 0.472 | 0.001918 |

| | | | | | |
|---|---|---|---|---|---|
| Timp2 | 2.08E-06 | -0.391467 | 0.182 | 0.293 | 0.038258 |
| Glcci1 | 2.47E-06 | -0.390506 | 0.136 | 0.205 | 0.045401 |
| Ptpn22 | 1.48E-06 | -0.389517 | 0.397 | 0.51 | 0.027163 |
| Hdac7 | 2.41E-07 | -0.388669 | 0.219 | 0.338 | 0.004434 |
| Fem1c | 6.37E-07 | -0.386132 | 0.149 | 0.264 | 0.011719 |
| Lfng | 1.07E-06 | -0.38438 | 0.311 | 0.441 | 0.019745 |
| Rad51b | 6.13E-10 | -0.383066 | 0.013 | 0.118 | 1.13E-05 |
| Npc2 | 2.03E-16 | -0.382486 | 0.844 | 0.855 | 3.74E-12 |
| Lgals1 | 4.40E-15 | -0.381377 | 0.93 | 0.961 | 8.10E-11 |
| Wls | 1.72E-06 | -0.380866 | 0.192 | 0.325 | 0.031734 |
| Ergic1 | 1.29E-07 | -0.374283 | 0.159 | 0.291 | 0.002376 |
| Ltb4r1 | 2.16E-07 | -0.373691 | 0.089 | 0.22 | 0.003981 |
| Got1 | 2.03E-08 | -0.373006 | 0.368 | 0.473 | 0.000373 |
| Ebi3 | 6.79E-07 | -0.371718 | 0.169 | 0.291 | 0.012488 |
| Fbxw11 | 3.10E-07 | -0.370512 | 0.179 | 0.297 | 0.00571 |
| Angptl2 | 2.78E-12 | -0.369813 | 0.003 | 0.112 | 5.11E-08 |
| Gimap5 | 8.84E-08 | -0.368352 | 0.517 | 0.6 | 0.001626 |
| Capg | 1.35E-15 | -0.367058 | 0.778 | 0.839 | 2.48E-11 |
| Psd4 | 1.67E-07 | -0.360409 | 0.248 | 0.371 | 0.003079 |
| Cd101 | 5.10E-10 | -0.359459 | 0.036 | 0.164 | 9.39E-06 |
| Cd80 | 1.61E-07 | -0.35677 | 0.07 | 0.193 | 0.002964 |
| Rapsn | 5.34E-08 | -0.355603 | 0.076 | 0.194 | 0.000983 |
| Dusp5 | 1.15E-08 | -0.354264 | 0.606 | 0.696 | 0.000211 |
| Sema4d | 1.07E-08 | -0.351674 | 0.52 | 0.612 | 0.000196 |
| Sae1 | 1.92E-07 | -0.350875 | 0.245 | 0.346 | 0.003531 |
| Trim30a | 1.04E-06 | -0.350096 | 0.43 | 0.525 | 0.019171 |
| Gimap3 | 3.08E-13 | -0.348844 | 0.828 | 0.851 | 5.66E-09 |
| Myo10 | 6.71E-09 | -0.345348 | 0.01 | 0.102 | 0.000123 |
| Cd27 | 2.55E-09 | -0.345006 | 0.56 | 0.661 | 4.70E-05 |
| Vsir | 9.98E-08 | -0.34409 | 0.454 | 0.511 | 0.001836 |
| Gm17767 | 1.18E-09 | -0.342513 | 0.046 | 0.183 | 2.17E-05 |
| AW112010 | 1.12E-10 | -0.341223 | 0.944 | 0.957 | 2.06E-06 |
| Il2rb | 6.10E-16 | -0.339227 | 0.828 | 0.835 | 1.12E-11 |
| Prf1 | 4.60E-07 | -0.334967 | 0.023 | 0.113 | 0.008469 |
| Srp14 | 1.05E-12 | -0.333963 | 0.583 | 0.624 | 1.93E-08 |
| Leprot | 1.82E-07 | -0.33037 | 0.166 | 0.272 | 0.003351 |
| Cst3 | 3.18E-07 | -0.326512 | 0.417 | 0.509 | 0.00585 |
| Snx2 | 1.96E-08 | -0.32571 | 0.487 | 0.545 | 0.00036 |
| Sep-07 | 4.74E-08 | -0.324836 | 0.573 | 0.663 | 0.000872 |
| Grina | 2.43E-06 | -0.323686 | 0.328 | 0.415 | 0.044738 |
| Jdp2 | 1.13E-06 | -0.322229 | 0.023 | 0.106 | 0.020779 |
| Pvrig | 2.40E-09 | -0.320447 | 0.043 | 0.173 | 4.41E-05 |
| Ikzf2 | 9.07E-08 | -0.318634 | 0.887 | 0.911 | 0.00167 |
| Akap13 | 3.29E-08 | -0.317724 | 0.719 | 0.748 | 0.000605 |
| Arf6 | 1.42E-09 | -0.316683 | 0.603 | 0.617 | 2.61E-05 |

| | | | | | |
|---|---|---|---|---|---|
| Pigx | 1.30E-06 | -0.312717 | 0.404 | 0.48 | 0.023908 |
| Ifitm3 | 3.67E-09 | -0.309732 | 0.301 | 0.504 | 6.75E-05 |
| Lrrfip1 | 1.59E-07 | -0.305903 | 0.55 | 0.6 | 0.002934 |
| H2-K1 | 1.97E-15 | -0.305549 | 0.997 | 1 | 3.62E-11 |
| Camk2n1 | 2.13E-06 | -0.304415 | 0.05 | 0.144 | 0.039276 |
| Eno1 | 7.36E-08 | -0.303535 | 0.579 | 0.585 | 0.001355 |
| Anxa11 | 9.51E-07 | -0.301956 | 0.407 | 0.49 | 0.017503 |
| Plin2 | 8.30E-07 | -0.301492 | 0.086 | 0.212 | 0.015273 |
| Tnfrsf8 | 2.06E-06 | -0.29872 | 0.043 | 0.141 | 0.037874 |
| Psme2 | 1.38E-12 | -0.298245 | 0.848 | 0.866 | 2.53E-08 |
| Lat2 | 2.27E-07 | -0.296141 | 0.046 | 0.156 | 0.004169 |
| Trac | 1.92E-08 | -0.293735 | 0.656 | 0.698 | 0.000354 |
| Cyth4 | 1.39E-07 | -0.289089 | 0.407 | 0.471 | 0.002565 |
| Crlf2 | 1.79E-07 | -0.288547 | 0.543 | 0.579 | 0.003285 |
| Gm2a | 1.57E-06 | -0.284554 | 0.467 | 0.494 | 0.028971 |
| Aldoa | 2.02E-08 | -0.282712 | 0.798 | 0.805 | 0.000372 |
| Cystm1 | 9.55E-08 | -0.280515 | 0.03 | 0.134 | 0.001757 |
| Fxyd5 | 1.95E-10 | -0.276664 | 0.95 | 0.964 | 3.59E-06 |
| Reep5 | 1.07E-08 | -0.276567 | 0.672 | 0.721 | 0.000198 |
| Igtp | 6.14E-09 | -0.276437 | 0.662 | 0.676 | 0.000113 |
| Rap1b | 1.93E-10 | -0.274212 | 0.848 | 0.851 | 3.54E-06 |
| Cflar | 7.10E-07 | -0.273362 | 0.414 | 0.446 | 0.013071 |
| Arl6ip1 | 3.80E-10 | -0.272086 | 0.861 | 0.868 | 6.99E-06 |
| Grcc10 | 2.73E-07 | -0.268861 | 0.586 | 0.593 | 0.005032 |
| Ctss | 3.89E-07 | -0.268828 | 0.563 | 0.585 | 0.007164 |
| H2-D1 | 2.44E-09 | -0.268215 | 0.993 | 0.996 | 4.50E-05 |
| Tes | 6.59E-07 | -0.265614 | 0.526 | 0.55 | 0.012126 |
| Ywhah | 2.04E-08 | -0.258248 | 0.573 | 0.596 | 0.000376 |
| H2-Q6 | 1.67E-07 | -0.255017 | 0.662 | 0.65 | 0.003064 |
| Sh2d2a | 4.91E-07 | -0.253641 | 0.53 | 0.558 | 0.009037 |
| H2-Q7 | 1.44E-09 | -0.248816 | 0.914 | 0.909 | 2.64E-05 |
| Ctla4 | 2.59E-08 | -0.248781 | 0.874 | 0.785 | 0.000477 |
| Sep-09 | 1.89E-09 | -0.248745 | 0.596 | 0.608 | 3.48E-05 |
| Cish | 1.34E-06 | -0.238674 | 0.57 | 0.595 | 0.024598 |
| Ctsd | 2.64E-06 | -0.236495 | 0.503 | 0.521 | 0.048659 |
| Cd3e | 1.01E-10 | -0.235665 | 1 | 1 | 1.87E-06 |
| Anxa6 | 3.30E-09 | -0.235532 | 0.679 | 0.687 | 6.07E-05 |
| Vim | 1.07E-06 | -0.232404 | 0.927 | 0.937 | 0.019734 |
| Pkp3 | 3.97E-07 | -0.231261 | 0.526 | 0.548 | 0.007296 |
| Bzw1 | 6.80E-08 | -0.227432 | 0.526 | 0.522 | 0.001252 |
| Sla | 7.39E-07 | -0.226944 | 0.682 | 0.677 | 0.013602 |
| Cd3d | 1.90E-08 | -0.226917 | 0.934 | 0.937 | 0.000349 |
| Tmbim6 | 5.62E-08 | -0.220637 | 0.861 | 0.864 | 0.001034 |
| Ypel3 | 4.28E-07 | -0.216565 | 0.639 | 0.622 | 0.007873 |
| Hmgb1 | 6.77E-09 | -0.214429 | 0.758 | 0.743 | 0.000125 |
| Foxp1 | 2.15E-06 | -0.211537 | 0.397 | 0.346 | 0.039502 |

| | | | | | |
|---|---|---|---|---|---|
| Capza2 | 5.20E-07 | -0.203521 | 0.616 | 0.611 | 0.00957 |
| H2-T23 | 2.37E-07 | -0.201219 | 0.864 | 0.842 | 0.004354 |
| Snx3 | 2.13E-07 | -0.201106 | 0.589 | 0.592 | 0.003921 |
| Ppp1r18 | 4.31E-10 | -0.197767 | 0.702 | 0.653 | 7.92E-06 |
| Tap1 | 1.65E-06 | -0.192224 | 0.722 | 0.693 | 0.030306 |
| Psmb9 | 3.08E-08 | -0.186928 | 0.758 | 0.723 | 0.000566 |
| Prkar1a | 1.82E-07 | -0.186377 | 0.745 | 0.739 | 0.003344 |
| Gimap1 | 1.00E-06 | -0.184165 | 0.659 | 0.64 | 0.01849 |
| H2-Q4 | 1.29E-06 | -0.181018 | 0.695 | 0.644 | 0.023786 |
| Slc9a3r1 | 4.42E-07 | -0.17531 | 0.589 | 0.572 | 0.008124 |
| Psma3 | 2.36E-07 | -0.170335 | 0.629 | 0.597 | 0.004342 |
| Malat1 | 2.05E-06 | -0.170302 | 0.997 | 0.997 | 0.037742 |
| Arf1 | 9.45E-08 | -0.168891 | 0.685 | 0.667 | 0.001738 |
| Capns1 | 2.09E-06 | -0.166667 | 0.503 | 0.484 | 0.038479 |
| Tpm3 | 8.57E-10 | -0.166412 | 0.831 | 0.778 | 1.58E-05 |
| Cd5 | 7.68E-08 | -0.150311 | 0.772 | 0.732 | 0.001412 |
| Selenow | 3.26E-09 | -0.148906 | 0.844 | 0.785 | 6.01E-05 |
| Itm2c | 1.21E-07 | -0.11731 | 0.666 | 0.581 | 0.002233 |
| Son | 2.04E-06 | 0.102127 | 0.785 | 0.633 | 0.037518 |
| Isca2 | 1.97E-06 | 0.11442 | 0.238 | 0.129 | 0.036167 |
| Mrpl51 | 8.18E-07 | 0.121144 | 0.298 | 0.174 | 0.015052 |
| Usp28 | 1.90E-07 | 0.129386 | 0.103 | 0.026 | 0.003487 |
| 4930523C07Rik | 5.38E-07 | 0.154379 | 0.454 | 0.299 | 0.009894 |
| Etfa | 5.92E-07 | 0.161147 | 0.325 | 0.184 | 0.010899 |
| Cops9 | 3.19E-07 | 0.163493 | 0.55 | 0.38 | 0.005878 |
| Arid4a | 1.79E-06 | 0.171499 | 0.447 | 0.293 | 0.032875 |
| Hspe1 | 8.87E-08 | 0.182738 | 0.543 | 0.361 | 0.001633 |
| Rpl9 | 1.42E-07 | 0.188703 | 0.977 | 0.937 | 0.00262 |
| Lpxn | 8.17E-08 | 0.188871 | 0.414 | 0.25 | 0.001504 |
| Fau | 2.56E-09 | 0.195551 | 0.99 | 0.976 | 4.71E-05 |
| Ighm | 2.49E-09 | 0.199094 | 0.656 | 0.457 | 4.59E-05 |
| Slc25a4 | 4.68E-08 | 0.199452 | 0.318 | 0.166 | 0.000861 |
| AB124611 | 1.76E-06 | 0.20102 | 0.142 | 0.049 | 0.032298 |
| Nfkbiz | 4.21E-09 | 0.204359 | 0.616 | 0.419 | 7.75E-05 |
| Smad3 | 5.52E-08 | 0.205148 | 0.185 | 0.072 | 0.001015 |
| Heg1 | 1.37E-06 | 0.205952 | 0.225 | 0.105 | 0.025227 |
| H2afz | 1.74E-06 | 0.206557 | 0.834 | 0.696 | 0.032029 |
| Tspan31 | 2.88E-09 | 0.20752 | 0.238 | 0.098 | 5.30E-05 |
| Ndufv3 | 1.88E-06 | 0.207866 | 0.695 | 0.531 | 0.034588 |
| Supt4a | 3.97E-07 | 0.211092 | 0.649 | 0.471 | 0.007301 |
| Rpl13 | 6.76E-11 | 0.211743 | 0.98 | 0.984 | 1.24E-06 |
| Prelid1 | 1.28E-06 | 0.213039 | 0.785 | 0.634 | 0.023622 |
| Cd69 | 1.00E-07 | 0.213245 | 0.623 | 0.436 | 0.001841 |
| Pdlim1 | 1.64E-08 | 0.213503 | 0.166 | 0.053 | 0.000303 |
| H2afj | 1.15E-08 | 0.21456 | 0.699 | 0.503 | 0.000211 |

| | | | | | |
|---|---|---|---|---|---|
| Rpsa | 1.39E-07 | 0.216065 | 0.993 | 0.963 | 0.002549 |
| H2afx | 1.44E-06 | 0.216513 | 0.209 | 0.092 | 0.026448 |
| Gm8369 | 1.45E-08 | 0.2171 | 0.536 | 0.355 | 0.000267 |
| Ube2d2a | 1.24E-06 | 0.217351 | 0.649 | 0.48 | 0.022734 |
| Bzw2 | 9.07E-11 | 0.217424 | 0.281 | 0.116 | 1.67E-06 |
| Flna | 1.58E-06 | 0.218677 | 0.536 | 0.369 | 0.028982 |
| Sh3bp5 | 6.22E-07 | 0.219503 | 0.136 | 0.046 | 0.011443 |
| Gmfg | 6.10E-07 | 0.219878 | 0.778 | 0.616 | 0.011231 |
| Dgka | 5.54E-07 | 0.220014 | 0.589 | 0.413 | 0.0102 |
| Smc4 | 1.09E-07 | 0.220173 | 0.586 | 0.401 | 0.002007 |
| Foxo1 | 1.63E-07 | 0.221959 | 0.566 | 0.389 | 0.003 |
| Rpl18a | 9.48E-11 | 0.222275 | 0.997 | 0.982 | 1.74E-06 |
| Emd | 6.64E-07 | 0.22251 | 0.437 | 0.272 | 0.012218 |
| Gm10076 | 1.26E-08 | 0.222747 | 0.358 | 0.189 | 0.000232 |
| S1pr4 | 2.79E-09 | 0.224125 | 0.47 | 0.28 | 5.13E-05 |
| Rpl30 | 4.79E-11 | 0.224792 | 0.974 | 0.959 | 8.82E-07 |
| Rexo2 | 1.04E-06 | 0.224921 | 0.464 | 0.302 | 0.019144 |
| Gpr183 | 2.16E-06 | 0.225932 | 0.424 | 0.269 | 0.039746 |
| Rpl24 | 3.15E-10 | 0.226924 | 0.944 | 0.913 | 5.79E-06 |
| Nsa2 | 4.72E-09 | 0.228083 | 0.791 | 0.607 | 8.69E-05 |
| Top1 | 7.97E-08 | 0.231985 | 0.609 | 0.421 | 0.001467 |
| Gclm | 2.60E-06 | 0.23243 | 0.589 | 0.422 | 0.047866 |
| Hipk2 | 1.19E-08 | 0.235121 | 0.232 | 0.097 | 0.000219 |
| Clec2i | 1.92E-08 | 0.237335 | 0.146 | 0.04 | 0.000353 |
| Rpl17 | 2.33E-11 | 0.238039 | 0.98 | 0.94 | 4.29E-07 |
| Fosb | 1.18E-06 | 0.238371 | 0.202 | 0.091 | 0.021792 |
| Serf2 | 6.47E-09 | 0.240088 | 0.934 | 0.872 | 0.000119 |
| Rpl19 | 8.56E-14 | 0.24251 | 0.983 | 0.977 | 1.57E-09 |
| Actb | 7.39E-12 | 0.244056 | 1 | 0.997 | 1.36E-07 |
| Rpl31 | 2.22E-07 | 0.245376 | 0.768 | 0.611 | 0.004091 |
| Eef2 | 3.96E-08 | 0.245972 | 0.934 | 0.862 | 0.000729 |
| Hnrnpa3 | 9.40E-08 | 0.247 | 0.815 | 0.654 | 0.001729 |
| Lockd | 5.08E-09 | 0.247201 | 0.192 | 0.064 | 9.34E-05 |
| Rps27rt | 3.99E-09 | 0.247904 | 0.328 | 0.162 | 7.34E-05 |
| Chd3 | 5.31E-07 | 0.248838 | 0.52 | 0.345 | 0.009777 |
| Cd226 | 4.38E-10 | 0.249279 | 0.212 | 0.074 | 8.07E-06 |
| Rpl23 | 8.31E-14 | 0.249696 | 0.993 | 0.97 | 1.53E-09 |
| Rps24 | 2.41E-13 | 0.251434 | 0.987 | 0.978 | 4.44E-09 |
| Cnn2 | 2.65E-06 | 0.252276 | 0.844 | 0.729 | 0.048684 |
| Cdc25b | 2.32E-07 | 0.253805 | 0.222 | 0.095 | 0.004261 |
| Abhd8 | 1.96E-06 | 0.254267 | 0.311 | 0.17 | 0.036044 |
| Sec61g | 3.57E-08 | 0.254465 | 0.801 | 0.631 | 0.000658 |
| Ndufv2 | 5.65E-07 | 0.254766 | 0.5 | 0.327 | 0.01039 |
| Grk6 | 1.05E-06 | 0.256037 | 0.331 | 0.184 | 0.019237 |
| Rpl12 | 3.86E-08 | 0.257741 | 0.907 | 0.818 | 0.00071 |
| Fas | 1.30E-06 | 0.258943 | 0.205 | 0.089 | 0.023882 |

| | | | | | |
|---|---|---|---|---|---|
| Btf3 | 5.46E-10 | 0.259296 | 0.884 | 0.794 | 1.00E-05 |
| Fyb | 1.13E-08 | 0.259444 | 0.858 | 0.702 | 0.000207 |
| Gm9844 | 1.71E-06 | 0.259688 | 0.315 | 0.174 | 0.031531 |
| Rps17 | 8.83E-07 | 0.260711 | 0.725 | 0.578 | 0.016242 |
| Arhgdib | 4.80E-09 | 0.261261 | 0.934 | 0.843 | 8.82E-05 |
| Tomm7 | 4.51E-08 | 0.261787 | 0.705 | 0.526 | 0.000831 |
| Rps5 | 6.54E-14 | 0.263201 | 0.964 | 0.946 | 1.20E-09 |
| Cox4i1 | 7.48E-12 | 0.263253 | 0.904 | 0.827 | 1.38E-07 |
| Rps8 | 7.36E-13 | 0.263566 | 0.977 | 0.971 | 1.35E-08 |
| Emg1 | 1.20E-08 | 0.265576 | 0.51 | 0.316 | 0.000221 |
| Rflnb | 5.70E-12 | 0.266844 | 0.142 | 0.026 | 1.05E-07 |
| Kdm7a | 3.29E-07 | 0.266857 | 0.325 | 0.173 | 0.006054 |
| Ndufa3 | 1.55E-07 | 0.267325 | 0.672 | 0.495 | 0.002855 |
| Rps23 | 2.80E-13 | 0.267481 | 0.98 | 0.949 | 5.15E-09 |
| H3f3a | 7.62E-10 | 0.268478 | 0.907 | 0.776 | 1.40E-05 |
| Evl | 6.18E-09 | 0.271248 | 0.242 | 0.097 | 0.000114 |
| Arpc3 | 4.17E-08 | 0.271257 | 0.871 | 0.741 | 0.000767 |
| Rasgrp2 | 1.01E-10 | 0.273349 | 0.235 | 0.081 | 1.86E-06 |
| Coq10b | 1.02E-11 | 0.273732 | 0.308 | 0.126 | 1.89E-07 |
| Eif3k | 9.13E-08 | 0.27395 | 0.748 | 0.597 | 0.00168 |
| Cd1d1 | 8.45E-10 | 0.274466 | 0.215 | 0.074 | 1.55E-05 |
| Eef1b2 | 6.10E-10 | 0.275518 | 0.884 | 0.797 | 1.12E-05 |
| Bola2 | 1.86E-09 | 0.275926 | 0.401 | 0.212 | 3.43E-05 |
| Rpl36a | 4.75E-09 | 0.281842 | 0.868 | 0.762 | 8.75E-05 |
| Saraf | 1.06E-07 | 0.283003 | 0.712 | 0.531 | 0.001941 |
| Il18r1 | 9.50E-12 | 0.283503 | 0.331 | 0.145 | 1.75E-07 |
| Rpl21 | 1.33E-13 | 0.283574 | 0.97 | 0.939 | 2.45E-09 |
| Rundc3b | 2.26E-07 | 0.284281 | 0.152 | 0.05 | 0.004149 |
| Jund | 2.16E-07 | 0.285234 | 0.911 | 0.803 | 0.003981 |
| Ndufa6 | 6.05E-08 | 0.286324 | 0.715 | 0.56 | 0.001112 |
| Atp5k | 4.57E-09 | 0.286846 | 0.507 | 0.31 | 8.41E-05 |
| Rpl5 | 2.54E-11 | 0.286905 | 0.907 | 0.752 | 4.68E-07 |
| Hspa1a | 1.34E-06 | 0.287613 | 0.129 | 0.041 | 0.024628 |
| Ppp2r5a | 6.03E-10 | 0.289662 | 0.381 | 0.194 | 1.11E-05 |
| Rpl27a | 4.23E-19 | 0.290366 | 0.97 | 0.954 | 7.78E-15 |
| Rpl34 | 8.47E-16 | 0.291597 | 0.983 | 0.959 | 1.56E-11 |
| Rps9 | 3.44E-17 | 0.292472 | 0.983 | 0.948 | 6.33E-13 |
| Eef1a1 | 1.22E-12 | 0.294412 | 0.99 | 0.98 | 2.25E-08 |
| Slc25a5 | 1.03E-08 | 0.296594 | 0.712 | 0.526 | 0.000189 |
| mt-Atp6 | 1.42E-11 | 0.297032 | 0.964 | 0.914 | 2.62E-07 |
| Tnfrsf25 | 8.92E-10 | 0.297131 | 0.212 | 0.072 | 1.64E-05 |
| Rps13 | 1.05E-17 | 0.297529 | 0.98 | 0.954 | 1.92E-13 |
| Eif3f | 2.03E-10 | 0.297996 | 0.868 | 0.737 | 3.73E-06 |
| Tmsb4x | 3.32E-14 | 0.298161 | 1 | 0.981 | 6.11E-10 |
| Rps3 | 2.19E-16 | 0.299174 | 0.974 | 0.939 | 4.03E-12 |
| Ptp4a2 | 1.63E-08 | 0.300708 | 0.748 | 0.571 | 0.000301 |

| | | | | | |
|---|---|---|---|---|---|
| Rps18 | 1.89E-11 | 0.303457 | 0.924 | 0.823 | 3.47E-07 |
| mt-Nd1 | 4.24E-08 | 0.304403 | 0.831 | 0.679 | 0.000781 |
| Rpl15 | 7.07E-13 | 0.304933 | 0.927 | 0.87 | 1.30E-08 |
| mt-Atp8 | 4.62E-08 | 0.304968 | 0.868 | 0.752 | 0.000851 |
| Rps16 | 6.60E-17 | 0.305442 | 0.993 | 0.971 | 1.21E-12 |
| mt-Nd3 | 1.26E-08 | 0.307163 | 0.404 | 0.224 | 0.000231 |
| Rpl4 | 3.56E-12 | 0.307573 | 0.894 | 0.776 | 6.55E-08 |
| Rps3a1 | 3.39E-18 | 0.308202 | 0.983 | 0.952 | 6.24E-14 |
| Rps10 | 8.96E-19 | 0.308644 | 0.983 | 0.97 | 1.65E-14 |
| Cfl1 | 4.62E-15 | 0.312629 | 0.97 | 0.923 | 8.50E-11 |
| Rpl6 | 2.23E-16 | 0.312842 | 0.954 | 0.917 | 4.10E-12 |
| Gata3 | 2.19E-07 | 0.312945 | 0.5 | 0.323 | 0.004031 |
| Cox7a2l | 4.43E-08 | 0.315636 | 0.656 | 0.478 | 0.000816 |
| Rpl22 | 2.33E-15 | 0.315877 | 0.94 | 0.861 | 4.29E-11 |
| Rps14 | 4.79E-18 | 0.315916 | 0.983 | 0.937 | 8.81E-14 |
| Tcf7 | 4.65E-13 | 0.317675 | 0.219 | 0.06 | 8.56E-09 |
| Twf2 | 7.02E-08 | 0.319326 | 0.467 | 0.291 | 0.001291 |
| B4galnt1 | 1.91E-08 | 0.322739 | 0.447 | 0.265 | 0.000352 |
| Pja1 | 1.34E-08 | 0.326749 | 0.272 | 0.122 | 0.000247 |
| Rpl29 | 3.53E-13 | 0.329751 | 0.94 | 0.868 | 6.49E-09 |
| Coro1a | 1.18E-15 | 0.330949 | 0.98 | 0.899 | 2.17E-11 |
| Stat4 | 3.15E-09 | 0.331178 | 0.483 | 0.285 | 5.80E-05 |
| Naca | 3.08E-17 | 0.33226 | 0.917 | 0.836 | 5.67E-13 |
| Rpl36 | 3.20E-17 | 0.333634 | 0.96 | 0.9 | 5.89E-13 |
| Rps20 | 2.05E-16 | 0.334019 | 0.97 | 0.941 | 3.77E-12 |
| S100a10 | 7.73E-10 | 0.335937 | 0.944 | 0.891 | 1.42E-05 |
| Rps12 | 3.49E-16 | 0.337364 | 0.98 | 0.921 | 6.42E-12 |
| mt-Co1 | 1.11E-25 | 0.339772 | 1 | 0.989 | 2.04E-21 |
| Rpl14 | 2.22E-13 | 0.340058 | 0.894 | 0.812 | 4.09E-09 |
| Rps27a | 1.17E-20 | 0.342509 | 0.977 | 0.957 | 2.16E-16 |
| Prkcq | 7.22E-08 | 0.34431 | 0.503 | 0.319 | 0.001329 |
| Rps4x | 1.51E-20 | 0.346228 | 0.983 | 0.93 | 2.77E-16 |
| Rpl18 | 1.18E-22 | 0.347288 | 0.98 | 0.967 | 2.17E-18 |
| Lat | 8.93E-14 | 0.347406 | 0.858 | 0.658 | 1.64E-09 |
| Tmsb10 | 3.03E-15 | 0.348866 | 0.99 | 0.989 | 5.58E-11 |
| Sub1 | 1.72E-12 | 0.349584 | 0.788 | 0.66 | 3.17E-08 |
| Ppp1r15a | 8.86E-07 | 0.351614 | 0.533 | 0.373 | 0.0163 |
| Ccr4 | 8.59E-10 | 0.353852 | 0.272 | 0.111 | 1.58E-05 |
| Rplp1 | 7.83E-23 | 0.355236 | 0.977 | 0.958 | 1.44E-18 |
| Dnajc15 | 1.18E-09 | 0.356947 | 0.43 | 0.24 | 2.17E-05 |
| Sit1 | 1.76E-09 | 0.357447 | 0.325 | 0.157 | 3.23E-05 |
| Tmem64 | 9.83E-10 | 0.358012 | 0.387 | 0.199 | 1.81E-05 |
| Rpl23a | 6.35E-12 | 0.361316 | 0.851 | 0.687 | 1.17E-07 |
| Rpl10a | 9.80E-20 | 0.364774 | 0.964 | 0.863 | 1.80E-15 |
| Il18rap | 1.32E-12 | 0.366405 | 0.235 | 0.071 | 2.42E-08 |
| Rack1 | 3.02E-16 | 0.36642 | 0.911 | 0.816 | 5.55E-12 |

| | | | | | |
|---|---|---|---|---|---|
| Cxcr4 | 4.24E-09 | 0.367866 | 0.232 | 0.089 | 7.80E-05 |
| mt-Co2 | 5.55E-20 | 0.36799 | 0.99 | 0.946 | 1.02E-15 |
| Actg1 | 2.94E-14 | 0.368257 | 0.997 | 0.992 | 5.40E-10 |
| Cd7 | 1.16E-12 | 0.373191 | 0.295 | 0.11 | 2.14E-08 |
| mt-Co3 | 3.68E-18 | 0.377906 | 0.97 | 0.947 | 6.78E-14 |
| Rplp2 | 1.10E-20 | 0.380992 | 0.964 | 0.901 | 2.03E-16 |
| Sec11c | 6.08E-15 | 0.382566 | 0.725 | 0.479 | 1.12E-10 |
| Rpl26 | 3.36E-20 | 0.384048 | 0.95 | 0.879 | 6.17E-16 |
| Rpl8 | 1.48E-19 | 0.385627 | 0.964 | 0.883 | 2.72E-15 |
| Gm10073 | 3.15E-12 | 0.386676 | 0.291 | 0.119 | 5.79E-08 |
| Klf3 | 1.66E-11 | 0.388234 | 0.235 | 0.077 | 3.05E-07 |
| Adk | 2.26E-09 | 0.388729 | 0.404 | 0.215 | 4.15E-05 |
| Rpl7 | 1.41E-21 | 0.389088 | 0.947 | 0.851 | 2.60E-17 |
| Rpl35 | 4.08E-19 | 0.390926 | 0.924 | 0.844 | 7.51E-15 |
| Klhl6 | 8.08E-13 | 0.391151 | 0.318 | 0.126 | 1.49E-08 |
| Skap1 | 3.10E-14 | 0.393841 | 0.881 | 0.681 | 5.70E-10 |
| Rps6 | 5.17E-20 | 0.394033 | 0.927 | 0.843 | 9.51E-16 |
| Usmg5 | 9.15E-10 | 0.394953 | 0.497 | 0.353 | 1.68E-05 |
| Rps25 | 1.11E-16 | 0.400475 | 0.834 | 0.656 | 2.04E-12 |
| Rps15 | 3.78E-20 | 0.400655 | 0.914 | 0.715 | 6.96E-16 |
| Serpinb1a | 2.29E-10 | 0.401152 | 0.129 | 0.024 | 4.21E-06 |
| Rpl32 | 2.47E-23 | 0.402068 | 0.983 | 0.948 | 4.55E-19 |
| Emp3 | 3.70E-11 | 0.402708 | 0.748 | 0.577 | 6.80E-07 |
| Stat5b | 1.57E-07 | 0.40311 | 0.507 | 0.327 | 0.00289 |
| Tespa1 | 2.31E-15 | 0.411433 | 0.311 | 0.103 | 4.26E-11 |
| Rps26 | 8.28E-24 | 0.42358 | 0.957 | 0.894 | 1.52E-19 |
| Hmgb2 | 4.66E-11 | 0.424496 | 0.785 | 0.581 | 8.58E-07 |
| Rpl38 | 8.79E-28 | 0.429134 | 0.95 | 0.885 | 1.62E-23 |
| Rgs2 | 4.28E-09 | 0.429222 | 0.368 | 0.191 | 7.87E-05 |
| Cd52 | 4.11E-19 | 0.429802 | 0.98 | 0.929 | 7.56E-15 |
| Rps21 | 1.43E-28 | 0.431663 | 0.977 | 0.914 | 2.64E-24 |
| Rpl10 | 1.16E-23 | 0.43259 | 0.927 | 0.82 | 2.13E-19 |
| Ms4a4b | 1.93E-32 | 0.435901 | 0.944 | 0.618 | 3.54E-28 |
| Rpl41 | 1.34E-32 | 0.441959 | 0.983 | 0.965 | 2.47E-28 |
| Ramp3 | 5.67E-11 | 0.44617 | 0.182 | 0.05 | 1.04E-06 |
| Cd40lg | 1.92E-14 | 0.446368 | 0.238 | 0.064 | 3.53E-10 |
| Rplp0 | 1.83E-25 | 0.4471 | 0.977 | 0.943 | 3.38E-21 |
| Il1rl1 | 3.53E-11 | 0.456121 | 0.255 | 0.092 | 6.50E-07 |
| S1pr1 | 1.21E-14 | 0.461085 | 0.338 | 0.128 | 2.23E-10 |
| Ms4a6b | 1.48E-19 | 0.463383 | 0.848 | 0.618 | 2.72E-15 |
| Rps19 | 4.20E-29 | 0.471194 | 0.96 | 0.912 | 7.73E-25 |
| Gramd3 | 8.14E-11 | 0.476776 | 0.387 | 0.189 | 1.50E-06 |
| Rpl37a | 1.09E-36 | 0.481512 | 0.97 | 0.937 | 2.00E-32 |
| Rpl35a | 8.66E-34 | 0.481852 | 0.97 | 0.922 | 1.59E-29 |
| Rpl37 | 1.59E-36 | 0.482124 | 0.967 | 0.93 | 2.93E-32 |
| Ass1 | 2.04E-14 | 0.488963 | 0.712 | 0.484 | 3.75E-10 |

| | | | | | |
|---|---|---|---|---|---|
| Ier2 | 1.29E-12 | 0.497331 | 0.626 | 0.407 | 2.37E-08 |
| Btg2 | 2.90E-16 | 0.500797 | 0.636 | 0.367 | 5.33E-12 |
| Slamf6 | 6.08E-17 | 0.501393 | 0.411 | 0.168 | 1.12E-12 |
| Tox | 1.36E-06 | 0.508854 | 0.424 | 0.266 | 0.025056 |
| Junb | 2.62E-16 | 0.51257 | 0.901 | 0.774 | 4.81E-12 |
| Zfp36 | 3.43E-11 | 0.513087 | 0.725 | 0.524 | 6.30E-07 |
| Cldnd1 | 7.10E-15 | 0.518733 | 0.381 | 0.161 | 1.31E-10 |
| Trat1 | 4.69E-21 | 0.526012 | 0.377 | 0.117 | 8.62E-17 |
| Rpl28 | 9.44E-33 | 0.536037 | 0.97 | 0.888 | 1.74E-28 |
| Cpm | 2.83E-11 | 0.542718 | 0.331 | 0.145 | 5.21E-07 |
| Itgb1 | 4.14E-18 | 0.560816 | 0.579 | 0.294 | 7.62E-14 |
| Rps11 | 2.11E-46 | 0.561015 | 0.983 | 0.95 | 3.88E-42 |
| Nrp1 | 2.58E-23 | 0.572453 | 0.334 | 0.081 | 4.75E-19 |
| Rpl39 | 3.19E-42 | 0.579773 | 0.97 | 0.937 | 5.88E-38 |
| Sh2d1a | 1.46E-17 | 0.580624 | 0.513 | 0.267 | 2.69E-13 |
| St6gal1 | 1.84E-24 | 0.58446 | 0.338 | 0.079 | 3.39E-20 |
| Rps28 | 1.54E-40 | 0.591976 | 0.937 | 0.824 | 2.84E-36 |
| Rps29 | 8.97E-46 | 0.594134 | 0.96 | 0.881 | 1.65E-41 |
| Rps27 | 3.85E-46 | 0.601168 | 0.957 | 0.913 | 7.08E-42 |
| Fos | 1.31E-16 | 0.640949 | 0.579 | 0.305 | 2.42E-12 |
| Klf2 | 1.21E-25 | 0.801081 | 0.632 | 0.296 | 2.22E-21 |
| Emb | 4.49E-48 | 0.971461 | 0.732 | 0.277 | 8.25E-44 |

**Supplementary Table 3.** Genes differentially expressed between tumor-infiltrating TEFF from untreated and aCTLA-4 treated mice

| Gene Symbol | p_val | avg_logFC | pct.1 | pct.2 | p_val_adj |
|---|---|---|---|---|---|
| Lag3 | 2.79E-57 | -0.7580391 | 0.138 | 0.315 | 5.14E-53 |
| Ccr8 | 2.44E-31 | -0.5232792 | 0.203 | 0.34 | 4.48E-27 |
| Rgs16 | 8.98E-19 | -0.5199569 | 0.123 | 0.217 | 1.65E-14 |
| Ccl4 | 3.04E-09 | -0.5134999 | 0.155 | 0.119 | 5.60E-05 |
| Ikzf2 | 1.09E-16 | -0.4343162 | 0.153 | 0.241 | 2.01E-12 |
| Ccr7 | 1.02E-10 | -0.4128753 | 0.079 | 0.13 | 1.87E-06 |
| Ctla4 | 1.42E-20 | -0.3931659 | 0.486 | 0.526 | 2.61E-16 |
| Pdcd1 | 1.30E-21 | -0.3666306 | 0.406 | 0.445 | 2.39E-17 |
| Tnfsf11 | 1.23E-09 | -0.3365181 | 0.274 | 0.346 | 2.27E-05 |
| St6galnac3 | 5.63E-07 | -0.3360138 | 0.23 | 0.299 | 0.010352 |
| Tigit | 9.00E-17 | -0.3175726 | 0.349 | 0.425 | 1.66E-12 |
| Eea1 | 2.97E-10 | -0.3166226 | 0.223 | 0.268 | 5.46E-06 |
| Tox | 1.49E-07 | -0.315184 | 0.139 | 0.201 | 0.0027394 |
| Vps37b | 1.92E-13 | -0.2965628 | 0.511 | 0.593 | 3.53E-09 |
| Isy1 | 3.09E-15 | -0.289549 | 0.292 | 0.373 | 5.69E-11 |
| Rps7 | 8.15E-58 | -0.2877563 | 0.98 | 0.975 | 1.50E-53 |
| Zfp36l1 | 3.26E-13 | -0.286593 | 0.535 | 0.602 | 6.00E-09 |
| Rabgap1l | 1.28E-06 | -0.2831017 | 0.401 | 0.447 | 0.0236354 |
| Foxp1 | 1.51E-08 | -0.2669914 | 0.255 | 0.319 | 0.0002772 |
| Trbv3 | 2.19E-07 | -0.2657209 | 0.207 | 0.209 | 0.0040251 |
| Cirbp | 7.62E-16 | -0.2597297 | 0.294 | 0.399 | 1.40E-11 |
| Irf7 | 3.05E-09 | -0.2592606 | 0.395 | 0.471 | 5.62E-05 |
| Tnfrsf4 | 4.77E-14 | -0.2586494 | 0.707 | 0.708 | 8.78E-10 |
| Phlpp1 | 5.60E-08 | -0.2571264 | 0.086 | 0.11 | 0.0010311 |
| Gdi2 | 4.51E-25 | -0.2566614 | 0.697 | 0.714 | 8.31E-21 |
| Cd27 | 1.89E-13 | -0.2526101 | 0.15 | 0.247 | 3.48E-09 |
| Pcbp2 | 9.76E-47 | -0.2525552 | 0.722 | 0.716 | 1.80E-42 |
| Sp100 | 4.39E-18 | -0.2508107 | 0.59 | 0.655 | 8.08E-14 |
| Itgav | 3.34E-09 | -0.2495025 | 0.167 | 0.205 | 6.14E-05 |
| Ubash3b | 6.45E-10 | -0.2467593 | 0.113 | 0.157 | 1.19E-05 |
| Izumo1r | 2.46E-11 | -0.2453128 | 0.093 | 0.169 | 4.53E-07 |
| Gimap6 | 1.06E-12 | -0.2444965 | 0.58 | 0.638 | 1.95E-08 |
| Txnip | 4.45E-10 | -0.2415532 | 0.4 | 0.474 | 8.18E-06 |
| Prkch | 3.38E-09 | -0.2390002 | 0.423 | 0.494 | 6.22E-05 |
| Dusp4 | 2.97E-12 | -0.2381556 | 0.159 | 0.197 | 5.47E-08 |
| Gimap3 | 5.97E-19 | -0.2378944 | 0.735 | 0.749 | 1.10E-14 |
| Ddx5 | 7.64E-34 | -0.234842 | 0.893 | 0.893 | 1.41E-29 |
| Inpp4b | 2.65E-12 | -0.2346509 | 0.523 | 0.639 | 4.87E-08 |
| Arhgap45 | 8.31E-20 | -0.2331156 | 0.696 | 0.766 | 1.53E-15 |
| Ypel3 | 1.62E-12 | -0.2331117 | 0.531 | 0.602 | 2.98E-08 |
| Gapdh | 2.11E-28 | -0.231011 | 0.829 | 0.803 | 3.88E-24 |
| Ikzf3 | 6.32E-10 | -0.2302878 | 0.116 | 0.192 | 1.16E-05 |

| | | | | | |
|---|---|---|---|---|---|
| Gm11808 | 4.86E-17 | -0.221917 | 0.468 | 0.533 | 8.94E-13 |
| Cd3e | 1.83E-41 | -0.2186401 | 1 | 1 | 3.38E-37 |
| Mdfic | 5.72E-10 | -0.2151291 | 0.283 | 0.344 | 1.05E-05 |
| Tnfrsf18 | 2.53E-12 | -0.2137578 | 0.77 | 0.8 | 4.66E-08 |
| Gpm6b | 2.26E-06 | -0.2130087 | 0.085 | 0.131 | 0.0415463 |
| Bcl2a1d | 2.19E-11 | -0.2126574 | 0.359 | 0.422 | 4.03E-07 |
| Mrps6 | 4.60E-10 | -0.2107682 | 0.125 | 0.192 | 8.47E-06 |
| Tnfrsf1b | 1.33E-07 | -0.2090381 | 0.259 | 0.327 | 0.0024389 |
| Sla | 3.59E-09 | -0.2043596 | 0.519 | 0.554 | 6.61E-05 |
| AC149090.1 | 3.95E-10 | -0.2039038 | 0.175 | 0.258 | 7.26E-06 |
| Slfn1 | 1.34E-09 | -0.1992561 | 0.541 | 0.588 | 2.46E-05 |
| Shisa5 | 1.16E-34 | -0.1975436 | 0.964 | 0.937 | 2.13E-30 |
| Tg | 3.95E-10 | -0.1958961 | 0.104 | 0.12 | 7.28E-06 |
| Gstp1 | 2.10E-12 | -0.1940826 | 0.481 | 0.545 | 3.86E-08 |
| Grcc10 | 1.98E-11 | -0.1916367 | 0.572 | 0.645 | 3.64E-07 |
| Gm20400 | 4.39E-13 | -0.1915696 | 0.074 | 0.139 | 8.08E-09 |
| Il4ra | 3.64E-08 | -0.191031 | 0.138 | 0.187 | 0.0006691 |
| Chchd2 | 1.77E-16 | -0.1878183 | 0.586 | 0.646 | 3.26E-12 |
| Igf2r | 9.36E-09 | -0.1872471 | 0.071 | 0.124 | 0.0001721 |
| Klrd1 | 2.04E-07 | -0.1835932 | 0.406 | 0.404 | 0.003756 |
| Il2rb | 1.85E-18 | -0.1835111 | 0.771 | 0.753 | 3.41E-14 |
| Eno1 | 1.45E-18 | -0.1831034 | 0.7 | 0.661 | 2.67E-14 |
| Cd2 | 1.24E-20 | -0.1829195 | 0.871 | 0.862 | 2.29E-16 |
| Dhx58 | 2.46E-11 | -0.1809776 | 0.091 | 0.168 | 4.52E-07 |
| Sap18 | 5.10E-11 | -0.1809 | 0.417 | 0.483 | 9.39E-07 |
| Col4a3bp | 1.47E-06 | -0.1760792 | 0.161 | 0.197 | 0.0270185 |
| Snx2 | 1.46E-07 | -0.1753906 | 0.273 | 0.326 | 0.0026794 |
| Ap3b1 | 1.03E-06 | -0.1740107 | 0.248 | 0.288 | 0.0190321 |
| Crlf2 | 3.50E-11 | -0.17398 | 0.474 | 0.494 | 6.44E-07 |
| Arl5a | 3.27E-09 | -0.1734587 | 0.123 | 0.142 | 6.01E-05 |
| mt-Nd4l | 8.51E-11 | -0.1731584 | 0.828 | 0.858 | 1.56E-06 |
| Bcl2a1b | 2.51E-07 | -0.1721698 | 0.488 | 0.549 | 0.0046093 |
| Cd3g | 5.34E-24 | -0.1720973 | 0.944 | 0.921 | 9.83E-20 |
| Cnbp | 7.85E-19 | -0.1719904 | 0.78 | 0.791 | 1.44E-14 |
| Camk2n1 | 2.20E-06 | -0.171768 | 0.117 | 0.16 | 0.0404602 |
| Rgs10 | 1.04E-06 | -0.1682724 | 0.333 | 0.393 | 0.0190551 |
| Trim30a | 4.10E-08 | -0.1674531 | 0.36 | 0.382 | 0.0007541 |
| Nfatc1 | 1.96E-06 | -0.1656406 | 0.326 | 0.375 | 0.0360864 |
| Gimap1 | 2.72E-10 | -0.1649816 | 0.64 | 0.68 | 5.00E-06 |
| Icos | 8.58E-12 | -0.1642486 | 0.811 | 0.786 | 1.58E-07 |
| Cd4 | 1.29E-14 | -0.1619628 | 1 | 1 | 2.37E-10 |
| Sdcbp2 | 1.69E-06 | -0.1617411 | 0.126 | 0.17 | 0.0310293 |
| Ccdc85b | 9.38E-08 | -0.1594793 | 0.142 | 0.201 | 0.0017253 |
| Sh2d2a | 4.64E-10 | -0.1594655 | 0.65 | 0.666 | 8.54E-06 |
| Itm2c | 1.94E-08 | -0.1592851 | 0.514 | 0.562 | 0.0003565 |

| | | | | | |
|---|---|---|---|---|---|
| Hnrnpk | 2.40E-15 | -0.1592606 | 0.717 | 0.738 | 4.42E-11 |
| Gps2 | 6.38E-07 | -0.1565106 | 0.24 | 0.315 | 0.0117428 |
| Lbh | 7.57E-08 | -0.1564259 | 0.363 | 0.374 | 0.0013923 |
| Ctsb | 2.87E-08 | -0.1560912 | 0.548 | 0.577 | 0.0005275 |
| Ext1 | 1.38E-09 | -0.1528784 | 0.111 | 0.1 | 2.53E-05 |
| Slc1a5 | 3.30E-07 | -0.1519521 | 0.353 | 0.4 | 0.0060704 |
| Cyth4 | 7.53E-07 | -0.151619 | 0.376 | 0.436 | 0.0138585 |
| Ctsw | 2.47E-15 | -0.1502067 | 0.572 | 0.538 | 4.55E-11 |
| Gdi1 | 2.93E-07 | -0.1474704 | 0.197 | 0.273 | 0.0053871 |
| Jak3 | 1.61E-06 | -0.1471883 | 0.363 | 0.413 | 0.0295752 |
| Psap | 4.39E-08 | -0.1470627 | 0.48 | 0.514 | 0.0008074 |
| Rasal3 | 3.62E-07 | -0.1465539 | 0.445 | 0.494 | 0.0066654 |
| Ftl1 | 3.24E-16 | -0.1450377 | 0.855 | 0.838 | 5.96E-12 |
| Pink1 | 5.01E-08 | -0.1401991 | 0.187 | 0.268 | 0.0009217 |
| Ubb | 2.55E-16 | -0.1397779 | 0.978 | 0.972 | 4.69E-12 |
| Adck5 | 2.84E-10 | -0.1383614 | 0.068 | 0.132 | 5.22E-06 |
| Lrrc8d | 6.54E-07 | -0.1368252 | 0.155 | 0.152 | 0.0120332 |
| Capg | 2.30E-07 | -0.1337718 | 0.624 | 0.616 | 0.004232 |
| Sv2c | 2.29E-06 | -0.1336192 | 0.091 | 0.114 | 0.0421833 |
| Malat1 | 3.57E-11 | -0.1332934 | 0.982 | 0.995 | 6.57E-07 |
| Arhgef1 | 2.08E-09 | -0.1316085 | 0.65 | 0.662 | 3.82E-05 |
| Lime1 | 7.41E-07 | -0.1315121 | 0.114 | 0.172 | 0.0136406 |
| Msi2 | 1.60E-06 | -0.1302825 | 0.221 | 0.213 | 0.0294894 |
| Sumo3 | 1.96E-06 | -0.1300569 | 0.07 | 0.112 | 0.0359801 |
| Hmgb1 | 1.11E-07 | -0.1293484 | 0.741 | 0.778 | 0.0020398 |
| H2-D1 | 8.95E-10 | -0.1226131 | 0.997 | 0.989 | 1.65E-05 |
| Ifi27l2a | 1.54E-13 | -0.1216847 | 0.834 | 0.904 | 2.83E-09 |
| mt-Atp8 | 1.73E-11 | -0.1205522 | 0.898 | 0.878 | 3.18E-07 |
| Prkar1a | 1.97E-16 | -0.1202917 | 0.785 | 0.761 | 3.63E-12 |
| Arpc1b | 3.66E-16 | -0.1201659 | 0.885 | 0.859 | 6.73E-12 |
| H3f3b | 1.19E-14 | -0.1196566 | 0.979 | 0.961 | 2.20E-10 |
| Nt5e | 1.99E-06 | -0.1191125 | 0.086 | 0.137 | 0.0366731 |
| H2-K1 | 3.67E-11 | -0.1184888 | 0.998 | 0.994 | 6.75E-07 |
| Stk17b | 5.97E-08 | -0.1182882 | 0.843 | 0.845 | 0.0010988 |
| Slfn2 | 2.24E-07 | -0.1170862 | 0.784 | 0.788 | 0.0041228 |
| Ppp1r18 | 5.28E-17 | -0.1166426 | 0.737 | 0.705 | 9.72E-13 |
| Acap1 | 5.14E-07 | -0.1164884 | 0.455 | 0.482 | 0.0094516 |
| Tmem50a | 2.44E-11 | -0.1155761 | 0.719 | 0.717 | 4.49E-07 |
| Tut4 | 1.82E-13 | -0.1153852 | 0.17 | 0.148 | 3.36E-09 |
| Atp5o | 1.27E-14 | -0.1119706 | 0.158 | 0.14 | 2.34E-10 |
| Laptm5 | 2.29E-13 | -0.1119031 | 0.948 | 0.926 | 4.21E-09 |
| 1810026B05Rik | 1.12E-14 | -0.1113108 | 0.115 | 0.1 | 2.06E-10 |
| Ctdnep1 | 2.13E-08 | -0.1098832 | 0.131 | 0.134 | 0.0003921 |
| Ndfip1 | 1.84E-07 | -0.1084988 | 0.89 | 0.887 | 0.0033892 |
| Ywhaz | 9.40E-16 | -0.1082378 | 0.877 | 0.847 | 1.73E-11 |

| | | | | | |
|---|---|---|---|---|---|
| Lrp10 | 8.86E-08 | -0.107985 | 0.547 | 0.552 | 0.0016301 |
| Phf1 | 2.17E-06 | -0.1061345 | 0.136 | 0.197 | 0.0399024 |
| Hnrnpa3 | 6.79E-07 | -0.1049799 | 0.786 | 0.802 | 0.0124926 |
| Msn | 2.81E-07 | -0.1025912 | 0.821 | 0.826 | 0.0051702 |
| Atp5e | 2.15E-06 | 0.1006896 | 0.906 | 0.88 | 0.0395647 |
| Gls | 1.02E-08 | 0.1015294 | 0.27 | 0.189 | 0.0001879 |
| Aurkaip1 | 2.32E-06 | 0.1024501 | 0.531 | 0.445 | 0.0426851 |
| Reep5 | 1.73E-08 | 0.1028242 | 0.759 | 0.669 | 0.0003184 |
| Klf2 | 2.53E-06 | 0.1029781 | 0.369 | 0.291 | 0.0466035 |
| Atp5j2 | 1.12E-06 | 0.1049993 | 0.831 | 0.774 | 0.020671 |
| B2m | 3.92E-11 | 0.1068982 | 0.997 | 0.986 | 7.21E-07 |
| Sec11c | 1.82E-06 | 0.1073305 | 0.573 | 0.486 | 0.0335682 |
| Rpl26 | 7.33E-08 | 0.1094473 | 0.974 | 0.955 | 0.0013489 |
| Timm8b | 1.30E-06 | 0.1102767 | 0.32 | 0.24 | 0.0239413 |
| Cks1b | 5.50E-07 | 0.1107261 | 0.108 | 0.058 | 0.0101123 |
| Uqcr11 | 8.26E-07 | 0.1116721 | 0.529 | 0.44 | 0.0152013 |
| Gng2 | 3.20E-07 | 0.1125515 | 0.655 | 0.565 | 0.0058887 |
| Snhg6 | 9.65E-08 | 0.1126065 | 0.2 | 0.129 | 0.0017755 |
| Psmb1 | 7.65E-07 | 0.113132 | 0.773 | 0.701 | 0.0140757 |
| Rpl34 | 1.76E-10 | 0.1133679 | 0.983 | 0.982 | 3.23E-06 |
| mt-Co1 | 2.95E-11 | 0.1141842 | 0.998 | 0.996 | 5.43E-07 |
| Rps6 | 1.53E-08 | 0.1143666 | 0.959 | 0.937 | 0.000281 |
| Btf3 | 1.83E-07 | 0.1143723 | 0.919 | 0.88 | 0.0033608 |
| Minos1 | 4.02E-08 | 0.1144515 | 0.386 | 0.37 | 0.0007391 |
| Kif2a | 4.21E-11 | 0.1148178 | 0.185 | 0.109 | 7.75E-07 |
| 1810037I17 Rik | 7.59E-07 | 0.1148793 | 0.714 | 0.637 | 0.0139729 |
| Arpp19 | 2.68E-06 | 0.1151097 | 0.625 | 0.544 | 0.0493273 |
| Rps19 | 4.52E-11 | 0.1166467 | 0.986 | 0.974 | 8.31E-07 |
| Rpl7a | 3.36E-10 | 0.1179879 | 0.962 | 0.934 | 6.17E-06 |
| Serf2 | 5.69E-10 | 0.1183518 | 0.94 | 0.919 | 1.05E-05 |
| Cenpw | 1.32E-08 | 0.1186066 | 0.141 | 0.077 | 0.0002425 |
| Rpl32 | 7.72E-10 | 0.1209984 | 0.99 | 0.979 | 1.42E-05 |
| Rack1 | 1.41E-10 | 0.1223046 | 0.958 | 0.931 | 2.59E-06 |
| Atp5k | 1.55E-07 | 0.1229004 | 0.477 | 0.384 | 0.002854 |
| Gpi1 | 6.14E-09 | 0.1237805 | 0.811 | 0.724 | 0.0001129 |
| Cox7a2 | 2.41E-07 | 0.124037 | 0.753 | 0.71 | 0.0044351 |
| Ptp4a2 | 4.84E-07 | 0.124283 | 0.664 | 0.58 | 0.0089041 |
| Gnai2 | 5.55E-09 | 0.1252438 | 0.754 | 0.673 | 0.0001022 |
| Rpl28 | 1.37E-10 | 0.1267289 | 0.984 | 0.972 | 2.52E-06 |
| Hist1h1b | 1.94E-08 | 0.1274236 | 0.134 | 0.079 | 0.0003576 |
| Ndufa6 | 9.62E-08 | 0.1276294 | 0.764 | 0.686 | 0.0017691 |
| Sem1 | 2.31E-09 | 0.1308311 | 0.829 | 0.783 | 4.26E-05 |
| Csnk2b | 1.09E-07 | 0.1330978 | 0.762 | 0.684 | 0.0020069 |
| Sh3bgrl3 | 9.17E-09 | 0.135386 | 0.961 | 0.937 | 0.0001687 |
| Eloc | 1.33E-07 | 0.1362668 | 0.556 | 0.462 | 0.0024443 |

| | | | | | |
|---|---|---|---|---|---|
| Nfkbia | 2.17E-07 | 0.1377577 | 0.77 | 0.698 | 0.0039844 |
| Cd52 | 1.27E-09 | 0.1380518 | 0.982 | 0.962 | 2.34E-05 |
| Tab2 | 4.81E-12 | 0.1384375 | 0.359 | 0.25 | 8.85E-08 |
| Dpm3 | 6.34E-07 | 0.1384837 | 0.42 | 0.335 | 0.0116617 |
| Rplp2 | 2.40E-15 | 0.1388035 | 0.98 | 0.962 | 4.42E-11 |
| Ndufv3 | 2.73E-08 | 0.139244 | 0.7 | 0.616 | 0.0005031 |
| Adam8 | 1.06E-08 | 0.1396152 | 0.236 | 0.154 | 0.0001952 |
| Cyba | 5.76E-12 | 0.1398427 | 0.916 | 0.854 | 1.06E-07 |
| Rps15 | 1.87E-11 | 0.1410843 | 0.924 | 0.878 | 3.44E-07 |
| Tmem256 | 5.42E-08 | 0.1410921 | 0.34 | 0.25 | 0.0009966 |
| Rps17 | 5.14E-09 | 0.1412131 | 0.773 | 0.706 | 9.46E-05 |
| Pfkp | 5.91E-09 | 0.1414871 | 0.425 | 0.323 | 0.0001087 |
| Cox5a | 2.03E-09 | 0.1433815 | 0.8 | 0.72 | 3.73E-05 |
| Cox8a | 3.20E-12 | 0.1434985 | 0.937 | 0.9 | 5.88E-08 |
| Cox4i1 | 4.47E-15 | 0.143611 | 0.94 | 0.888 | 8.23E-11 |
| Gpr183 | 2.86E-10 | 0.1438069 | 0.53 | 0.419 | 5.27E-06 |
| S1pr4 | 1.84E-06 | 0.1438156 | 0.373 | 0.291 | 0.0338249 |
| Glipr2 | 1.73E-08 | 0.1443138 | 0.522 | 0.421 | 0.0003175 |
| Rpl36 | 3.52E-14 | 0.1455488 | 0.973 | 0.962 | 6.48E-10 |
| S100a11 | 2.06E-07 | 0.1465407 | 0.867 | 0.815 | 0.003799 |
| Rpl24 | 9.86E-16 | 0.1477488 | 0.98 | 0.967 | 1.81E-11 |
| Nabp1 | 2.54E-06 | 0.1481945 | 0.433 | 0.354 | 0.0467488 |
| Ltb4r1 | 2.15E-08 | 0.1486314 | 0.121 | 0.063 | 0.0003958 |
| Lockd | 7.93E-09 | 0.1491434 | 0.149 | 0.083 | 0.0001459 |
| Klk8 | 3.81E-07 | 0.1493259 | 0.378 | 0.291 | 0.007008 |
| Emb | 2.09E-13 | 0.1540898 | 0.749 | 0.633 | 3.84E-09 |
| Rpl31 | 3.76E-13 | 0.1557469 | 0.832 | 0.773 | 6.92E-09 |
| Cks2 | 1.67E-06 | 0.1560155 | 0.33 | 0.253 | 0.0308167 |
| Higd1a | 2.12E-08 | 0.157563 | 0.602 | 0.506 | 0.0003906 |
| Gm10076 | 6.09E-10 | 0.1582528 | 0.41 | 0.305 | 1.12E-05 |
| Fam162a | 4.35E-07 | 0.1592432 | 0.382 | 0.303 | 0.0079948 |
| Hint1 | 1.04E-11 | 0.1595719 | 0.798 | 0.716 | 1.91E-07 |
| Rps25 | 1.08E-12 | 0.1603429 | 0.884 | 0.838 | 1.99E-08 |
| Mrpl33 | 5.67E-11 | 0.1619233 | 0.695 | 0.596 | 1.04E-06 |
| Selenoh | 8.95E-08 | 0.165129 | 0.31 | 0.227 | 0.0016466 |
| Gmfg | 1.23E-11 | 0.167897 | 0.775 | 0.677 | 2.27E-07 |
| Atp2b1 | 9.34E-08 | 0.1702134 | 0.452 | 0.358 | 0.0017177 |
| Lgals7 | 1.19E-07 | 0.1715249 | 0.267 | 0.186 | 0.0021934 |
| mt-Nd3 | 2.21E-07 | 0.1722663 | 0.403 | 0.32 | 0.0040721 |
| Rpl22l1 | 1.10E-12 | 0.1732471 | 0.856 | 0.791 | 2.02E-08 |
| Cox6a1 | 7.08E-12 | 0.1759541 | 0.73 | 0.642 | 1.30E-07 |
| Emp3 | 1.02E-09 | 0.1764639 | 0.653 | 0.556 | 1.87E-05 |
| Cops9 | 4.32E-11 | 0.1776784 | 0.593 | 0.493 | 7.96E-07 |
| Slamf7 | 9.61E-10 | 0.1777595 | 0.252 | 0.163 | 1.77E-05 |
| Rps21 | 1.27E-24 | 0.1814046 | 0.976 | 0.974 | 2.34E-20 |
| Iqgap2 | 1.34E-09 | 0.1824622 | 0.308 | 0.213 | 2.46E-05 |

| | | | | | |
|---|---|---|---|---|---|
| Rps27 | 1.89E-13 | 0.1833866 | 0.981 | 0.965 | 3.47E-09 |
| Klf6 | 1.23E-07 | 0.1841607 | 0.635 | 0.543 | 0.0022702 |
| Actg1 | 1.43E-14 | 0.1873159 | 0.997 | 0.989 | 2.63E-10 |
| Tagln2 | 9.02E-12 | 0.187382 | 0.868 | 0.797 | 1.66E-07 |
| Tmsb10 | 8.57E-20 | 0.1875638 | 0.999 | 0.995 | 1.58E-15 |
| Rps26 | 4.01E-21 | 0.1876225 | 0.985 | 0.974 | 7.38E-17 |
| Il1r2 | 1.54E-07 | 0.188581 | 0.218 | 0.144 | 0.0028295 |
| Txn1 | 2.21E-12 | 0.1899691 | 0.758 | 0.674 | 4.07E-08 |
| Tomm20 | 2.21E-12 | 0.1907368 | 0.668 | 0.562 | 4.07E-08 |
| Rpl23a | 9.29E-17 | 0.1915175 | 0.893 | 0.838 | 1.71E-12 |
| Atp5l | 1.32E-19 | 0.1916931 | 0.883 | 0.843 | 2.44E-15 |
| S100a13 | 2.21E-13 | 0.1934554 | 0.782 | 0.687 | 4.07E-09 |
| Bola2 | 3.27E-12 | 0.1951523 | 0.417 | 0.299 | 6.02E-08 |
| Il18rap | 6.66E-11 | 0.1954098 | 0.35 | 0.244 | 1.23E-06 |
| Tnp2 | 3.00E-09 | 0.197758 | 0.157 | 0.101 | 5.51E-05 |
| Rps29 | 1.56E-26 | 0.1978246 | 0.981 | 0.958 | 2.87E-22 |
| Prelid1 | 1.13E-13 | 0.1986036 | 0.809 | 0.723 | 2.09E-09 |
| Cox6c | 1.96E-19 | 0.2030928 | 0.866 | 0.789 | 3.61E-15 |
| Slc2a3 | 8.52E-08 | 0.203847 | 0.251 | 0.177 | 0.0015678 |
| Ndufa3 | 4.57E-16 | 0.2137659 | 0.716 | 0.61 | 8.41E-12 |
| Ifitm1 | 1.46E-22 | 0.2144883 | 0.503 | 0.337 | 2.68E-18 |
| Gclm | 5.40E-15 | 0.214759 | 0.632 | 0.521 | 9.93E-11 |
| Nfkbiz | 5.88E-10 | 0.2160051 | 0.518 | 0.413 | 1.08E-05 |
| Rpl39 | 9.13E-27 | 0.2184903 | 0.982 | 0.97 | 1.68E-22 |
| Cd40lg | 3.05E-16 | 0.2187102 | 0.575 | 0.433 | 5.61E-12 |
| S100a10 | 1.12E-15 | 0.2232457 | 0.931 | 0.893 | 2.07E-11 |
| Cox7c | 1.05E-21 | 0.2256704 | 0.87 | 0.78 | 1.94E-17 |
| Stmn1 | 2.38E-06 | 0.2281783 | 0.16 | 0.102 | 0.0438684 |
| Pclaf | 1.58E-08 | 0.2291802 | 0.111 | 0.055 | 0.0002908 |
| Gm10073 | 3.21E-13 | 0.2292589 | 0.311 | 0.224 | 5.90E-09 |
| Rpl37a | 1.68E-35 | 0.2299472 | 0.988 | 0.972 | 3.09E-31 |
| Tpi1 | 2.03E-11 | 0.2316825 | 0.649 | 0.559 | 3.73E-07 |
| Tomm7 | 9.24E-22 | 0.2320229 | 0.759 | 0.667 | 1.70E-17 |
| Krtcap2 | 2.90E-22 | 0.2335857 | 0.804 | 0.728 | 5.34E-18 |
| Anxa2 | 1.46E-09 | 0.2342959 | 0.667 | 0.594 | 2.69E-05 |
| Ero1l | 5.51E-10 | 0.2385159 | 0.313 | 0.216 | 1.01E-05 |
| Rpl41 | 9.81E-39 | 0.2422146 | 0.994 | 0.986 | 1.80E-34 |
| Id2 | 4.55E-20 | 0.2479305 | 0.814 | 0.688 | 8.38E-16 |
| Pgk1 | 6.07E-15 | 0.2543244 | 0.744 | 0.655 | 1.12E-10 |
| Rps28 | 4.53E-39 | 0.2546286 | 0.969 | 0.943 | 8.33E-35 |
| Cd69 | 1.96E-14 | 0.2557888 | 0.616 | 0.488 | 3.60E-10 |
| Gadd45g | 5.80E-11 | 0.2561304 | 0.164 | 0.091 | 1.07E-06 |
| Mki67 | 7.70E-07 | 0.258359 | 0.133 | 0.078 | 0.0141695 |
| Rpl37 | 1.54E-49 | 0.2664032 | 0.983 | 0.976 | 2.84E-45 |
| 2010107E04Rik | 3.04E-23 | 0.2706229 | 0.538 | 0.481 | 5.60E-19 |

| | | | | | |
|---|---|---|---|---|---|
| Aldoa | 1.66E-17 | 0.2708488 | 0.91 | 0.843 | 3.05E-13 |
| Usmg5 | 4.94E-27 | 0.2760464 | 0.56 | 0.502 | 9.09E-23 |
| Lgals1 | 4.25E-18 | 0.2868518 | 0.916 | 0.895 | 7.82E-14 |
| Rpl38 | 2.07E-56 | 0.289361 | 0.971 | 0.957 | 3.81E-52 |
| Vim | 1.86E-19 | 0.2936983 | 0.876 | 0.791 | 3.43E-15 |
| Serpinb6b | 2.75E-17 | 0.295358 | 0.456 | 0.312 | 5.06E-13 |
| Sec61g | 4.59E-37 | 0.31167 | 0.819 | 0.699 | 8.44E-33 |
| Hmgb2 | 1.46E-10 | 0.3151823 | 0.741 | 0.677 | 2.68E-06 |
| Lgals3 | 1.73E-08 | 0.3182595 | 0.488 | 0.406 | 0.0003187 |
| Cdkn1a | 1.97E-19 | 0.3282755 | 0.323 | 0.185 | 3.62E-15 |
| AA467197 | 8.03E-18 | 0.3308997 | 0.322 | 0.192 | 1.48E-13 |
| Mif | 6.79E-28 | 0.3435424 | 0.825 | 0.741 | 1.25E-23 |
| Ifitm3 | 2.81E-15 | 0.3575316 | 0.663 | 0.539 | 5.18E-11 |
| Furin | 5.99E-24 | 0.3577732 | 0.449 | 0.287 | 1.10E-19 |
| Crip1 | 7.30E-31 | 0.4553203 | 0.91 | 0.842 | 1.34E-26 |
| Ifng | 3.32E-08 | 0.6383384 | 0.325 | 0.237 | 0.0006106 |
| Csf2 | 3.37E-10 | 0.6621 | 0.182 | 0.106 | 6.21E-06 |

**Supplementary Table 4.** Marker genes for CD4+ T cell clusters (Cd4Cre, Bcl6-/- and Prdm1-/- mice)

| p_val | avg_logFC | pct.1 | pct.2 | p_val_adj | cluster | gene |
|---|---|---|---|---|---|---|
| 0 | 1.000675 | 0.735 | 0.209 | 0 | 0 | Foxp3 |
| 0 | 0.940465 | 0.644 | 0.266 | 0 | 0 | Izumo1r |
| 0 | 0.816254 | 0.618 | 0.268 | 0 | 0 | Ighm |
| 0 | 0.757673 | 0.721 | 0.435 | 0 | 0 | Ass1 |
| 0 | 0.685934 | 0.872 | 0.366 | 0 | 0 | Ikzf2 |
| 0 | 0.618467 | 0.877 | 0.642 | 0 | 0 | Pim1 |
| 0 | 0.613446 | 0.496 | 0.186 | 0 | 0 | Sell |
| 1.11E-288 | 0.685301 | 0.877 | 0.719 | 1.76E-284 | 0 | H2afz |
| 7.43E-283 | 0.64352 | 0.728 | 0.485 | 1.18E-278 | 0 | Cyb5a |
| 5.06E-254 | 0.581607 | 0.936 | 0.823 | 8.06E-250 | 0 | Ltb |
| 2.69E-240 | 0.616886 | 0.447 | 0.181 | 4.28E-236 | 0 | Klrg1 |
| 1.75E-234 | 0.71688 | 0.503 | 0.233 | 2.79E-230 | 0 | Ecm1 |
| 4.48E-230 | 0.562598 | 0.737 | 0.505 | 7.14E-226 | 0 | Pglyrp1 |
| 1.29E-219 | 0.791286 | 0.378 | 0.148 | 2.06E-215 | 0 | Cd74 |
| 6.87E-147 | 0.700984 | 0.616 | 0.422 | 1.09E-142 | 0 | Zfp36 |
| 0 | 0.758286 | 0.546 | 0.212 | 0 | 1 | Tcf7 |
| 0 | 0.557006 | 0.996 | 0.943 | 0 | 1 | Rps27 |
| 0 | 0.474185 | 0.975 | 0.897 | 0 | 1 | Rps29 |
| 0 | 0.467271 | 0.999 | 0.967 | 0 | 1 | Rps24 |
| 0 | 0.462494 | 0.999 | 0.968 | 0 | 1 | Rps16 |
| 2.43E-295 | 0.510358 | 0.963 | 0.86 | 3.87E-291 | 1 | Rpl12 |
| 1.00E-268 | 0.690966 | 0.463 | 0.181 | 1.60E-264 | 1 | Itga4 |
| 5.21E-217 | 0.616072 | 0.546 | 0.28 | 8.30E-213 | 1 | S1pr1 |
| 2.50E-173 | 1.466601 | 0.505 | 0.295 | 3.98E-169 | 1 | Ccl5 |
| 2.66E-171 | 0.525097 | 0.376 | 0.162 | 4.23E-167 | 1 | Klf3 |
| 9.26E-133 | 0.498349 | 0.438 | 0.235 | 1.47E-128 | 1 | Gm2682 |
| 3.77E-114 | 0.451681 | 0.636 | 0.468 | 6.01E-110 | 1 | Cxcr3 |
| 1.28E-103 | 0.498451 | 0.563 | 0.373 | 2.03E-99 | 1 | Klf2 |
| 7.18E-38 | 0.503025 | 0.326 | 0.221 | 1.14E-33 | 1 | Hspa1a |
| 1.65E-31 | 0.454661 | 0.301 | 0.21 | 2.63E-27 | 1 | Hspa1b |
| 0 | 1.428689 | 0.308 | 0.057 | 0 | 2 | Csf2 |
| 0 | 0.832608 | 0.534 | 0.138 | 0 | 2 | Tnfsf8 |
| 0 | 0.829231 | 0.308 | 0.031 | 0 | 2 | Igfbp7 |
| 0 | 0.777491 | 0.888 | 0.481 | 0 | 2 | Bhlhe40 |
| 0 | 0.702628 | 0.943 | 0.663 | 0 | 2 | Ostf1 |
| 0 | 0.684007 | 0.581 | 0.176 | 0 | 2 | Tnfsf11 |
| 3.17E-299 | 0.861596 | 0.36 | 0.086 | 5.04E-295 | 2 | Il1r2 |
| 6.14E-296 | 0.66319 | 0.881 | 0.493 | 9.78E-292 | 2 | Cxcr6 |
| 4.21E-267 | 0.731185 | 0.726 | 0.344 | 6.70E-263 | 2 | Pdcd1 |
| 4.54E-242 | 0.833677 | 0.364 | 0.103 | 7.22E-238 | 2 | Rgs16 |
| 1.39E-225 | 0.9141 | 0.408 | 0.134 | 2.21E-221 | 2 | Lgals7 |
| 1.53E-221 | 0.89849 | 0.62 | 0.303 | 2.44E-217 | 2 | Plac8 |

| | | | | | | |
|---|---|---|---|---|---|---|
| 9.19E-220 | 0.669903 | 0.415 | 0.139 | 1.46E-215 | 2 | Klrc1 |
| 1.49E-151 | 0.875992 | 0.474 | 0.219 | 2.37E-147 | 2 | Ifitm1 |
| 1.58E-116 | 0.955326 | 0.343 | 0.156 | 2.52E-112 | 2 | Trbv3 |
| 0 | 1.255273 | 0.85 | 0.55 | 0 | 3 | Gzmb |
| 0 | 1.117794 | 0.674 | 0.177 | 0 | 3 | Klrg1 |
| 0 | 1.117099 | 0.592 | 0.21 | 0 | 3 | Lag3 |
| 0 | 1.083209 | 0.724 | 0.295 | 0 | 3 | Cst7 |
| 0 | 1.030299 | 0.986 | 0.732 | 0 | 3 | Tnfrsf4 |
| 0 | 1.014396 | 0.885 | 0.373 | 0 | 3 | Il2ra |
| 0 | 1.014016 | 0.852 | 0.437 | 0 | 3 | Tigit |
| 0 | 1.005903 | 0.858 | 0.54 | 0 | 3 | Ddit4 |
| 0 | 0.92319 | 0.968 | 0.589 | 0 | 3 | Tnfrsf9 |
| 0 | 0.914941 | 0.834 | 0.459 | 0 | 3 | Gimap7 |
| 0 | 0.880204 | 0.985 | 0.896 | 0 | 3 | Gapdh |
| 0 | 0.86589 | 0.792 | 0.326 | 0 | 3 | Arl5a |
| 0 | 0.858402 | 0.89 | 0.605 | 0 | 3 | Zfp36l1 |
| 5.67E-204 | 0.93884 | 0.303 | 0.083 | 9.03E-200 | 3 | Rln3 |
| 4.13E-145 | 0.910056 | 0.395 | 0.165 | 6.58E-141 | 3 | Cd74 |
| 0 | 1.429864 | 0.609 | 0.207 | 0 | 4 | Ifitm1 |
| 0 | 1.063705 | 0.81 | 0.374 | 0 | 4 | Ifitm2 |
| 0 | 0.680672 | 0.914 | 0.494 | 0 | 4 | Cxcr6 |
| 0 | 0.568906 | 0.997 | 0.951 | 0 | 4 | Cd52 |
| 0 | 0.472144 | 0.999 | 0.936 | 0 | 4 | Rpl29 |
| 7.41E-284 | 0.694447 | 0.809 | 0.405 | 1.18E-279 | 4 | Ctsw |
| 1.56E-280 | 1.096092 | 0.549 | 0.209 | 2.49E-276 | 4 | Cd7 |
| 4.31E-244 | 0.597835 | 0.923 | 0.634 | 6.86E-240 | 4 | Id2 |
| 9.74E-216 | 0.592729 | 0.434 | 0.149 | 1.55E-211 | 4 | Anxa1 |
| 1.16E-197 | 0.561732 | 0.886 | 0.631 | 1.85E-193 | 4 | Itgb1 |
| 3.81E-171 | 0.625918 | 0.731 | 0.432 | 6.06E-167 | 4 | Ifitm3 |
| 2.82E-170 | 0.470745 | 0.855 | 0.546 | 4.50E-166 | 4 | Nkg7 |
| 1.77E-161 | 0.506189 | 0.771 | 0.487 | 2.82E-157 | 4 | Zyx |
| 1.30E-129 | 0.515988 | 0.454 | 0.206 | 2.07E-125 | 4 | Klrd1 |
| 6.14E-123 | 0.693684 | 0.641 | 0.41 | 9.78E-119 | 4 | Lgals3 |
| 0 | 2.090111 | 0.989 | 0.451 | 0 | 5 | Isg15 |
| 0 | 2.060296 | 0.885 | 0.176 | 0 | 5 | Ifit3 |
| 0 | 1.854165 | 0.896 | 0.198 | 0 | 5 | Ifit1 |
| 0 | 1.620959 | 0.69 | 0.088 | 0 | 5 | Ifit3b |
| 0 | 1.512321 | 0.694 | 0.08 | 0 | 5 | Rsad2 |
| 0 | 1.457886 | 0.834 | 0.186 | 0 | 5 | Usp18 |
| 0 | 1.444273 | 0.937 | 0.453 | 0 | 5 | Isg20 |
| 0 | 1.422354 | 0.67 | 0.086 | 0 | 5 | Mx1 |
| 0 | 1.301608 | 0.872 | 0.387 | 0 | 5 | Phf11b |
| 0 | 1.287664 | 0.957 | 0.597 | 0 | 5 | Bst2 |
| 0 | 1.265772 | 0.925 | 0.419 | 0 | 5 | Irf7 |
| 0 | 1.23747 | 0.735 | 0.216 | 0 | 5 | Slfn5 |
| 0 | 1.220269 | 0.831 | 0.267 | 0 | 5 | Ifi208 |

| | | | | | | |
|---|---|---|---|---|---|---|
| 0 | 1.137689 | 0.764 | 0.252 | 0 | 5 | Daxx |
| 3.05E-122 | 1.185561 | 0.475 | 0.25 | 4.86E-118 | 5 | Ly6c2 |
| 0 | 0.966273 | 0.92 | 0.562 | 0 | 6 | Tpi1 |
| 0 | 0.94101 | 0.993 | 0.882 | 0 | 6 | Aldoa |
| 0 | 0.905663 | 0.91 | 0.61 | 0 | 6 | Pgk1 |
| 8.17E-280 | 0.774544 | 0.93 | 0.675 | 1.30E-275 | 6 | Mif |
| 2.52E-270 | 0.679429 | 0.973 | 0.807 | 4.01E-266 | 6 | Eno1 |
| 1.50E-255 | 0.8407 | 0.713 | 0.328 | 2.39E-251 | 6 | Ctla2a |
| 3.76E-192 | 1.064297 | 0.709 | 0.39 | 5.98E-188 | 6 | Ifitm2 |
| 9.33E-191 | 0.685187 | 0.891 | 0.642 | 1.49E-186 | 6 | Id2 |
| 7.53E-162 | 0.903376 | 0.614 | 0.312 | 1.20E-157 | 6 | Plac8 |
| 2.84E-160 | 0.830387 | 0.64 | 0.326 | 4.52E-156 | 6 | Hilpda |
| 6.43E-156 | 0.768619 | 0.735 | 0.437 | 1.02E-151 | 6 | Ifitm3 |
| 6.87E-108 | 0.733563 | 0.658 | 0.413 | 1.09E-103 | 6 | Lgals3 |
| 2.10E-104 | 0.680302 | 0.378 | 0.165 | 3.34E-100 | 6 | Lta |
| 6.59E-83 | 0.822945 | 0.464 | 0.255 | 1.05E-78 | 6 | Ly6c2 |
| 5.27E-65 | 0.864173 | 0.414 | 0.231 | 8.40E-61 | 6 | Ifitm1 |
| 0 | 1.130375 | 0.94 | 0.873 | 0 | 7 | Mbnl1 |
| 6.83E-208 | 1.12811 | 0.704 | 0.47 | 1.09E-203 | 7 | Lnpep |
| 3.07E-183 | 1.411057 | 0.727 | 0.513 | 4.89E-179 | 7 | Gm26917 |
| 3.74E-156 | 1.022369 | 0.698 | 0.531 | 5.96E-152 | 7 | Mycbp2 |
| 8.32E-138 | 1.035298 | 0.633 | 0.43 | 1.32E-133 | 7 | Nr3c1 |
| 3.73E-129 | 0.977062 | 0.429 | 0.195 | 5.95E-125 | 7 | Arhgap26 |
| 5.20E-128 | 1.004808 | 0.556 | 0.347 | 8.28E-124 | 7 | 4932438A13Rik |
| 5.98E-125 | 0.977378 | 0.579 | 0.366 | 9.52E-121 | 7 | Fryl |
| 7.12E-125 | 0.976285 | 0.629 | 0.449 | 1.13E-120 | 7 | Utrn |
| 1.79E-120 | 1.054638 | 0.536 | 0.327 | 2.86E-116 | 7 | Themis |
| 2.33E-102 | 1.019585 | 0.693 | 0.605 | 3.71E-98 | 7 | Inpp4b |
| 2.72E-92 | 1.072168 | 0.372 | 0.18 | 4.34E-88 | 7 | Lyst |
| 5.27E-68 | 1.029433 | 0.493 | 0.349 | 8.40E-64 | 7 | Rnf213 |
| 3.61E-53 | 1.019709 | 0.382 | 0.242 | 5.75E-49 | 7 | Bcl2 |
| 1.40E-29 | 0.976831 | 0.349 | 0.259 | 2.23E-25 | 7 | Slfn5 |
| 0 | 1.687096 | 0.891 | 0.425 | 0 | 8 | Ikzf2 |
| 0 | 1.092708 | 1 | 0.997 | 0 | 8 | Malat1 |
| 3.04E-284 | 1.428923 | 0.776 | 0.401 | 4.85E-280 | 8 | Il2ra |
| 1.43E-247 | 1.384275 | 0.695 | 0.352 | 2.28E-243 | 8 | Arl5a |
| 4.87E-235 | 1.239878 | 0.8 | 0.607 | 7.76E-231 | 8 | Wnk1 |
| 1.48E-209 | 1.108405 | 0.89 | 0.82 | 2.36E-205 | 8 | Icos |
| 6.11E-198 | 1.256178 | 0.616 | 0.309 | 9.74E-194 | 8 | Itgav |
| 2.79E-182 | 1.262782 | 0.454 | 0.165 | 4.44E-178 | 8 | Neb |
| 2.65E-143 | 1.168565 | 0.425 | 0.167 | 4.23E-139 | 8 | Itgae |
| 1.93E-136 | 1.091459 | 0.589 | 0.373 | 3.08E-132 | 8 | Plec |
| 5.39E-126 | 1.177279 | 0.744 | 0.575 | 8.59E-122 | 8 | Ctla4 |
| 7.33E-120 | 1.124721 | 0.578 | 0.369 | 1.17E-115 | 8 | Malt1 |
| 1.77E-118 | 1.11577 | 0.483 | 0.251 | 2.81E-114 | 8 | Lrba |
| 8.62E-107 | 1.12666 | 0.471 | 0.25 | 1.37E-102 | 8 | Ccr5 |

| | | | | | | |
|---|---|---|---|---|---|---|
| 1.82E-75 | 1.124683 | 0.609 | 0.504 | 2.89E-71 | 8 | Rabgap1l |
| 1.99E-281 | 1.321085 | 0.792 | 0.249 | 3.17E-277 | 9 | Nr4a1 |
| 1.94E-228 | 1.161367 | 0.948 | 0.597 | 3.09E-224 | 9 | Btg2 |
| 2.01E-221 | 0.989855 | 0.996 | 0.833 | 3.20E-217 | 9 | Junb |
| 2.28E-206 | 1.012406 | 0.889 | 0.46 | 3.62E-202 | 9 | Ier2 |
| 1.70E-182 | 0.97342 | 0.955 | 0.682 | 2.70E-178 | 9 | Nfkbia |
| 8.95E-182 | 0.898824 | 0.666 | 0.223 | 1.43E-177 | 9 | Nfkbid |
| 2.09E-181 | 0.904108 | 0.921 | 0.497 | 3.32E-177 | 9 | Tnfaip3 |
| 9.99E-144 | 1.05061 | 0.748 | 0.355 | 1.59E-139 | 9 | Gadd45b |
| 1.46E-142 | 0.889126 | 0.881 | 0.552 | 2.32E-138 | 9 | Dusp5 |
| 4.56E-139 | 0.887569 | 0.79 | 0.409 | 7.27E-135 | 9 | Nfkbiz |
| 1.87E-131 | 0.843786 | 0.45 | 0.137 | 2.98E-127 | 9 | Dusp10 |
| 5.52E-120 | 0.797934 | 0.337 | 0.084 | 8.79E-116 | 9 | Fosb |
| 4.25E-119 | 0.869393 | 0.822 | 0.482 | 6.77E-115 | 9 | Vps37b |
| 4.04E-67 | 0.813401 | 0.44 | 0.194 | 6.43E-63 | 9 | Rgcc |
| 3.49E-40 | 0.806623 | 0.511 | 0.288 | 5.56E-36 | 9 | Fos |
| 9.70E-247 | 1.112477 | 0.541 | 0.118 | 1.55E-242 | 10 | Nr4a3 |
| 1.02E-241 | 0.950647 | 0.644 | 0.179 | 1.63E-237 | 10 | C1qbp |
| 2.71E-215 | 0.92935 | 0.394 | 0.07 | 4.31E-211 | 10 | Myc |
| 2.79E-196 | 3.119191 | 0.309 | 0.049 | 4.44E-192 | 10 | Il2 |
| 8.64E-183 | 1.382502 | 0.642 | 0.224 | 1.38E-178 | 10 | Nfkbid |
| 2.56E-163 | 0.972432 | 0.875 | 0.552 | 4.07E-159 | 10 | Eif4a1 |
| 9.81E-146 | 1.26618 | 0.65 | 0.254 | 1.56E-141 | 10 | Nr4a1 |
| 6.81E-138 | 2.083034 | 0.329 | 0.074 | 1.08E-133 | 10 | Csf2 |
| 1.95E-134 | 2.600701 | 0.487 | 0.164 | 3.11E-130 | 10 | Ifng |
| 4.62E-132 | 0.942132 | 0.756 | 0.355 | 7.36E-128 | 10 | Gadd45b |
| 2.06E-119 | 1.075807 | 0.506 | 0.18 | 3.29E-115 | 10 | Irf8 |
| 1.36E-91 | 0.938592 | 0.51 | 0.208 | 2.16E-87 | 10 | Tnfsf11 |
| 9.49E-66 | 1.370879 | 0.398 | 0.163 | 1.51E-61 | 10 | Tnf |
| 1.95E-65 | 1.065669 | 0.608 | 0.357 | 3.10E-61 | 10 | Cd40lg |
| 9.10E-41 | 2.2857 | 0.303 | 0.138 | 1.45E-36 | 10 | Ccl4 |
| 0 | 1.89083 | 0.762 | 0.032 | 0 | 11 | Tmem176b |
| 0 | 1.776198 | 0.705 | 0.025 | 0 | 11 | Tmem176a |
| 5.28E-210 | 0.973387 | 0.443 | 0.034 | 8.40E-206 | 11 | Aqp3 |
| 8.00E-117 | 0.877635 | 0.362 | 0.04 | 1.27E-112 | 11 | Ccr6 |
| 3.40E-61 | 0.705337 | 0.371 | 0.074 | 5.41E-57 | 11 | Nebl |
| 6.80E-54 | 0.92527 | 0.586 | 0.198 | 1.08E-49 | 11 | Ccr4 |
| 4.35E-44 | 0.751877 | 0.443 | 0.127 | 6.94E-40 | 11 | Serpinb1a |
| 2.50E-37 | 0.595577 | 0.352 | 0.097 | 3.99E-33 | 11 | Ramp1 |
| 3.40E-34 | 0.63563 | 0.305 | 0.081 | 5.41E-30 | 11 | 1110034G24Rik |
| 5.56E-32 | 0.777526 | 0.876 | 0.582 | 8.85E-28 | 11 | Il7r |
| 3.60E-30 | 0.815961 | 0.752 | 0.518 | 5.73E-26 | 11 | Gpx1 |
| 2.52E-29 | 0.600708 | 0.876 | 0.641 | 4.01E-25 | 11 | Emb |
| 7.25E-24 | 0.618201 | 0.357 | 0.131 | 1.15E-19 | 11 | Ltb4r1 |
| 1.87E-23 | 0.653192 | 0.4 | 0.166 | 2.98E-19 | 11 | Acsbg1 |
| 3.52E-17 | 0.655853 | 0.638 | 0.398 | 5.60E-13 | 11 | Klf2 |

| | | | | | | |
|---|---|---|---|---|---|---|
| 0 | 1.052668 | 0.402 | 0.008 | 0 | 12 | Trgv2 |
| 2.14E-131 | 1.05238 | 0.364 | 0.024 | 3.41E-127 | 12 | Fcer1g |
| 1.38E-90 | 0.736987 | 0.356 | 0.033 | 2.19E-86 | 12 | Xcl1 |
| 5.23E-65 | 1.6453 | 0.826 | 0.268 | 8.33E-61 | 12 | Ly6c2 |
| 1.61E-42 | 0.607916 | 1 | 0.976 | 2.56E-38 | 12 | Rpl13 |
| 1.29E-39 | 0.650288 | 1 | 0.952 | 2.05E-35 | 12 | Rps20 |
| 1.07E-37 | 1.18618 | 0.758 | 0.322 | 1.71E-33 | 12 | Ccl5 |
| 1.18E-37 | 0.905824 | 0.712 | 0.247 | 1.87E-33 | 12 | Bcl2 |
| 2.83E-34 | 0.752815 | 0.902 | 0.441 | 4.51E-30 | 12 | Ctsw |
| 5.22E-34 | 0.713377 | 0.992 | 0.813 | 8.31E-30 | 12 | Rpl36a |
| 1.13E-32 | 0.615757 | 1 | 0.916 | 1.79E-28 | 12 | Rps18 |
| 2.55E-29 | 0.603432 | 1 | 0.874 | 4.05E-25 | 12 | Rpl12 |
| 1.82E-28 | 0.643194 | 0.5 | 0.153 | 2.90E-24 | 12 | Pde3b |
| 2.38E-15 | 0.590078 | 0.53 | 0.239 | 3.79E-11 | 12 | Cd7 |
| 5.02E-13 | 0.719722 | 0.424 | 0.191 | 7.99E-09 | 12 | Lyst |

**Supplementary Table 5.** Genes differentially expressed between *Cd4* Cre and *Bcl6* -/- TEFF or Cd4Cre and Prdm1-/- TEFF

| GeneID | p_val | avg_logFC | pct.1 | pct.2 | p_val_adj | Comparison |
|---|---|---|---|---|---|---|
| Gzmb | 5.9856052993667e-320 | -1.797256 | 0.076 | 0.497 | 9.53267499 | Cre_Blimp |
| Gm26917 | 1.43E-134 | -0.922099 | 0.403 | 0.672 | 2.29E-130 | Cre_Blimp |
| Hspa1a | 7.02E-81 | -0.891346 | 0.18 | 0.388 | 1.12E-76 | Cre_Blimp |
| AY036118 | 4.84E-155 | -0.844114 | 0.571 | 0.828 | 7.71E-151 | Cre_Blimp |
| Hspa1b | 6.83E-78 | -0.830448 | 0.15 | 0.35 | 1.09E-73 | Cre_Blimp |
| Ccl4 | 6.23E-104 | -0.784767 | 0.03 | 0.196 | 9.91E-100 | Cre_Blimp |
| Ifi27l2a | 2.28E-170 | -0.752804 | 0.774 | 0.951 | 3.62E-166 | Cre_Blimp |
| Jund | 6.41E-255 | -0.735796 | 0.732 | 0.937 | 1.02E-250 | Cre_Blimp |
| Malat1 | 7.10E-277 | -0.714283 | 0.992 | 1 | 1.13E-272 | Cre_Blimp |
| Ifi209 | 1.40E-158 | -0.694156 | 0.23 | 0.555 | 2.23E-154 | Cre_Blimp |
| Isg15 | 6.00E-129 | -0.692184 | 0.295 | 0.597 | 9.55E-125 | Cre_Blimp |
| Lars2 | 1.00E-106 | -0.673272 | 0.657 | 0.859 | 1.60E-102 | Cre_Blimp |
| Gm42418 | 8.04E-236 | -0.662107 | 0.999 | 1 | 1.28E-231 | Cre_Blimp |
| Ifit3 | 7.96E-88 | -0.651168 | 0.152 | 0.372 | 1.27E-83 | Cre_Blimp |
| Mxd1 | 2.99E-84 | -0.648593 | 0.224 | 0.446 | 4.76E-80 | Cre_Blimp |
| Rnf213 | 4.65E-124 | -0.632918 | 0.188 | 0.469 | 7.40E-120 | Cre_Blimp |
| Tnfrsf9 | 9.81E-76 | -0.631649 | 0.337 | 0.564 | 1.56E-71 | Cre_Blimp |
| Stat1 | 5.51E-120 | -0.629605 | 0.573 | 0.81 | 8.77E-116 | Cre_Blimp |
| Ecm1 | 3.16E-52 | -0.611233 | 0.065 | 0.192 | 5.04E-48 | Cre_Blimp |
| Gbp7 | 1.80E-134 | -0.606935 | 0.24 | 0.54 | 2.87E-130 | Cre_Blimp |
| Ifit2 | 1.76E-60 | -0.596678 | 0.072 | 0.215 | 2.81E-56 | Cre_Blimp |
| Ifi208 | 6.89E-111 | -0.585966 | 0.214 | 0.481 | 1.10E-106 | Cre_Blimp |
| Rsad2 | 8.55E-52 | -0.564982 | 0.071 | 0.202 | 1.36E-47 | Cre_Blimp |
| Slfn5 | 4.27E-57 | -0.564387 | 0.191 | 0.37 | 6.80E-53 | Cre_Blimp |
| Mx1 | 1.19E-52 | -0.563771 | 0.09 | 0.231 | 1.90E-48 | Cre_Blimp |
| Jun | 2.87E-44 | -0.551476 | 0.172 | 0.324 | 4.58E-40 | Cre_Blimp |
| Ankrd11 | 3.97E-175 | -0.537331 | 0.444 | 0.781 | 6.32E-171 | Cre_Blimp |
| Smchd1 | 2.23E-93 | -0.531456 | 0.312 | 0.564 | 3.55E-89 | Cre_Blimp |
| Ifit3b | 6.12E-61 | -0.525911 | 0.083 | 0.234 | 9.75E-57 | Cre_Blimp |
| Gbp9 | 5.49E-111 | -0.525217 | 0.139 | 0.386 | 8.75E-107 | Cre_Blimp |
| Ifit1 | 8.09E-86 | -0.521954 | 0.152 | 0.367 | 1.29E-81 | Cre_Blimp |
| Rgs1 | 2.14E-59 | -0.518558 | 0.456 | 0.649 | 3.41E-55 | Cre_Blimp |
| Adam19 | 5.83E-74 | -0.515657 | 0.198 | 0.402 | 9.28E-70 | Cre_Blimp |
| Pdcd1 | 2.17E-48 | -0.50609 | 0.368 | 0.531 | 3.45E-44 | Cre_Blimp |
| Daxx | 2.22E-75 | -0.500523 | 0.198 | 0.411 | 3.54E-71 | Cre_Blimp |
| Ccl5 | 1.30E-51 | -0.500346 | 0.239 | 0.412 | 2.06E-47 | Cre_Blimp |
| Tmem64 | 1.78E-71 | -0.490429 | 0.135 | 0.324 | 2.83E-67 | Cre_Blimp |
| Rgs16 | 5.89E-54 | -0.485124 | 0.062 | 0.188 | 9.38E-50 | Cre_Blimp |
| Ifi214 | 1.65E-88 | -0.484662 | 0.073 | 0.258 | 2.63E-84 | Cre_Blimp |
| Ccr5 | 4.89E-72 | -0.482929 | 0.054 | 0.201 | 7.79E-68 | Cre_Blimp |
| Stat3 | 5.14E-96 | -0.477687 | 0.454 | 0.702 | 8.19E-92 | Cre_Blimp |

| | | | | | | |
|---|---|---|---|---|---|---|
| Parp14 | 7.39E-68 | -0.477274 | 0.28 | 0.491 | 1.18E-63 | Cre_Blimp |
| Lag3 | 1.24E-31 | -0.475335 | 0.131 | 0.229 | 1.98E-27 | Cre_Blimp |
| Slfn8 | 3.28E-83 | -0.46955 | 0.13 | 0.334 | 5.22E-79 | Cre_Blimp |
| Arl5a | 8.18E-89 | -0.468811 | 0.08 | 0.268 | 1.30E-84 | Cre_Blimp |
| Bst2 | 8.39E-74 | -0.468636 | 0.469 | 0.685 | 1.34E-69 | Cre_Blimp |
| Trim30a | 9.65E-87 | -0.46794 | 0.272 | 0.516 | 1.54E-82 | Cre_Blimp |
| Isg20 | 1.25E-63 | -0.466713 | 0.324 | 0.53 | 2.00E-59 | Cre_Blimp |
| Herc6 | 2.13E-73 | -0.465131 | 0.166 | 0.368 | 3.39E-69 | Cre_Blimp |
| Dnajb1 | 4.20E-51 | -0.465085 | 0.278 | 0.457 | 6.69E-47 | Cre_Blimp |
| CAAA01147332.1 | 3.05E-77 | -0.46262 | 0.247 | 0.424 | 4.85E-73 | Cre_Blimp |
| Wnk1 | 5.82E-83 | -0.457417 | 0.378 | 0.615 | 9.28E-79 | Cre_Blimp |
| Hsph1 | 1.13E-61 | -0.457084 | 0.149 | 0.326 | 1.80E-57 | Cre_Blimp |
| Fam102a | 2.74E-82 | -0.450589 | 0.353 | 0.591 | 4.37E-78 | Cre_Blimp |
| Dusp2 | 1.34E-40 | -0.449175 | 0.344 | 0.506 | 2.13E-36 | Cre_Blimp |
| Ccr8 | 6.50E-38 | -0.447957 | 0.165 | 0.291 | 1.04E-33 | Cre_Blimp |
| Usp25 | 5.83E-78 | -0.446108 | 0.26 | 0.487 | 9.29E-74 | Cre_Blimp |
| Irf7 | 2.95E-77 | -0.440795 | 0.326 | 0.561 | 4.70E-73 | Cre_Blimp |
| Itgb3 | 9.07E-66 | -0.433128 | 0.167 | 0.358 | 1.44E-61 | Cre_Blimp |
| Maf | 2.66E-48 | -0.431887 | 0.279 | 0.448 | 4.23E-44 | Cre_Blimp |
| Samhd1 | 1.49E-56 | -0.430272 | 0.527 | 0.707 | 2.37E-52 | Cre_Blimp |
| Slfn1 | 5.61E-76 | -0.430035 | 0.5 | 0.713 | 8.93E-72 | Cre_Blimp |
| Stat2 | 3.62E-70 | -0.425822 | 0.128 | 0.313 | 5.76E-66 | Cre_Blimp |
| Tnks2 | 3.73E-99 | -0.424679 | 0.251 | 0.511 | 5.94E-95 | Cre_Blimp |
| Ifi203 | 5.19E-75 | -0.422614 | 0.447 | 0.671 | 8.26E-71 | Cre_Blimp |
| Malt1 | 4.17E-61 | -0.421171 | 0.195 | 0.385 | 6.64E-57 | Cre_Blimp |
| Zbp1 | 4.62E-83 | -0.420765 | 0.397 | 0.638 | 7.36E-79 | Cre_Blimp |
| Rtp4 | 2.61E-104 | -0.419605 | 0.261 | 0.527 | 4.16E-100 | Cre_Blimp |
| Baz1a | 1.23E-69 | -0.413299 | 0.275 | 0.491 | 1.96E-65 | Cre_Blimp |
| Ifih1 | 1.34E-60 | -0.412867 | 0.136 | 0.309 | 2.14E-56 | Cre_Blimp |
| Ccr2 | 1.57E-40 | -0.412813 | 0.218 | 0.371 | 2.50E-36 | Cre_Blimp |
| Atp2b4 | 2.60E-73 | -0.411717 | 0.145 | 0.341 | 4.14E-69 | Cre_Blimp |
| Peli1 | 5.05E-80 | -0.411408 | 0.501 | 0.726 | 8.04E-76 | Cre_Blimp |
| Myh9 | 2.96E-101 | -0.41126 | 0.665 | 0.872 | 4.71E-97 | Cre_Blimp |
| Mycbp2 | 6.67E-91 | -0.409562 | 0.418 | 0.671 | 1.06E-86 | Cre_Blimp |
| Oasl2 | 1.10E-60 | -0.408591 | 0.092 | 0.247 | 1.75E-56 | Cre_Blimp |
| Ccnd2 | 5.63E-73 | -0.407326 | 0.471 | 0.689 | 8.97E-69 | Cre_Blimp |
| Gzmk | 4.09E-30 | -0.404576 | 0.044 | 0.106 | 6.51E-26 | Cre_Bcl6 |
| Gbp4 | 1.85E-64 | -0.403922 | 0.272 | 0.48 | 2.95E-60 | Cre_Blimp |
| Dtx3l | 9.15E-75 | -0.403822 | 0.2 | 0.413 | 1.46E-70 | Cre_Blimp |
| Zfp292 | 7.62E-71 | -0.403588 | 0.277 | 0.496 | 1.21E-66 | Cre_Blimp |
| Gimap3 | 2.47E-81 | -0.402684 | 0.709 | 0.874 | 3.94E-77 | Cre_Blimp |
| Odc1 | 3.60E-47 | -0.400596 | 0.507 | 0.632 | 5.73E-43 | Cre_Bcl6 |
| Cab39 | 5.34E-98 | -0.398147 | 0.234 | 0.489 | 8.51E-94 | Cre_Blimp |
| Mndal | 1.85E-68 | -0.397277 | 0.477 | 0.687 | 2.95E-64 | Cre_Blimp |
| Camk1d | 3.60E-38 | -0.393618 | 0.235 | 0.383 | 5.74E-34 | Cre_Blimp |

| | | | | | | |
|---|---|---|---|---|---|---|
| Akap13 | 4.11E-72 | -0.389296 | 0.662 | 0.838 | 6.54E-68 | Cre_Blimp |
| Cxcr6 | 3.27E-42 | -0.389183 | 0.583 | 0.733 | 5.20E-38 | Cre_Blimp |
| Pkn1 | 2.91E-88 | -0.387718 | 0.242 | 0.485 | 4.64E-84 | Cre_Blimp |
| Samd9l | 5.10E-57 | -0.384518 | 0.138 | 0.305 | 8.13E-53 | Cre_Blimp |
| Helz2 | 6.91E-65 | -0.384407 | 0.142 | 0.323 | 1.10E-60 | Cre_Blimp |
| 2010111I01Rik | 8.80E-74 | -0.384148 | 0.317 | 0.544 | 1.40E-69 | Cre_Blimp |
| Nsd3 | 4.07E-93 | -0.382799 | 0.479 | 0.728 | 6.47E-89 | Cre_Blimp |
| Ddx58 | 1.14E-72 | -0.381046 | 0.184 | 0.391 | 1.82E-68 | Cre_Blimp |
| Hif1a | 4.91E-56 | -0.380362 | 0.725 | 0.87 | 7.83E-52 | Cre_Blimp |
| Slamf6 | 6.23E-68 | -0.37921 | 0.196 | 0.358 | 9.92E-64 | Cre_Bcl6 |
| 9930111J21Rik2 | 5.86E-59 | -0.378178 | 0.198 | 0.383 | 9.33E-55 | Cre_Blimp |
| Xaf1 | 8.80E-77 | -0.376504 | 0.238 | 0.463 | 1.40E-72 | Cre_Blimp |
| Ifi213 | 1.66E-70 | -0.37562 | 0.221 | 0.433 | 2.64E-66 | Cre_Blimp |
| Son | 8.45E-98 | -0.375567 | 0.533 | 0.778 | 1.35E-93 | Cre_Blimp |
| Ikzf2 | 3.53E-66 | -0.374104 | 0.138 | 0.293 | 5.62E-62 | Cre_Bcl6 |
| Nr3c1 | 9.18E-78 | -0.372842 | 0.29 | 0.523 | 1.46E-73 | Cre_Blimp |
| Chd3 | 6.31E-77 | -0.369321 | 0.271 | 0.5 | 1.00E-72 | Cre_Blimp |
| Hspa5 | 2.25E-51 | -0.36809 | 0.598 | 0.766 | 3.59E-47 | Cre_Blimp |
| Ppp6r1 | 5.52E-77 | -0.367476 | 0.138 | 0.338 | 8.79E-73 | Cre_Blimp |
| Chmp4b | 3.67E-94 | -0.367304 | 0.589 | 0.806 | 5.85E-90 | Cre_Blimp |
| Prrc2c | 3.93E-67 | -0.364072 | 0.561 | 0.76 | 6.26E-63 | Cre_Blimp |
| Galnt6 | 6.54E-58 | -0.362493 | 0.167 | 0.344 | 1.04E-53 | Cre_Blimp |
| Tln1 | 1.61E-76 | -0.362006 | 0.499 | 0.723 | 2.56E-72 | Cre_Blimp |
| Prkch | 4.79E-99 | -0.36135 | 0.303 | 0.569 | 7.63E-95 | Cre_Blimp |
| Dgat1 | 8.59E-42 | -0.360945 | 0.38 | 0.551 | 1.37E-37 | Cre_Blimp |
| Mknk2 | 2.15E-91 | -0.360463 | 0.193 | 0.43 | 3.42E-87 | Cre_Blimp |
| Etnk1 | 2.42E-47 | -0.356703 | 0.167 | 0.325 | 3.85E-43 | Cre_Blimp |
| Slamf1 | 1.72E-58 | -0.356652 | 0.21 | 0.399 | 2.74E-54 | Cre_Blimp |
| Srrm2 | 8.27E-99 | -0.356498 | 0.663 | 0.873 | 1.32E-94 | Cre_Blimp |
| Ppp2ca | 2.89E-97 | -0.356238 | 0.401 | 0.612 | 4.60E-93 | Cre_Bcl6 |
| Il2rb | 1.07E-70 | -0.355791 | 0.669 | 0.847 | 1.71E-66 | Cre_Blimp |
| Parp9 | 8.99E-59 | -0.355569 | 0.202 | 0.387 | 1.43E-54 | Cre_Blimp |
| Klf6 | 1.88E-53 | -0.355 | 0.435 | 0.63 | 3.00E-49 | Cre_Blimp |
| Ncor1 | 2.60E-105 | -0.352758 | 0.526 | 0.783 | 4.14E-101 | Cre_Blimp |
| Bcl2l11 | 6.47E-56 | -0.352133 | 0.11 | 0.266 | 1.03E-51 | Cre_Blimp |
| Gnai2 | 4.64E-111 | -0.352079 | 0.724 | 0.881 | 7.39E-107 | Cre_Blimp |
| Klf13 | 3.81E-90 | -0.351849 | 0.299 | 0.551 | 6.07E-86 | Cre_Blimp |
| Neat1 | 2.22E-24 | -0.351374 | 0.222 | 0.333 | 3.53E-20 | Cre_Blimp |
| Lbh | 2.42E-57 | -0.349573 | 0.276 | 0.471 | 3.85E-53 | Cre_Blimp |
| Zc3hav1 | 2.48E-76 | -0.349229 | 0.431 | 0.663 | 3.95E-72 | Cre_Blimp |
| Cdkn1b | 2.53E-83 | -0.34743 | 0.308 | 0.551 | 4.02E-79 | Cre_Blimp |
| Ms4a4c | 2.38E-33 | -0.34722 | 0.208 | 0.346 | 3.79E-29 | Cre_Blimp |
| Rora | 7.86E-50 | -0.343691 | 0.415 | 0.552 | 1.25E-45 | Cre_Bcl6 |
| Cmpk2 | 1.63E-25 | -0.34231 | 0.096 | 0.187 | 2.59E-21 | Cre_Blimp |

| | | | | | | |
|---|---|---|---|---|---|---|
| Zdhhc18 | 1.85E-93 | -0.340678 | 0.225 | 0.47 | 2.95E-89 | Cre_Blimp |
| Iigp1 | 2.19E-26 | -0.340633 | 0.092 | 0.186 | 3.48E-22 | Cre_Blimp |
| Pmepa1 | 6.66E-62 | -0.339927 | 0.097 | 0.257 | 1.06E-57 | Cre_Blimp |
| Ccdc88c | 8.13E-59 | -0.339833 | 0.221 | 0.413 | 1.29E-54 | Cre_Blimp |
| Ascc3 | 5.35E-69 | -0.33979 | 0.169 | 0.365 | 8.52E-65 | Cre_Blimp |
| Arl4c | 8.89E-43 | -0.338884 | 0.334 | 0.506 | 1.42E-38 | Cre_Blimp |
| Gm20400 | 1.01E-29 | -0.338683 | 0.117 | 0.202 | 1.61E-25 | Cre_Blimp |
| Ifng | 1.24E-27 | -0.338478 | 0.151 | 0.265 | 1.97E-23 | Cre_Blimp |
| Neurl3 | 7.56E-66 | -0.337683 | 0.227 | 0.429 | 1.20E-61 | Cre_Blimp |
| Pim1 | 1.60E-41 | -0.336863 | 0.472 | 0.64 | 2.55E-37 | Cre_Blimp |
| Lta | 6.99E-30 | -0.336343 | 0.153 | 0.244 | 1.11E-25 | Cre_Bcl6 |
| Gtpbp2 | 4.10E-73 | -0.336279 | 0.178 | 0.385 | 6.53E-69 | Cre_Blimp |
| Grb2 | 6.69E-113 | -0.335928 | 0.323 | 0.607 | 1.07E-108 | Cre_Blimp |
| Ppp1r12a | 1.13E-92 | -0.335898 | 0.603 | 0.83 | 1.81E-88 | Cre_Blimp |
| Ubc | 2.97E-57 | -0.335762 | 0.703 | 0.85 | 4.72E-53 | Cre_Blimp |
| Cyld | 3.74E-65 | -0.335608 | 0.27 | 0.481 | 5.96E-61 | Cre_Blimp |
| Mdfic | 5.10E-47 | -0.335334 | 0.208 | 0.373 | 8.13E-43 | Cre_Blimp |
| Arih1 | 2.09E-81 | -0.334853 | 0.191 | 0.413 | 3.32E-77 | Cre_Blimp |
| Trafd1 | 6.65E-51 | -0.334537 | 0.235 | 0.414 | 1.06E-46 | Cre_Blimp |
| Cd6 | 8.60E-63 | -0.334053 | 0.413 | 0.625 | 1.37E-58 | Cre_Blimp |
| Tap1 | 4.52E-75 | -0.333646 | 0.54 | 0.757 | 7.19E-71 | Cre_Blimp |
| Nktr | 2.95E-53 | -0.333087 | 0.301 | 0.492 | 4.70E-49 | Cre_Blimp |
| Igtp | 2.48E-54 | -0.332665 | 0.402 | 0.6 | 3.95E-50 | Cre_Blimp |
| Ago2 | 7.44E-58 | -0.331593 | 0.22 | 0.409 | 1.18E-53 | Cre_Blimp |
| Cmtm6 | 3.70E-45 | -0.33047 | 0.199 | 0.359 | 5.89E-41 | Cre_Blimp |
| Ugcg | 1.79E-50 | -0.330318 | 0.138 | 0.294 | 2.85E-46 | Cre_Blimp |
| Tap2 | 3.70E-65 | -0.330018 | 0.545 | 0.746 | 5.89E-61 | Cre_Blimp |
| Lrp10 | 3.82E-75 | -0.328814 | 0.454 | 0.677 | 6.08E-71 | Cre_Blimp |
| Tnfrsf18 | 4.35E-55 | -0.328068 | 0.686 | 0.838 | 6.93E-51 | Cre_Blimp |
| Cish | 1.38E-44 | -0.327664 | 0.337 | 0.51 | 2.20E-40 | Cre_Blimp |
| Prex1 | 2.19E-64 | -0.327127 | 0.313 | 0.526 | 3.48E-60 | Cre_Blimp |
| Ptk2b | 5.70E-71 | -0.326509 | 0.237 | 0.453 | 9.07E-67 | Cre_Blimp |
| Camk2n1 | 6.12E-49 | -0.325838 | 0.09 | 0.225 | 9.75E-45 | Cre_Blimp |
| Rnaset2b | 5.20E-85 | -0.325767 | 0.085 | 0.272 | 8.28E-81 | Cre_Blimp |
| Zufsp | 3.92E-46 | -0.32546 | 0.165 | 0.321 | 6.25E-42 | Cre_Blimp |
| Pml | 1.89E-54 | -0.324754 | 0.12 | 0.277 | 3.00E-50 | Cre_Blimp |
| Ddx60 | 9.00E-48 | -0.324464 | 0.065 | 0.187 | 1.43E-43 | Cre_Blimp |
| Lnpep | 5.97E-48 | -0.32439 | 0.394 | 0.578 | 9.51E-44 | Cre_Blimp |
| Gm4070 | 2.17E-60 | -0.323725 | 0.149 | 0.326 | 3.45E-56 | Cre_Blimp |
| Cflar | 1.14E-52 | -0.322999 | 0.29 | 0.479 | 1.81E-48 | Cre_Blimp |
| Chd7 | 2.49E-46 | -0.322748 | 0.267 | 0.441 | 3.96E-42 | Cre_Blimp |
| Lyst | 1.00E-40 | -0.322212 | 0.108 | 0.235 | 1.59E-36 | Cre_Blimp |
| Fam46a | 1.82E-31 | -0.320875 | 0.196 | 0.296 | 2.90E-27 | Cre_Bcl6 |
| Nlrc5 | 4.08E-51 | -0.320794 | 0.235 | 0.415 | 6.50E-47 | Cre_Blimp |
| Ccnl2 | 1.63E-52 | -0.319714 | 0.314 | 0.504 | 2.59E-48 | Cre_Blimp |
| Zfp36l1 | 1.45E-49 | -0.315597 | 0.483 | 0.624 | 2.30E-45 | Cre_Bcl6 |

| | | | | | | |
|---|---|---|---|---|---|---|
| Ube2l6 | 7.27E-50 | -0.314232 | 0.121 | 0.269 | 1.16E-45 | Cre_Blimp |
| Kbtbd11 | 6.85E-53 | -0.313727 | 0.166 | 0.335 | 1.09E-48 | Cre_Blimp |
| Skil | 3.76E-53 | -0.313159 | 0.185 | 0.36 | 5.99E-49 | Cre_Blimp |
| Kat6a | 8.22E-63 | -0.312779 | 0.134 | 0.309 | 1.31E-58 | Cre_Blimp |
| Ptprc | 5.28E-71 | -0.312558 | 0.872 | 0.966 | 8.41E-67 | Cre_Blimp |
| Rabgap1l | 4.90E-47 | -0.312335 | 0.401 | 0.54 | 7.80E-43 | Cre_Bcl6 |
| Oas3 | 4.88E-46 | -0.312292 | 0.127 | 0.272 | 7.78E-42 | Cre_Blimp |
| Dusp4 | 5.36E-33 | -0.311515 | 0.113 | 0.223 | 8.54E-29 | Cre_Blimp |
| Ahnak | 8.47E-35 | -0.311162 | 0.559 | 0.709 | 1.35E-30 | Cre_Blimp |
| Itk | 3.72E-64 | -0.310881 | 0.497 | 0.706 | 5.93E-60 | Cre_Blimp |
| Jak2 | 1.23E-60 | -0.309839 | 0.217 | 0.411 | 1.96E-56 | Cre_Blimp |
| Gpr55 | 1.93E-60 | -0.309129 | 0.035 | 0.144 | 3.08E-56 | Cre_Blimp |
| Irf8 | 4.66E-31 | -0.308974 | 0.093 | 0.197 | 7.42E-27 | Cre_Blimp |
| Entpd1 | 2.50E-56 | -0.308848 | 0.054 | 0.179 | 3.98E-52 | Cre_Blimp |
| Ets1 | 1.40E-57 | -0.308614 | 0.753 | 0.893 | 2.23E-53 | Cre_Blimp |
| Trim25 | 6.80E-56 | -0.307888 | 0.11 | 0.265 | 1.08E-51 | Cre_Blimp |
| Aebp2 | 1.30E-56 | -0.307622 | 0.282 | 0.478 | 2.07E-52 | Cre_Blimp |
| Plec | 1.78E-33 | -0.306528 | 0.226 | 0.366 | 2.83E-29 | Cre_Blimp |
| Tnfsf8 | 9.92E-21 | -0.306473 | 0.198 | 0.285 | 1.58E-16 | Cre_Bcl6 |
| Arhgef1 | 8.43E-72 | -0.306405 | 0.553 | 0.764 | 1.34E-67 | Cre_Blimp |
| Ptbp3 | 3.97E-72 | -0.306063 | 0.434 | 0.661 | 6.32E-68 | Cre_Blimp |
| Ifi204 | 3.03E-25 | -0.30605 | 0.108 | 0.197 | 4.82E-21 | Cre_Blimp |
| Tapbp | 5.18E-52 | -0.305941 | 0.52 | 0.705 | 8.25E-48 | Cre_Blimp |
| Cdk6 | 9.60E-28 | -0.305065 | 0.196 | 0.314 | 1.53E-23 | Cre_Blimp |
| Sesn3 | 3.29E-41 | -0.304154 | 0.307 | 0.429 | 5.23E-37 | Cre_Bcl6 |
| Irf1 | 9.02E-55 | -0.303792 | 0.381 | 0.579 | 1.44E-50 | Cre_Blimp |
| Pitpna | 7.60E-111 | -0.303658 | 0.342 | 0.624 | 1.21E-106 | Cre_Blimp |
| Abhd17a | 1.45E-91 | -0.302972 | 0.205 | 0.442 | 2.31E-87 | Cre_Blimp |
| Smg1 | 3.47E-46 | -0.302393 | 0.241 | 0.412 | 5.52E-42 | Cre_Blimp |
| Ly6c2 | 4.02E-11 | -0.300485 | 0.273 | 0.353 | 6.40E-07 | Cre_Blimp |
| Sp100 | 3.55E-66 | -0.300478 | 0.61 | 0.8 | 5.65E-62 | Cre_Blimp |
| Ttc39b | 1.60E-52 | -0.300313 | 0.374 | 0.568 | 2.54E-48 | Cre_Blimp |
| Birc6 | 1.00E-50 | -0.300313 | 0.357 | 0.547 | 1.60E-46 | Cre_Blimp |
| Rgcc | 2.10E-17 | 0.300236 | 0.263 | 0.204 | 3.34E-13 | Cre_Blimp |
| S100a11 | 8.55E-60 | 0.30996 | 0.842 | 0.806 | 1.36E-55 | Cre_Blimp |
| Lif | 3.61E-57 | 0.311533 | 0.129 | 0.026 | 5.76E-53 | Cre_Blimp |
| Pdlim1 | 2.80E-38 | 0.31213 | 0.349 | 0.248 | 4.46E-34 | Cre_Blimp |
| Uba52 | 4.35E-37 | 0.312711 | 0.899 | 0.797 | 6.92E-33 | Cre_Blimp |
| Prr13 | 1.65E-98 | 0.312956 | 0.74 | 0.743 | 2.63E-94 | Cre_Blimp |
| Btf3 | 4.81E-113 | 0.31549 | 0.899 | 0.862 | 7.67E-109 | Cre_Blimp |
| Serinc3 | 4.04E-44 | 0.316931 | 0.558 | 0.482 | 6.43E-40 | Cre_Blimp |
| Tuba1b | 5.49E-36 | 0.321631 | 0.359 | 0.305 | 8.74E-32 | Cre_Blimp |
| Igfbp7 | 4.48E-33 | 0.322812 | 0.134 | 0.06 | 7.14E-29 | Cre_Bcl6 |
| Naca | 5.32E-121 | 0.327846 | 0.926 | 0.9 | 8.47E-117 | Cre_Blimp |
| Eef1b2 | 1.62E-115 | 0.333551 | 0.906 | 0.862 | 2.59E-111 | Cre_Blimp |
| Cd74 | 1.27E-19 | 0.333742 | 0.151 | 0.085 | 2.02E-15 | Cre_Blimp |

| | | | | | | |
|---|---|---|---|---|---|---|
| Anxa1 | 1.78E-32 | 0.33638 | 0.259 | 0.164 | 2.83E-28 | Cre_Blimp |
| Sh3bgrl3 | 5.95E-147 | 0.34391 | 0.942 | 0.917 | 9.48E-143 | Cre_Bcl6 |
| Npm1 | 1.48E-102 | 0.351932 | 0.863 | 0.819 | 2.35E-98 | Cre_Blimp |
| Rack1 | 4.11E-136 | 0.361311 | 0.949 | 0.909 | 6.55E-132 | Cre_Blimp |
| Vim | 3.93E-79 | 0.366634 | 0.883 | 0.818 | 6.26E-75 | Cre_Bcl6 |
| Il7r | 4.46E-37 | 0.37218 | 0.631 | 0.525 | 7.11E-33 | Cre_Blimp |
| Id2 | 3.16E-86 | 0.372473 | 0.852 | 0.753 | 5.04E-82 | Cre_Bcl6 |
| Tmsb4x | 1.67E-158 | 0.374023 | 0.994 | 0.997 | 2.66E-154 | Cre_Bcl6 |
| Klrc1 | 3.30E-27 | 0.374369 | 0.325 | 0.23 | 5.26E-23 | Cre_Blimp |
| Ifitm3 | 4.81E-50 | 0.395509 | 0.492 | 0.483 | 7.67E-46 | Cre_Blimp |
| Ccr7 | 7.59E-48 | 0.401632 | 0.247 | 0.108 | 1.21E-43 | Cre_Blimp |
| Eif3e | 5.19E-125 | 0.410938 | 0.683 | 0.606 | 8.26E-121 | Cre_Blimp |
| Glrx | 1.09E-109 | 0.426996 | 0.707 | 0.505 | 1.74E-105 | Cre_Bcl6 |
| Capg | 9.73E-106 | 0.428433 | 0.743 | 0.612 | 1.55E-101 | Cre_Bcl6 |
| Prdx1 | 1.26E-127 | 0.428811 | 0.663 | 0.609 | 2.01E-123 | Cre_Blimp |
| Rbm3 | 1.36E-194 | 0.435732 | 0.891 | 0.871 | 2.17E-190 | Cre_Blimp |
| S100a10 | 9.11E-170 | 0.447268 | 0.932 | 0.88 | 1.45E-165 | Cre_Bcl6 |
| Lgals1 | 2.25E-134 | 0.44795 | 0.922 | 0.878 | 3.59E-130 | Cre_Bcl6 |
| Heph | 1.44E-64 | 0.45882 | 0.23 | 0.216 | 2.29E-60 | Cre_Blimp |
| Plac8 | 1.20E-41 | 0.473964 | 0.484 | 0.366 | 1.91E-37 | Cre_Bcl6 |
| Nkg7 | 6.25E-103 | 0.489662 | 0.786 | 0.654 | 9.96E-99 | Cre_Bcl6 |
| Crip1 | 1.39E-125 | 0.50063 | 0.898 | 0.829 | 2.21E-121 | Cre_Bcl6 |
| Hilpda | 4.85E-37 | 0.512029 | 0.412 | 0.306 | 7.72E-33 | Cre_Bcl6 |
| AC163354.1 | 1.21E-108 | 0.517569 | 0.355 | 0.145 | 1.93E-104 | Cre_Bcl6 |
| Lgals7 | 6.02E-42 | 0.524021 | 0.258 | 0.151 | 9.59E-38 | Cre_Bcl6 |
| Prdm1 | 3.64E-88 | 0.540195 | 0.288 | 0.105 | 5.79E-84 | Cre_Blimp |
| Nme2 | 8.92E-98 | 0.548089 | 0.603 | 0.42 | 1.42E-93 | Cre_Blimp |
| Cd7 | 7.43E-50 | 0.568141 | 0.347 | 0.226 | 1.18E-45 | Cre_Blimp |
| S100a6 | 2.04E-140 | 0.583964 | 0.847 | 0.736 | 3.25E-136 | Cre_Bcl6 |
| Ifitm2 | 1.99E-86 | 0.594211 | 0.6 | 0.427 | 3.17E-82 | Cre_Bcl6 |
| 1500009L16Rik | 1.51E-254 | 0.640874 | 0.375 | 0.069 | 2.40E-250 | Cre_Bcl6 |
| Lgals3 | 1.42E-96 | 0.664934 | 0.55 | 0.359 | 2.26E-92 | Cre_Bcl6 |
| Ifitm1 | 5.35E-47 | 0.698538 | 0.367 | 0.275 | 8.52E-43 | Cre_Blimp |
| Il13 | 3.43E-39 | 0.886446 | 0.104 | 0.031 | 5.46E-35 | Cre_Bcl6 |
| Csf2 | 5.97E-42 | 0.929523 | 0.161 | 0.069 | 9.50E-38 | Cre_Bcl6 |
