## Supplementary Figs 1-2 and Tables 6-8 for "A post-transcriptional regulatory checkpoint controls the response of tumor-infiltrating cytotoxic CD4^+^ T cells to immunotherapy"

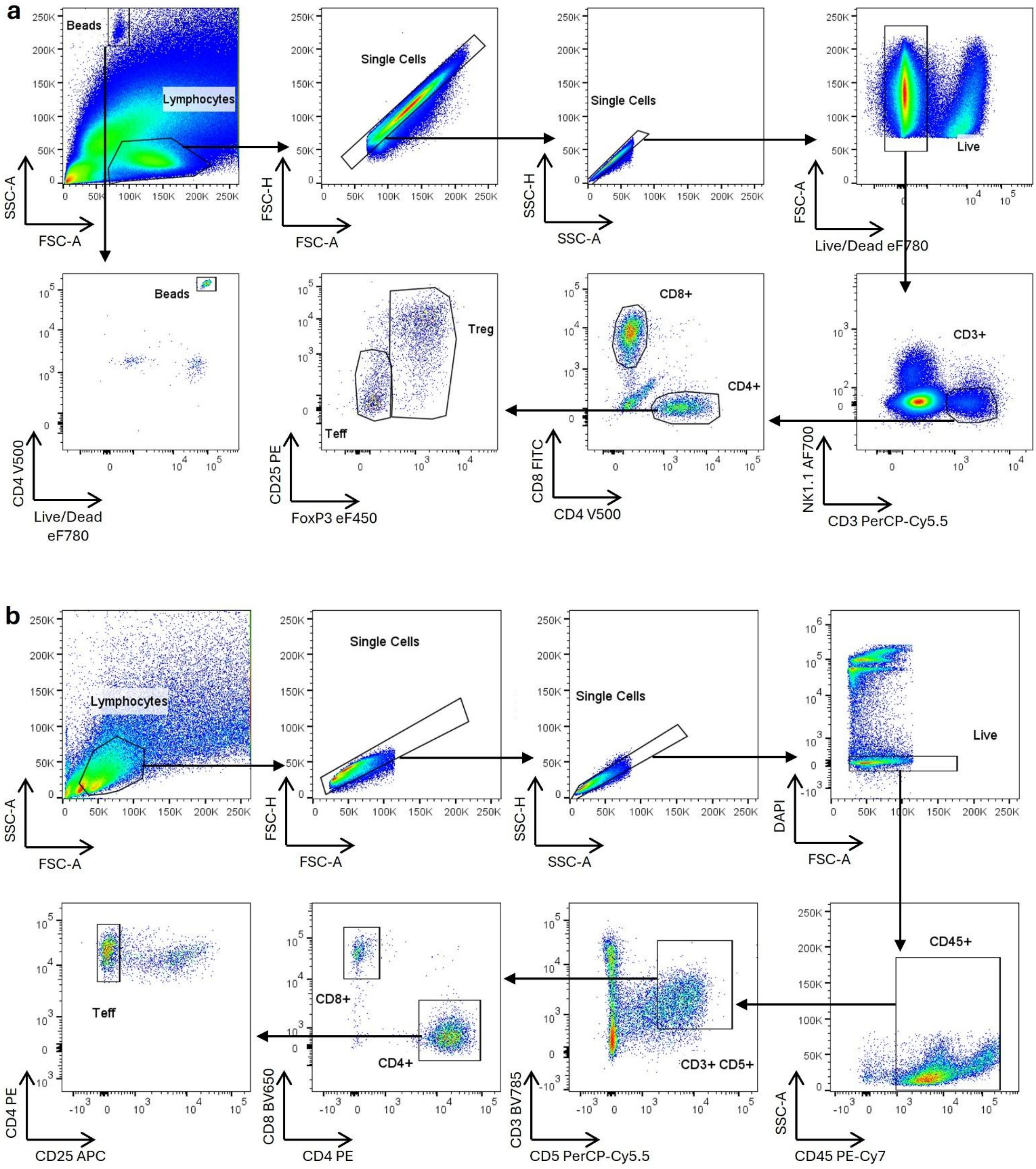


**Supplementary Fig. 1 Gating strategy for mouse samples.**

Gating strategies applied throughout manuscript. **a,** Gating strategy applied to mouse samples for flow cytometry analysis. **b,** Gating strategy applied to mouse samples for FACS for RNA analysis.

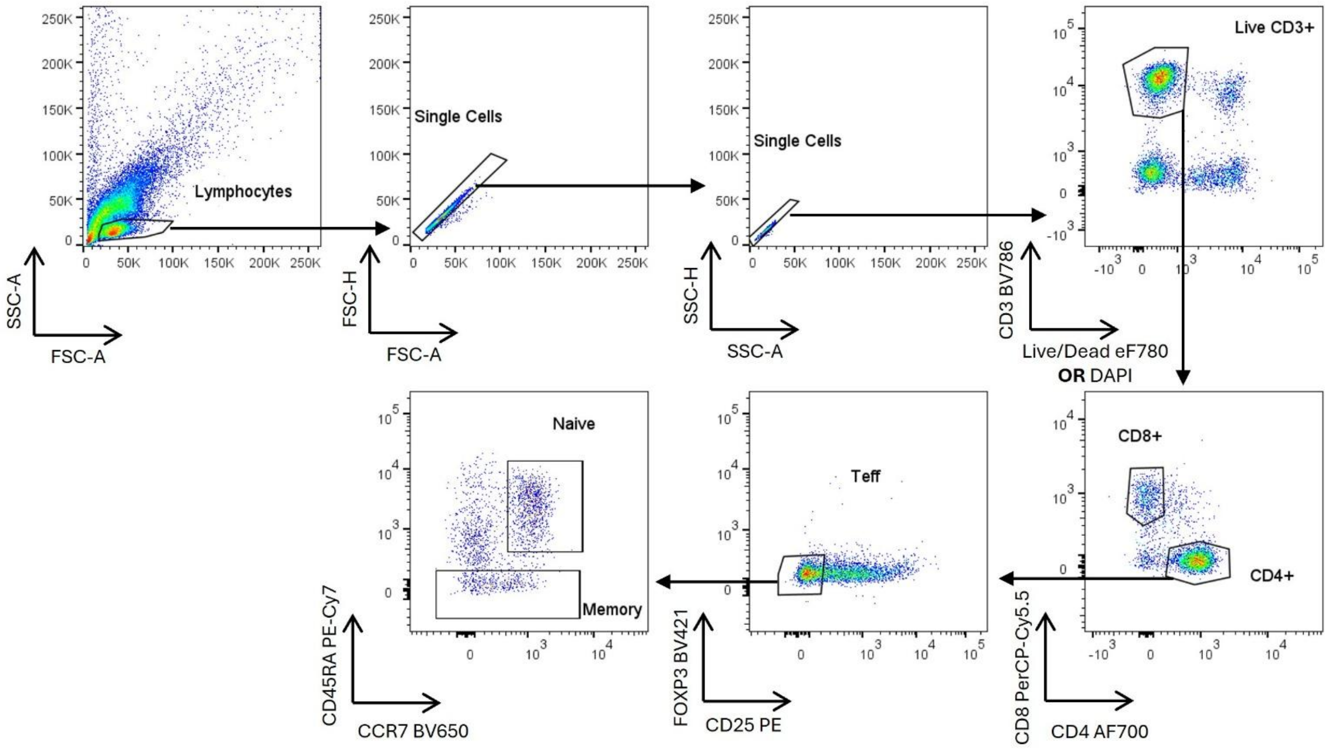


**Supplementary Fig. 2 Gating strategy for human PBMC and TIL samples.**
Gating strategy applied to human samples, applicable to Figure 1 and Extended Data Fig. 2. For flow cytometry analysis, live/dead stain used was fixable viability dye (eFluor 780, ThermoFisher). For FACS samples, DAPI was employed as live/dead stain. Remaining gating is applicable to both flow cytometry and FACS, with the exception of FOXP3, which was only used in flow cytometry analysis. In this case, $CD25^{-}$ Teff were gated against CD4.

| Supplementary Table 6 | Antibodies used for FACS and flow cytometry | |
|---|---|---|
| **Antibody** | **Source** | **Catalogue number/identifier** |
| **Anti-mouse antigen antibodies** | | |
| Anti-mouse FoxP3-eFluor450 (FJK-16s) | Thermo Fisher | Cat#48-5773; RRID: AB_1518812 |
| Anti-mouse CD8α-BV650 (53-6.7) | Thermo Fisher | Cat#100742; RRID:AB_2563056 |
| Anti-mouse CD8α-AF488 (53-6.7) | Biolegend | Cat#100726; RRID:AB_493423 |
| Anti-mouse/human Granzyme B monoclonal antibody (GB11), APC | Thermo Fisher | Cat#GRB05; RRID: AB_2536539 |
| Anti-mouse NK1.1-BUV395 (PK136) | BD Biosciences | Cat#564144; RRID: AB_2738618 |
| Anti-mouse CD4-BUV496 (GK1.5) | BD Biosciences | Cat#612952; RRID: AB_2813886 |
| Anti-mouse CD45-BUV563 (30-F11) | BD Biosciences | Cat#612924; RRID: AB_2722550 |
| Anti-mouse/human CD11b-BUV661 (M1/70) | BD Biosciences | Cat#612977; RRID: AB_2870249 |
| Anti-mouse CD3-BUV737 (17A2) | BD Biosciences | Cat#612803; RRID: AB_2738781 |
| Anti-mouse CD8α-BUV805 (53-6.7)<br>Anti-mouse CD3-BV785 (17A2) | BD Biosciences<br>BioLegend | Cat#564920; RRID: AB_2870186<br>Cat#100231; RRID:AB_11218805 |
| Anti-mouse CD3-PerCPCy5.5 (17A2) | BioLegend | Cat#100217; RRID:AB_1595597 |
| Anti-mouse CD4-V500 (RM4-5) | BD Biosciences | 560782; RRID:AB_1937315 |
| Anti-mouse CD4-PE (RM4-5) | BD Biosciences | Cat#553049; RRID:AB_394585 |
| Anti-mouse CD25-PE (PC61.5)<br>Anti-mouse CD25-eFluor660 (7D4) | Thermo Fisher<br>Thermo Fisher | Cat#12-0251; RRID:AB_465608<br>Cat#50-0252; RRID:AB_11149355 |
| Anti-mouse CD25-APC (PC61) | BioLegend | Cat# 102012; RRID: AB_312861 |
| Anti-mouse KLRG1-PE-Cy7 (2F1) | BioLegend | Cat#138416; RRID:AB_2561736 |
| Anti-mouse CD5 PerCP-Cy5.5 (53-7.3) | Thermo Fisher | Cat#45-0051; RRID:AB_914334 |
| Anti-mouse CD45-PE-Cy7 (30-F11) | BioLegend | Cat#103114; RRID:AB_312979 |
| Anti-mouse CD223 (LAG-3)–BV650 (C9B7W) | BioLegend | Cat#125227; RRID: AB_2687209 |
| Anti-mouse TNFa–PE-Cy7 (MP6-XT22) | BioLegend | Cat# 506323; RRID: AB_2204356 |
| Anti-mouse IFNg-AlexaFluor488 (XMG1.2) | BioLegend | Cat#505813; RRID: AB_493312 |
| Anti-mouse FoxP3-AF700 (FJK-16S) | eBioscience | Cat#56-5773-80; RRID: AB_1210557 |
| Anti-mouse NK1.1-AlexaFluor700 (PK136) | Thermo Fisher | Cat#56-5941; RRID: AB_2574505 |
| Anti-mouse NK1.1-BV650 (PK136) | BioLegend | Cat#108735; RRID: AB_11147949 |
| Anti-mouse/human CD11b-BV711 (M1/70) | BioLegend | Cat#101242; RRID: AB_2563310 |
| Anti-mouse CD279 (PD-1)-PE-Dazzle594 (29F.1A12) | BioLegend | Cat#135228; RRID: AB_2566006 |
| Anti-mouse CD223 (LAG-3)-PerCP-eF710 (C9B7W) | eBioscience | Cat#46-2231-82; RRID: AB_11151334 |
| Anti-mouse CD366 (TIM-3)-PE (8B.2C12) | eBioscience | Cat#12-5871-82; RRID: AB_465977 |
| Anti-mouse/human/rat CD278 (ICOS)-PE-Cy7 (C398.4A) | BioLegend | Cat#313520; RRID: AB_10643411 |
| Anti-mouse CD137 (4-1BB)-Biotin (17B5) | BioLegend | Cat#106104; RRID: AB_313241 |
| **Anti-human antigen antibodies** | | |
| Anti-human CD8-PerCP-Cy5.5 (SK1) | BioLegend | Cat#344709; RRID:AB_2044009 |
| Anti-human FoxP3-BV421 (206D) | BioLegend | Cat#320124; RRID:AB_2565972 |
| Anti-human CCR7-BV650 (G043H7) | BioLegend | Cat#353233; RRID:AB_2562041 |
| Anti-human CD3-BV786 (SK7) | BioLegend | Cat#344841; RRID:AB_2616890 |
| Anti-human CD4-AF700 (OKT4) | eBioscience | Cat#56-0048; RRID:AB_10718539 |
| Anti-human CD25-PE (24212) | R&D Systems | Cat# FAB1020P; RRID:AB_356980 |
| Anti-human CD45RA-PE-Cy7 (HI100) | BioLegend | Cat#304125; RRID:AB_10709440 |

| **Supplementary Table 7** | **Antibodies used for CyTOF** | |
|---|---|---|
| **Antibody** | **Source** | **Catalogue number/identifier** |
| **Antibodies used in CyTOF analysis** | | |
| Anti-mouse CD45 110-Cd (30-F11) | BioLegend | Cat#103102; Conjugated in-house |
| Anti-mouse CD3 106-Cd (17A2) | BioLegend | Cat#100202; Conjugated in-house |
| Anti-mouse CD4 153-Eu (RM4-5) | BioLegend | Cat#100506; Conjugated in-house |
| Anti-mouse CD8α 154-Sm (2.43) | BioXcell | Cat# BE0061; Conjugated in-house |
| Anti-mouse FoxP3 157-Gd (FJK-16S) | eBioscience | Cat#15247457; Conjugated in-house |
| Anti-pSTAT5[Y694] 144-Nd (47/Stat5) | BD Biosciences | Cat#611965; Conjugated in-house |
| Anti-pSTAT3[Y705] 169-Tm (4/P-Stat3) | BD Biosciences | Cat#612357; Conjugated in-house |

| **Supplementary Table 8** | **Primers used for RT-qPCR** | |
|---|---|---|
| **RT-qPCR primers** | **Forward** | **Reverse** |
| **Mouse** | | |
| *Gzmb* mRNA | CTGGGTCTTCTCCTGTTCTTTG | GACCCTACATGGCCTTACTTTC |
| *Prdm1* mRNA | GAAGGGAACACGCTTTGGAC | GATTCACGTAGCGCATCCAG |
| *Zfp36l1* mRNA | CTTCACGACACACCAGATCCTAGT | TGCTGTAGTTGAGCATCTTGTTACC |
| *GFPZFP36L1* mRNA | CGTGTCTGCCACCATCTT | CCCACTGCCTTTCTGTCC |
| *Hprt1* mRNA | TCAGTCAACGGGGGACATAAA | GGGGCTGTACTGCTTAACCAG |
| *Gzmb* pre-mRNA | ACTGCTGCTCACTGTGAAGG | AGCACCTTAGGATGAAGGCG |
| **Human** | | |
| *GZMB* mRNA | TGCGAATCTGACTTACGCCAT | GGAGGCATGCCATTGTTTCG |
| *ACTB* mRNA | AGCACAGAGCCTCGCCTTT | TCATCCATGGTGAGCTGGC |
| **Taqman assays** | **Assay** | |
| *Zfp36* | Mm00457144_m1(ThermoFisher) | |
| *Hprt* | Mm03024075_m1(ThermoFisher) | |
